# Cure-screening predicts mechanism of post-antibiotic relapse in Tuberculosis

**DOI:** 10.64898/2026.01.04.697520

**Authors:** Christian T. Michael, Maral Budak, Philana Ling Lin, Denise Kirschner

**Affiliations:** Kirschner lab, Department of Microbiology & Immunology, University of Michigan - Michigan Medicine, Ann Arbor, MI, USA; Department of Pediatrics, UPMC Children’s Hospital of Pittsburgh, University of Pittsburgh School of Medicine, Pittsburgh, PA, USA

**Keywords:** Multi-scale Modeling, Paucibacillary tuberculosis, Diagnostic sensitivity

## Abstract

Tuberculosis (TB) remains a global health concern, as *Mycobacterium tuberculosis* (Mtb) currently infects a quarter of the world’s population. Though many TB patients sterilize infection with short-course treatment, regimens shorter than 4 months risk post-treatment, potentially facilitating drug resistance. Granulomas are spatially-heterogeneous hallmark immune structures that form during TB, comprising immune cells and *caseum*, necrotic tissue that can trap Mtb in a non-replicating state. Two mechanisms causing relapse have been hypothesized: *persistence*, where treatment kills all replicating Mtb, and relapse follows once Mtb trapped within caseum returns to a replicative niche; and *threshold*, where replicating Mtb remain alive below detectable levels. Typically, clinical relapse is described as TB recurrence <2 years after a misdiagnosis of cure upon treatment completion (MDxC). Comparatively, many experimental models cannot screen for cure and examine relapse ∼2-months after treatment completion. Capacity to untangle these considerations *in vivo* are limited. Here, we examine the impact of study design (e.g., cure screening) on mechanisms underpinning reported instances of relapse using our computational model capturing whole-host Mtb infection dynamics, *HostSim*. Simulations uncover rates of reported relapse depending on whether hosts are screened for cure upon treatment completion. If not screened, then relapse is likely driven by incomplete sterilization of replicating bacteria; whereas cure-screened relapse is most likely to be caused by gradual expansion of non-replicating Mtb from within caseum into cellular areas of a granuloma. This suggests that TB patients that relapse after MDxC may best be treated with caseum-penetrating antibiotics such as rifamycin-class antibiotics.

**Importance:** Incomplete treatment of TB leads to risks relapse, which may occur years later. The threat of relapse is the main reason TB treatment takes 4-9 months. Predictors of relapse are not well-defined given variability in technical definitions of relapse and nuances of study designs. Here, we simulate both clinical and experimental relapse studies, including multiple diagnostic tests and relapse definitions, using biologically-based computation. We find that two hypothesized types of relapse that each potentially require a different treatment strategy are simultaneously at play. Simulations suggest that relapse after a “cure” diagnosis (most clinical studies) is caused by reactivation of non-replicating bacteria hidden from treatment within necrotic granuloma tissue or other sites, whereas experiments that cannot test for cure post-treatment are likely to report relapse caused by incomplete sterilization of replicating Mtb. Predictions depend on the currently unclear relationship between bacterial burden and clinical symptoms.

## 1. Introduction

Pulmonary tuberculosis (TB) remains a dire concern for countries across the globe and requires months of intensive treatment. A staggering one-quarter of the human population is infected with *Mycobacterium tuberculosis* (Mtb)^1^, with over 10,000 cases provisionally reported in the United States in 2025^2^. Approximately 90% of those infected individuals harbor clinically latent TB infection (LTBI), while the remainder progress to clinically active TB disease^3–6^. Accurate diagnosis and successful treatment of Mtb infection is required to eradicate TB disease worldwide, as incomplete treatment and poor compliance have led to a rise in drug-resistant Mtb strains^7^. The World Health Organization (WHO)- and Centers for Disease Control and Prevention (CDC)-recommended treatments for drug-susceptible, active TB are between 4-9 months using multiple drugs^8,9^, and the WHO-recommended treatment for latent Mtb infection takes 3-6 months^10^ often with many side effects. The WHO has identified a need for shorter treatment regimens to improve adherence and lower health costs^11^.

Many patients likely only need three months of treatment or less, though shortening the recommended treatment duration is dangerous as we cannot predict who will relapse^12^. Relapse (or recurrent TB disease consequent to effects of incomplete antibiotic treatment) poses a risk to patient health and promotes evolution of drug-resistant Mtb strains^13–15^. Standard treatment with HRZE—a first-line regimen of care for treating drug-susceptible TB, consisting of Isoniazid (H), Rifampicin (R), Pyrazinamide (Z), and Ethambutol (E)^9^—results in ∼5% relapse rate with six months of treatment or ∼a 20% rate for shorter-course treatment^16^. Still, that more than half of TB patients are successfully treated within 2-3 months^12^ presents a critical balancing act for public health agencies and clinicians alike as shortening treatment has the potential to reduce the risk of drug-induced side effects, financial and human public health resources.

Granulomas, the site of pulmonary Mtb infection and hallmark of human TB, pose a central challenge to studying relapse^a^. These complex lesions form during Mtb infection and they provide infection-site-specific pharmacokinetic and pharmacodynamic (PK/PD) barriers to treatment. They primarily form within lungs and mediastinal lymph nodes in humans^21–24^. Necrotic cores found in most granulomas (caseum) can harbor non-replicating Mtb, which may be a source of delayed relapse (see Appendix A1 for more on granulomas)^21,25^.

Though disease relapse is consequent to incomplete Mtb sterilization during treatment, the precise sequence of events within granulomas leading to clinical relapse is unknown. A recent review^12^ theorized two mutually-compatible mechanisms that may lead to relapse: persistence and threshold. *Persistence* suggests a reservoir of non-replicating Mtb exists after TB treatment that can later evolve the host into a subclinical state of TB^14^; and *threshold* implies there are a few replicating Mtb that are below a level of test detection (LOD) that remain after TB treatment or reside in a test-inaccessible compartment until treatment stops and subsequently, Mtb repopulate^b^. These mechanisms are challenging to separate both experimentally and clinically^12^ so it is currently not known what drives relapse.

There are few high-resolution relapse datasets as (i) studies aimed at shortening treatment in humans are designed to consider the increased associated relapse risk^12^, and (ii) animal studies face financial and translational challenges. Clinical trials are gold standard for analyzing treatment regimens and their relapse potential, but can take years to complete and cost an average of $40M^27^. These are often longer-course treatments designed to minimize relapse likelihood, thus relapses appear as rare events in human datasets by design. Multiple *in vivo* animal studies are employed to study TB outcomes^24^ including relapse—most notably including expensive nonhuman primates (NHPs)^28–30^ and a variety of relapse mouse models (RMMs)^31–34^, each with advantages and disadvantages.

It is not currently known how differences in relapse study designs between human and animal studies impact translatability of conclusions. To our knowledge, studies addressing relapse assume (often by necessity) that relapse is principally driven by a uniform set of mechanisms across all study designs. Differences between human and animal studies may be in part be due to either study design, how relapse is measured, or underlying physiological differences. Currently, precise relapse measurements are not directly compared between humans and animal studies. We bridge this gap by using a whole-host quantitative systems pharmacology model for treatment of TB called *HostSim*.

### 1.1 TB relapse in humans

We summarize the clinical presentation of active TB and relapse to clarify terminology. Clinically, active TB is associated with signs (e.g., cavitary lesions on chest x-rays, fever, etc.) and symptoms (cough, fatigue) of TB, usually with microbiologic confirmation of Mtb infection (sputum). During treatment, signs and symptoms often improve (improved chest x-ray, sputum smear and culture are Mtb-negative, resolution of cough and weight loss). Subclinical TB is another form of TB disease that is clinically asymptomatic but has diagnostic markers associated with active TB, such as Mtb present in sputum or bronchoalveolar lavage fluid (BALF), unlike most LTBI cases^4^. Clinically, relapse refers to having new signs and symptoms of TB after resolution by the end of treatment (Figure 1) and after exogenous Mtb re-infection has been ruled out^12^ (though in practice, ruling out reinfection is challenging^35^).

**Figure 1:**
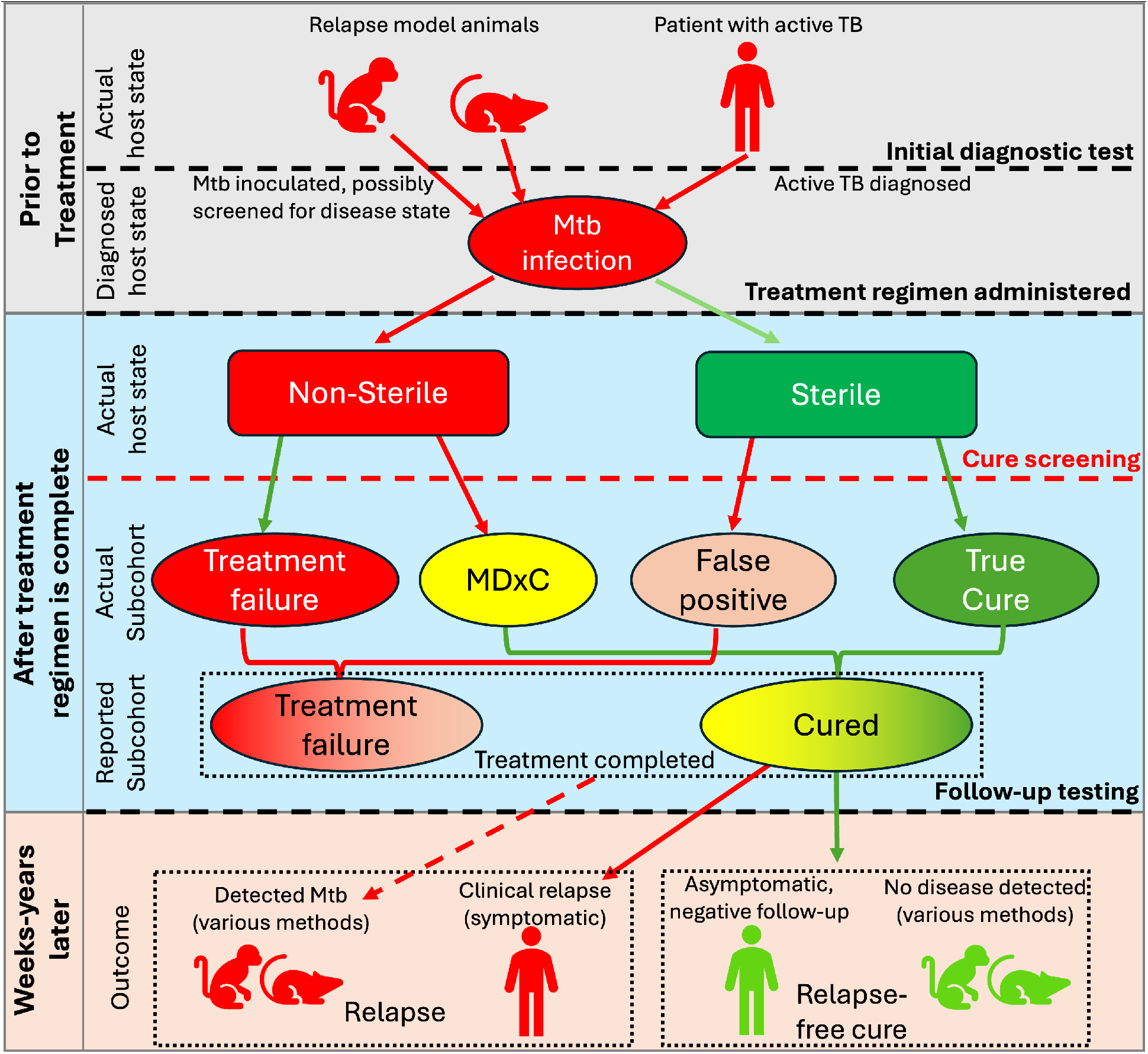
Flowchart of TB relapse assessment. We show how misdiagnosed cure (MDxC) and cure screening affect the assessment of post-treatment TB relapse. Note that some animals like RMMs cannot be resampled, and thus relapse is considered subsequent to treatment completion rather after MDxC (i.e., cure-screened relapse). Additionally, reported disease state may be impacted by interchanging tests used for diagnosis, cure-screening, and follow-up (e.g., follow-up being based on symptoms or bacteriological results).

Clinical relapse is typically defined as occurring after a patient has been misdiagnosed as cured (MDxC)^c^. Occasionally, relapse is grouped with treatment failure under the primary endpoints of “unfavorable outcomes” in studies with few reported instances of TB relapse^36,37^. The rate of relapse is typically very low in trials (<5% in most cases), as clinical trials prioritize patient safety and may be cancelled if the risk of relapse is unacceptably high^12,38^. It is impossible to eliminate MDxC cases, as to date there are no studies that can prove sterilization from all sites of infection. That is, there is no test that can determine whether a patient is completely treated.

While the goal of treatment is always sterilization of Mtb from all sites (lungs, lymph nodes, extrapulmonary sites may have varying burdens), resolution of symptoms and/or negative sputum cultures are typically used as a surrogate for assessing true sterilization. Patients treated for TB include those who are considered “TB-cured” (e.g., sputum culture negative after treatment is completed) and “TB-treated” (meaning they complete treatment but did not necessarily confirm that sputum was Mtb-negative). The additional step of cure-screening with a bacteriological confirmation of cure is not always performed, and it is common to conflate “TB-cured” with “TB treated” as equivalent stand-ins for sterilization. However, as relapse clinically follows MDxC, it may be that the most influential mechanisms driving relapse (and optimal re-treatment) are different between “TB-cured” and “TB-treated” patients; or may differ by clinical and microbiologic measures used to assess patients for cure. For this reason, we delineate between “cure-screened relapse” to indicate relapse after a Mtb-infected host has relapse subsequent to a microbiologic test for Mtb negativity, and “unscreened relapse” as those relapse that follows from treatment completion without a microbiologic test at the time of treatment completion.

Relapse assessment is not uniformly performed, varying based on test availability, operational definition, and consistency of medical care. Relapse is typically reported using a composite of one or more measures including positive bacterial smears or cultures, rapid-test molecular analysis, radiographic exams, and/or clinical TB symptom presentation^12,29,36,37,39^. Confounding things, each test may potentially identify different patients as having relapsed if tests used have different levels of detection (LODs). Some tests are commonly used to determine cure after treatment (e.g., two consecutive Mtb-negative sputum smears^40^) though no tests have been validated to predict relapse. Though specific tests can vary, some form of cure-screening test is performed in most human studies that we re-examine here, except where noted. We find that many studies report relapse only as a recurrence of symptomatic active TB disease, possibly with bacteriological follow-up^16,41–44^, though detection of non-symptomatic infection after treatment is sometimes reported or discussed as relapse^37^. Appendix Table A1 presents a list of diagnostic tests used to assess TB diagnosis, cure, and relapse (based in part on ^12^).

Although relapse can often be based on symptom presentation, the connection between patient symptoms, diagnostic inflammatory markers^d^, and lung bacterial burden remains murky in both human and animal-model TB. Even if one knew the precise number of Mtb and their metabolic state at every timepoint within lungs, we may not know whether culturable or symptomatic relapse will result. Patients with subclinical TB remain asymptomatic while shedding viable Mtb, which was historically considered a hallmark feature of clinical TB^4,45^. (This phenomenon is observed in NHPs, referred to as percolators, and comprises ∼5% of macaques^46–48^.) Previous studies found 10-25% of LTBI patients exhibited neutrophilic whole-blood transcriptomic signatures consistent with active TB^4,49^. In tandem with this, we observed in simulated virtual patients that having levels of inflammation markers such as interferon gamma (IFN-γ) do not predict host outcomes^50^. Conversely, clinical studies have shown multiple inflammatory markers present in patients with LTBI^51^. Together, this suggests that not all asymptomatic cases are equal^4^, with some engaging in an ongoing immune battles likely to contain metabolically-active Mtb from escaping granulomas, while other cases involve surveillance over non-replicating, caseum-trapped Mtb. This creates a spectrum of inflammatory signatures and symptoms that suggest more than one modality of relapse is possible.

### 1.2 Animal models of TB relapse

Both RMMs and NHPs, the most common animal relapse models, provide detailed relapse datasets but bear some substantial differences from each other and from human relapse studies. RMMs can be readily treated with human-equivalent doses (i.e., corresponding average plasma concentrations^52,53^) of first and second-line antibiotics. However, most mice do not form necrotic granulomas with the exception of C3HeB/FeJ mice^34,54,55^, and mice have disease states incongruent from those in humans (e.g., mice experience chronic Mtb infection). NHPs most closely represent human TB, but require expensive resources and expertise and sample sizes can be small^56^. Animal models have been given (intentionally) weak antibiotic regimens to allow for reliable and experimentally-controllable relapse^29^, though this experimental formulation of relapse may capture a disease state with large (if potentially asymptomatic) bacterial burdens at treatment completion.

In many cases, it is necessary to use proxy measures in lieu of host state when studying relapse in animals, and it is unclear whether these proxies over/underrepresent different mechanisms driving relapse. Assessing cure in animal models is not always comparable to diagnostic tests used in humans (e.g., repeated sampling of CFU in RMMs is not consistently possible^57^ or species-specific disease presentation^58^), thus individual animals are often not screened for cure when studying relapse (Figure 1) and use alternative measures. For instance, CFU enumeration in mouse lung homogenates or NHP granulomas is used as one proxy for clinical relapse (Appendix Table A1). It should be noted that some of these tests are likely more accurate than measures used in humans— e.g., tissue homogenates can capture non-replicating Mtb, though it is unclear whether human sputum CFU measures capture the same outcomes. Of course, animals cannot be followed post necropsy.

### 1.3 Assessing translatability of animal and clinical relapse using *HostSim*

If both persistence- and threshold-driven relapse can occur *in vivo* as hypothesized^12^, it is unclear whether both differ in terms of optimal treatment, or whether they are uniformly captured in different study designs. While each measurement and proxy has advantages and disadvantages (^12^ and Appendix Table A1), none can categorically predict relapse. For example, it was found in NHPs that measuring IFN-γ in BALF was insufficient for assessing effective antibiotic treatment efficacy^59^. Sputum culture conversion at eight-weeks is a common endpoint for clinical trials, but is not necessarily a good indicator of sterilizing activity^60–62^.

To understand how relapse measures and study designs impact translatability and conclusions of relapse studies, we recreate multiple *in vivo* relapse studies using a computational model of relapse. For this, we extend our virtual host treatment model of pulmonary TB, *HostSim*, to capture mechanisms that may lead to either persistence- or threshold-driven relapse, as well as perform virtual diagnostic tests that mimic *in vivo* diagnostic tests. *HostSim* bridges the translational gap between animal and human relapse studies by reproducing both clinical and experimental relapse studies and comparing conclusions. Our simulations suggest relapse subsequent to cure screening is less frequent and primarily *persistence-driven*—i.e., driven by a slow, continuous release of non-replicating bacteria derived from caseum. This can take a longer time before Mtb levels exceed LOD, potentially explaining the fact that some humans take up to 2 years to relapse^12,63,64^. On the other hand, simulating relapse without cure-screening upon treatment completion more often leaves a reservoir of replicating bacteria, leading to rapid *threshold-driven* relapse. Though this trend is true regardless of LOD, the relative frequency of these relapse types depends on the LOD of diagnostic tests used. Additionally, we derived estimates of LOD ranges in our simulations due to a current lack of understanding of how lung CFU relates to sputum/BALF CFU.

## 2. Results

### 2.1 Simulating misdiagnosed cure of antibiotic TB treatment (MDxC)

To simulate relapse, we study treatment of virtual hosts using *HostSim* to capture a plausible spectrum of infection and treatment outcomes observed in both humans and animals. This includes capturing Mtb trajectories before and during treatment, relative rates of reported MDxC, and subsequent Mtb recurrence.

#### 2.1.1 - HostSim simulates Mtb-immune interactions within heterogeneous virtual hosts

To mechanistically simulate host outcomes during Mtb infection and treatment we use *HostSim*, our recently-published, calibrated and validated whole-host pulmonary TB model^61,65,66^. Briefly, *HostSim* is a hybrid agent-based and ordinary differential equation model that represents interactions between host-immune cell populations in lungs, uninfected lung-draining lymph nodes, and a blood compartment. We can also track different Mtb subpopulations within multiple lung granulomas of a virtual host (within caseum, macrophages, or tissue). *HostSim* allows us to simulate temporal trajectories of various T cell and macrophage population sizes, as well as subpopulations of replicating and non-replicating Mtb within each granuloma for each host (see Methods) and their total counts within lungs. Recently, we calibrated *HostSim* to compare multiple measurements of drug efficacy of various first- and second-line antibiotic regimens by predicting the time-course trajectory of Mtb over time during treatment of drug-susceptible infection^61^.

We simulate a heterogeneous set of virtual patients (our “virtual cohort”) that we treat with each treatment and relapse scenarios. For all analyses, we generate a single virtual cohort of 500 Mtb-infected hosts, each patient inoculated with 13 bacteria leading to 13 granulomas at the start at *time* = 0^61^. We allow each virtual patient to develop naturally a mature Mtb infection until 300 days p.i. then take a snapshot of each granuloma simulation. From those snapshots, we can use our virtual cohort to study multiple *what-if* scenarios where we treat an entire cohort with one antibiotic regimen, halt treatment, then observe regimen-specific host and cohort outcomes both short- and long-term.

*HostSim* simulations capture a spectrum of host outcomes before, during, and after treatment. In Figure 2, we show dynamics of simulated Mtb bacterial trajectories over time during and after treatment simulating 8 weeks of HRZE. To simulate relapse, we first capture the spectrum of treatment outcomes: cure, (both MDxC and true sterilization), and treatment failure. To assess for cure, we implement virtual diagnostic tests by mimicking those performed *in vivo*, typically by comparing replicating or total Mtb counts against an established LOD (Methods).

**Figure 2:**
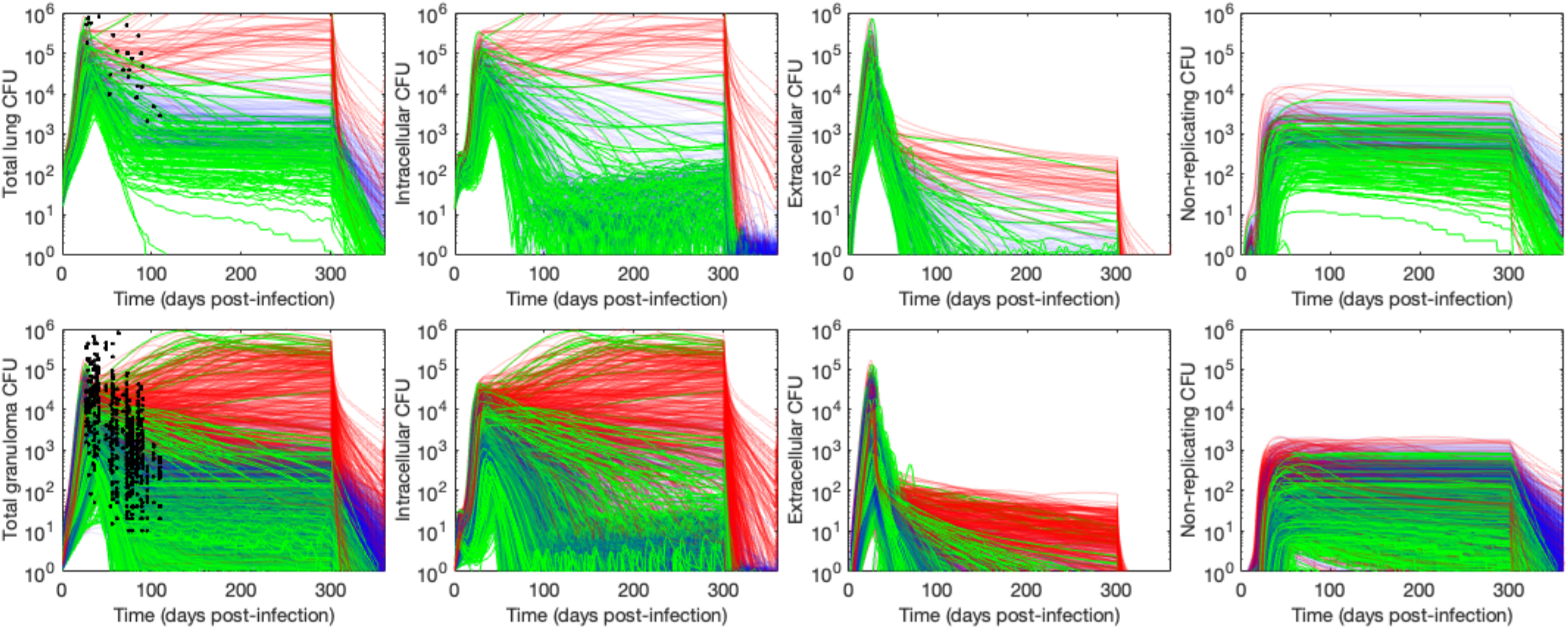
Infection trajectory of virtual hosts and granulomas before and after HRZE treatment. Each panel shows Mtb populations within *n* = 500 hosts (top row) and individual granulomas (13 for each patient, bottom row). All hosts are infected with Mtb at time 0 and, for visibility, we only show CFU from primary granulomas. Treatment begins at day 300 p.i. and ends at day 360 (i.e., after 2 months of treatment). Bacterial trajectories of virtual hosts with active disease prior to treatment (Methods) are shown in red. Bacterial trajectories of hosts that sterilized with 0 CFU (**true cure**) by the end of treatment are shown in green, and all other virtual hosts are shown in blue. As in our previous work ^61,65^, we compare CFU prior to treatment with analogous published CFU levels in NHP lungs and granulomas (black dots)^67–70^.

#### 2.1.2 - HostSim captures misdiagnosed cure rates reflective of regimen-specific relapse rates

To validate that our model captures MDxC, we simulate multiple short-course (8-week) antibiotic regimens and ensure that the relative MDxC rates reflect relative relapse rates of the well-studied regimens HRZE, RMZE, BPaL, and BPaLM^12,32,34,71–73^. We simulate MDxC by subtracting the proportion of virtual hosts or granulomas that truly sterilize (CFU = 0) from those with CFU below a set LOD; this difference isolates hosts that would likely be misdiagnosed as cured, or granulomas that would be assessed as sterile. Although actual relapse studies use a variety of diagnostic tests, we compare true virtual host sterilization against *Virtual Mtb plate* results, which mimic highly accurate experimental lung and granuloma homogenate cultures capable of detecting non-replicating bacteria (see Methods and Table S1). Specifically, we assume that whole-host homogenates have a LOD of 50 CFU, and granuloma-scale LOD is 10 CFU (Methods and ^61,74^). To determine whether pre-treatment CFU has an impact on MDxC rates, we delineate between hosts and granulomas that are high/low CFU prior to treatment, as in our previous work^61^. For this, we say that a *host* that is initially-high-CFU (>total 10,000 CFU pre-treatment) and initially-low-CFU (<total 10,000 CFU pre-treatment), and a *granuloma* that is initially-high-CFU (>1,000 CFU pre-treatment) and initially-low-CFU (<1,000 CFU pre-treatment). It should be noted that high CFU hosts may also have some low CFU granulomas but must have at least one high-CFU granuloma.

Our simulations show regimen-dependent MDxC rates that reflect relative risk of relapse in other studies. Our initial observations are not surprising in that most short-course mono-treatments for initially-low-CFU hosts results in large rates of MDxC and have little impact on initially-high-CFU hosts. Notably, HRZE has a high MDxC rate, particularly in high-CFU patients (42% sterile vs 76% cure), when compared with regimens including Moxifloxacin or Bedaquiline (e.g., BPaL with 74% sterile, 80% cured and RMZE with 80% sterile and 96% cured). This suggests that short-course treatment of high CFU-burden patients with only 8 weeks of HRZE will have a higher risk of MDxC, consistent with both (i) the high (∼40%) relapse rates found for short-course HRZE treatment in a 2016-2018 clinical study^12,71–73^; and with (ii) short-course mouse studies directly showing higher relapse rates in HRZ and HRZE than in BPaL, BPaLM, or RMZE ^32,34^.

##### Regardless of regimen, both the scale we measure and virtual pre-treatment CFU levels influence MDxC levels

Rates of MDxC are increased when observing initially-high-CFU hosts, when compared to initially-low-CFU hosts, particularly after multi-drug regimens (Figure 3). This positive-control case serves as model validation, reaffirming that hosts with severe infection are those most likely to experience cure-screened relapse. This pattern holds at both the granuloma and whole-host scales. Finally, we observe that large granuloma-scale MDxC rates amplify into even larger rates at the host scale, as previously predicted^75^.

**Figure 3:**
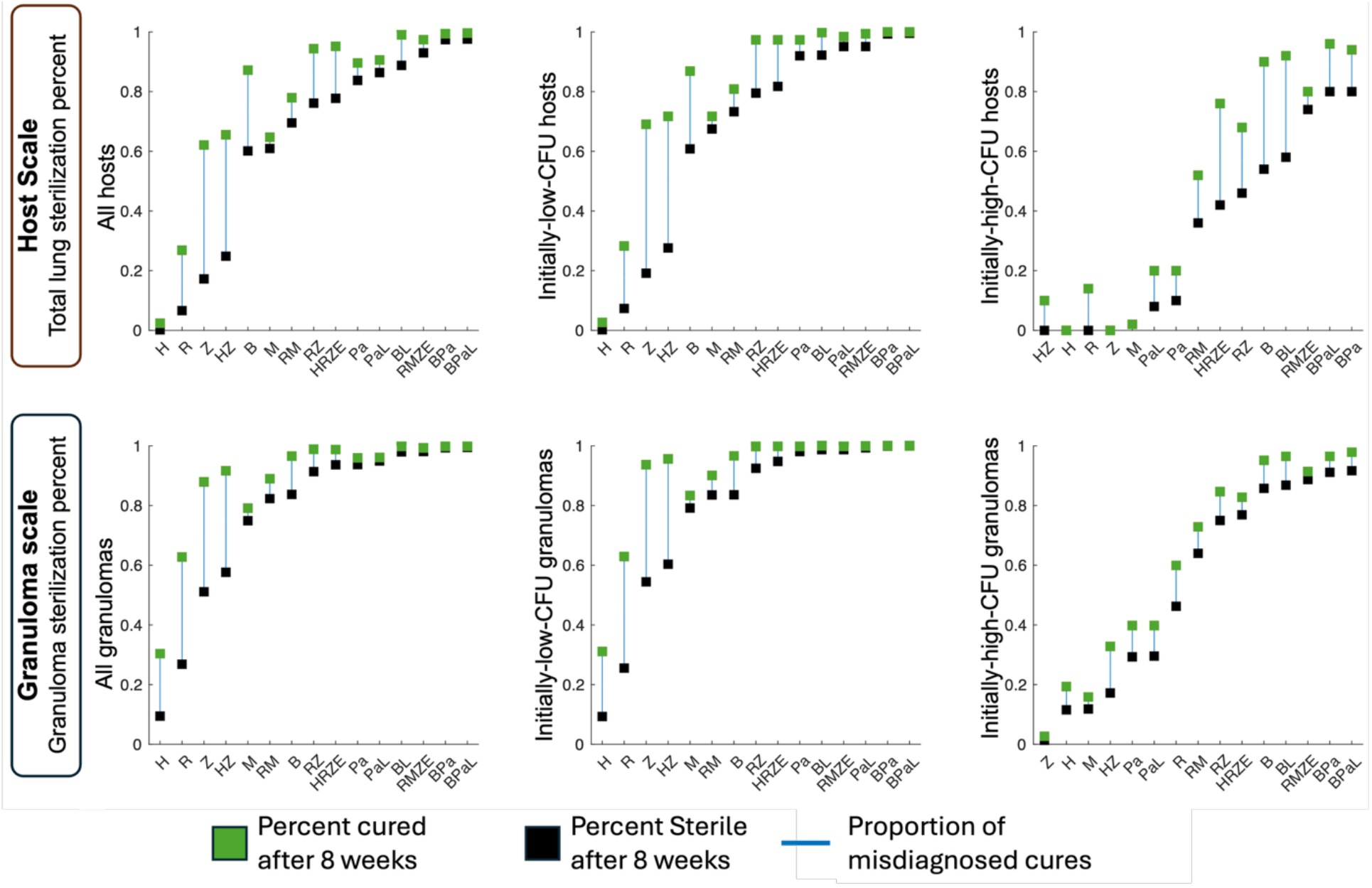
Misdiagnosed cure (MDxC) rates are specific to spatial scale and antibiotic regimen. We show cure and sterilization fractions of virtual hosts and granulomas after short-course antibiotic regimens. We administer a drug regimen for 8 weeks (regimens group (i), see Methods) to each of 500 virtual hosts in a virtual cohort (shown in top row) and a total of 6500 primary granulomas (shown in bottom row). For each regimen, the percentages for sterilization (black boxes) and lack of detectable infection (green boxes) after 8 weeks of treatment are shown. The left column shows treatment results for each of these virtual hosts and granulomas, whereas the center (right) columns show results for only hosts and granulomas that were low-CFU (high-CFU) pre-treatment. We notice that higher-CFU granulomas and hosts are more liable to have subclinical infection (i.e., non-sterile but with CFU < 10 (granuloma) or 50 (host)).

#### 2.1.3 - Most cure misdiagnosis stems from incomplete sterilization of caseum

In our studies of virtual hosts with TB treated for 2 months that experience MDxC after short-course treatment (Figure 3), we observe that viable Mtb are found within replicative niches only in those virtual hosts/granulomas that had high-CFU prior to treatment, and this was regardless of regimen used. After HRZE treatment, 7/500 virtual hosts have MDxC, and 6 of those 7 are initially-high-CFU. Within the MDxC initially-low-CFU host, nearly all CFU levels were caseum-bound and non-replicating. One of the six initially-high-CFU hosts with MDxC held Mtb intracellularly (inside macrophages), while the others held Mtb within caseum. Pooling HRZE-treated granulomas from all hosts, we observe thirteen virtual granulomas had CFU below LOD—two initially-low-CFU and eleven initially-high-CFU. In both initially-low-CFU granulomas with MDxC, all Mtb are trapped within caseum. Of the eleven high-CFU granulomas, eight harbor Mtb solely within caseum while the other three retain Mtb within macrophages at treatment completion. We have the ability to treat the same cohort with different regimens to compare outcomes and after treating the same virtual cohort with a different second-line regimen, BPaL^32,76^, only one initially-high-CFU host hosts replicating CFU below LOD at treatment completion, with all bacteria held entirely within caseum of an initially-low-CFU granuloma at treatment completion. Similar trends followed for other simulated regimens.

Overall, our simulations suggest that relapse studies will likely differentially capture relapse processes—either persistence- and threshold-driven relapse—that are based on CFU levels of each host prior to treatment. Using *HostSim* to identify MDxC after short-course treatment (and assuming that the diagnostic test can detect Mtb within caseum), we predict that hosts with low CFU levels prior to treatment likely harbor Mtb trapped within caseum; and, conversely, we predict MDxC hosts harboring replicating Mtb likely had high CFU levels prior to treatment. This suggests that (i) if clinical relapse occurs in initially-low-CFU hosts, it will likely be persistence-driven; and (ii) if a host experiences threshold-driven relapse, it likely had high CFU levels prior to treatment.

### 2.2 Validating a relapse model by recreating animal relapse studies

With the ability to capture a variety of host states that lead to misdiagnosed cure, we extend and calibrate *HostSim* during treatment to capture recurrent TB well beyond the time of treatment completion, i.e. to identify if and when relapse occurs. Given that we capture MDxC both within replicative and non-replicative niches, we represent mechanisms that can allow for either threshold- or persistence-driven relapse. For persistence relapse, we allow virtual non-replicating Mtb to be stochastically transported out of caseum over time into the cellular region of the granuloma (a role hypothesized to be played by neutrophils^77^), allowing a possibility for caseum-trapped Mtb to become replicative and lead to relapse and/or initiate nascent granuloma foundation^77^. Given the efficacy of clinically-administered regimens, there are sparse relapse data to calibrate the mechanism for stochastic release of Mtb. However, we assume this mechanism is also responsible for reactivation of LTBI, and so we calibrate rates of caseum-transport by recapitulating reactivation of LTBI during anti-TNF treatment such as etanercept or infliximab^78,79^ (Methods). The threshold relapse mechanism is implicitly captured by our model, as Mtb replication is always allowed during or after treatment, and this remains true for numbers of live intracellular or tissue Mtb at the end of treatment (if it is a MDxC).

We define virtual relapse per-study-design and classify that event as either persistence- or threshold-driven. To prepare each virtual host for relapse assessment, we (1) treat the virtual cohort with different antibiotic regimens, (2) if specified by the study design, screen hosts for cure upon treatment completion, and (3) simulate their infection progression without-treatment for a follow-up period of time. At some future time point post-treatment-completion, we run a follow-up diagnostic test on each host to determine relapse.

We classify simulations that result in relapse as threshold-driven if there is at least 1 replicating Mtb at treatment completion, and persistence-driven otherwise (any remaining Mtb are trapped within caseum).

#### 2.2.1 - Recreating a radiographic study of relapse in primates

To validate our relapse model, we replicate the design of a recent relapse study performed in NHPs that employed simian immunodeficiency virus (SIV)^29^. There, NHPs were treated with short-course antibiotic regimens for active-TB only monkeys (8 weeks of isoniazid and rifampicin, i.e. HR). After a one-month rest, they were then infected with SIV to induce relapse^5^. Relapse was primarily measured via observation of newly seeded granulomas using radiolabeled PET/CT scans; the study reported that 8/12 NHPs relapsed. To replicate this experiment using *HostSim*, we represent HIV-1 co-infection as initiating a linear decline of CD4^+^ T cells that we calibrated to T-cell blood concentrations reported from the 1990s^80,81^ (and measured prior to any treatment studies, see Appendix A3).

Standard relapse studies only admit humans (or NHPs) with specific pre-treatment disease states (e.g., active TB), so we similarly predict virtual cohort disease states and use them to allow for direct comparison to this study. We classify virtual hosts as having active TB if they have continued active bacterial growth, which we characterize by a species-specific fold-increase in CFU between treatment completion and follow-up (Methods). To ensure results are robust to different characterizations of active TB in our virtual patients, we perform a separate analysis considering the case where virtual hosts maintain high-CFU granulomas (CFU>1000 for over a month) as having active TB.

After recreating the NHP study, our simulations well-reproduce the NHP relapse rates^29^. Virtual disease state classification allows us to examine relapse rates within five sub-cohorts of our original *n* = 500 virtual hosts (Table 1). As in the original study, we simulate hosts with 2 months of HR treatment and assess MDxC upon treatment-completion using a *Virtual Clinic score*, a virtual diagnostic test we use as a proxy for clinical evaluation based on appearance of new granulomas and increasing Mtb counts (Methods). After 4 weeks of drug rest, we simulate virtual HIV infection for a further eight weeks. At that follow-up time, we assess each virtual host in the full cohort for relapse using a *Virtual PET/CT* method (see Methods) to mimic PET/CT-based tests used to determine relapse in the NHP study^29^ (Table 1). Using both of our NHP-like active-TB virtual relapse sub-cohorts result in ∼50% relapse, close to the 8/12 NHPs from their study data^29^.

**Table 1:** Predicted radiographic relapse rates by sub-cohort. Bolded cells indicate simulated results that are the most directly comparable to those from a recent NHP relapse experiment^**29**^. Columns 1-4 of each row describe the sub-cohort, frequency of relapse, and whether these relapses are persistence-driven (P) or threshold-driven (T). Columns 5-6 perform repeat analysis on the same sub-cohort but describe only cure-screened relapse. † Note that this this may include treatment of virtual hosts with subclinical or LTBI and may bear closer resemblance to reactivation than clinical relapse.

| Sub-cohort disease state classification | Does {CFU > 1000} also mean active? | # relapse out of # treated [% relapsed at 8 weeks] | Proportion of P vs T relapse (no screening) %P \| %T | # cure-screened relapse out of # cured in SC [% relapsed at 8 weeks] | Proportion of P vs T relapse (cure-screened) %P \| %T |
| --- | --- | --- | --- | --- | --- |
| Active (NHP-like) | ✗ | **50/100 [50%]** | 16% P \| 84% T | 20/68 [29%] | 40% P \| 60% T |
| Active (NHP-like) | ✓ | **56/109 [51%]** | 20% P \| 80% T | 24/75 [32%] | 46% P \| 54% T |
| Active (Human-like) | ✗ | 42/47 [89%] | 5% P \| 95% T | 12/15 [80%] | 17% P \| 83% T |
| Active (Human-like) | ✓ | 52/63 [82%] | 13% P \| 87% T | 20/29 [69%] | 35% P \| 65% T |
| All† (Active + subclinical) | N/A | 82/483 [17%] | 34% P \| 66% T | 49/447 [11%] | 56% P \| 44% T |

We find that the majority of cure-screened relapse appears as persistence-driven relapse, which mostly exhibit slowly-increasing levels of intracellular bacteria (Table 1). Relapse frequency per-sub-cohort decreases by 6-20% with cure screening. Simulated cure-screened relapse rates tends to be slower, persistence-driven (Figure 4), and feature a low (stochastically-oscillating) level of Mtb as bacteria move between niches too fine-grained for experimental detection ^50,82^. More than half of simulated relapses subsequent to treatment failure (i.e., those that would have been excluded via cure-screening) results from incomplete sterilization of macrophages (Figure 4, red curves), indicating threshold-driven relapse. Some of these relapses result in rapid bacterial regrowth in 1-2-week periods that restore CFU to pre-treatment levels. Others exhibit either slow or no regrowth of Mtb populations.

**Figure 4:**
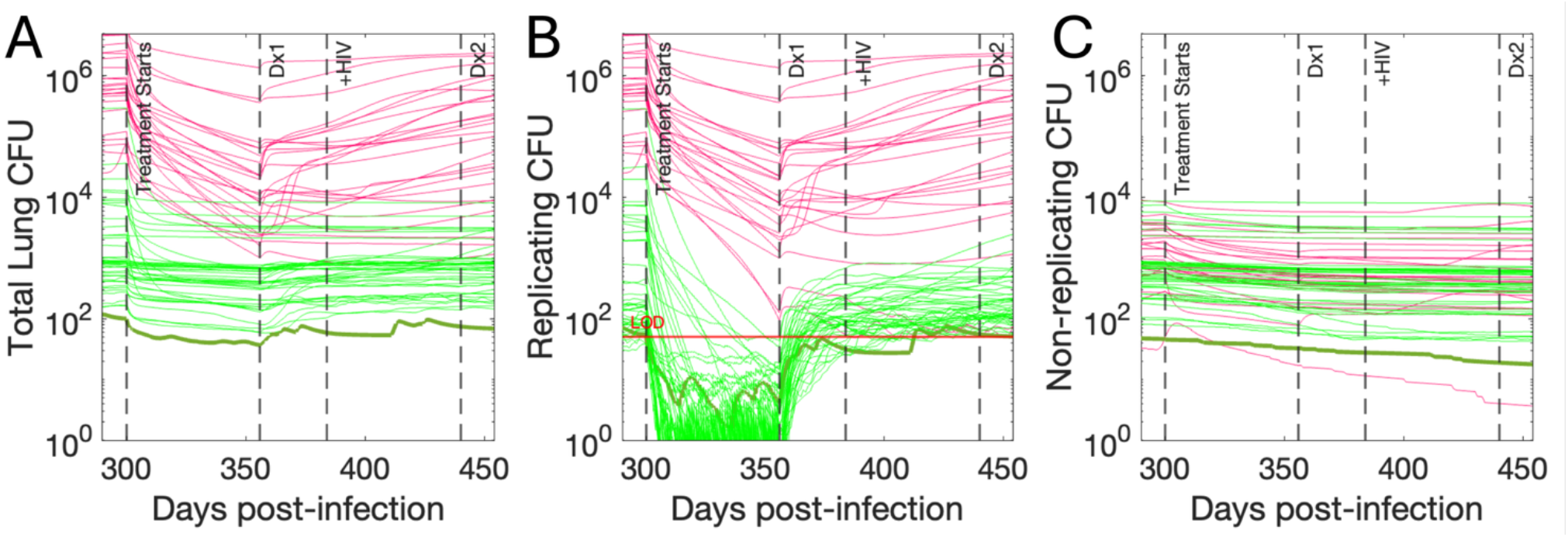
Relapse of virtual hosts after short course of HR and HIV-1 co-infection as in ^29^. Virtual host CFU levels over time for whole-lung total CFU (A), replicating (intracellular + replicating extracellular) Mtb only (B), and non-replicating Mtb only (C). Each line corresponds to a trajectory from one host that was categorized as having active TB prior to receiving HR (using NHP-like classification) then relapsing after the end of treatment. Key time points in days post-infection are shown, indicating the start of treatment, time of cure-treatment (Dx1), the onset of HIV, and the relapse follow-up time (Dx2). Cure-screened relapses are colored green, while those that would be excluded by cure-screening are colored red. The *Virtual Clinic score* assumes that non-replicating bacteria are not detectable, but the LOD of 50 is shown for replicating CFU. We have highlighted one representative virtual host (dark green) that, despite qualifying as relapse by this study design, may closely resemble subclinical disease or LTBI disease.

#### 2.2.2 Simulated relapse most closely matches caseum-forming animal model relapse rates

Since our virtual relapse studies reproduce NHP relapse, we survey similarities between *HostSim* and other relapse models by simulating several animal relapse studies. We chose these studies principally by whether sufficient details were included to perform analogous simulations, and the RMM studies tested many more regimens than those that we recreate here. For applicable comparison, we compare *in vivo* relapse rates to the frequency of relapse in virtual hosts with NHP-like and mouse-like active-TB (Methods). The NHP study that we replicate are the same simulations that we describe in Section 2.2.1. To replicate mouse studies, we allow the full virtual cohort of *n* = 500 to develop mature Mtb infection. We then administer a comparative antibiotic regimen, simulate for three months follow-up time with no treatment before relapse assessment. To compare *HostSim* with the RMM, we develop a test we call *Virtual lung homogenate* (Methods). All homogenate-based virtual diagnostic tests assume a 10-CFU LOD that can detect non-replicating Mtb, as true experimental LOD is challenging to estimate. For this reason, we only claim qualitative comparisons.

We find that our simulations recapitulate RMM and NHP trends, though relapse rates are closer to rates measured from animals whose granulomas contain caseum (NHPs and C3HeB/FeJ mice) than other mice (Table 2). As discussed in 2.2.1, we captured NHP relapse rates after HR+SIV using *Virtual PET/CT*, though all virtual hosts had at least one granuloma with >10CFU by follow-up. While *HostSim* recapitulates relapse rates of several RMMs (treated with RMZ and HRZ), we found that *HostSim* has higher relapse rates than RMMs for three regimens (HRZE, RMZE, BPaMZ). For HRZE and RMZE, the relapse rates were higher *HostSim* than in C3HeB/FeJ mice, which in turn had higher relapse rates than other RMMs. The relapse rate following BPaMZ treatment was substantially lower in RMMs compared to *HostSim*, likely due to the limited ability of Bedaquiline and pretomanid to penetrate into caseum^22,83^, which is absent in the RMM.

**Table 2:** Comparison between animal relapse studies and analogous simulated relapse rates. The first two columns show the reference animal relapse study and treatment regimen (see **Methods** for duration and dosage). The third and fourth columns show the corresponding *in vivo* and simulated relapse rates, respectively. † – This work did not measure relapse with CFU; this percentage reflects what percentage of NHPs had detectable lung CFU at time of assessment out of all NHPs whose CFU could be counted (See original study ^29^ Fig 3C).

| Replicated *in-vivo* animal species studied for relapse | Treatment regimen | *In vivo* relapse % | Virtual relapse % out of analogous sub-cohort |
| --- | --- | --- | --- |
| NHP (<sup>29</sup>) | HR with comorbid SIV | 67% | 51% (56/109)<br>(*Virtual PET/CT*) |
|  |  | 72%† | 100% (109/109)<br>(*Virtual granuloma homogenate*) |
| Mouse (<sup>41</sup>) | 2RMZ + 2RM | 42% | 50% (52/104) |
| Mouse (<sup>33</sup>) | 2HRZ+ 1HR | 87% | 74% (78/105) |
| C3HeB/FeJ Mouse (<sup>34</sup>) | 2HRZE+ 2HR | 13-27% | 53% (55/104) |
| Mouse (<sup>34</sup>) | 2HRZE+ 2HR | 0-7% |  |
| C3HeB/FeJ Mouse (<sup>34</sup>) | 2RMZE+ 1RM | 20-60% | 41% (43/104) |
| Mouse (<sup>34</sup>) | 2RMZE+ 1RM | 0-20% |  |
| Mouse (<sup>32</sup>) | 1BPamZ+1BPam | 7% | 48% (50/104) |

### 2.3 Translating *HostSim* relapse to clinical studies

We apply antibiotic treatment to *HostSim* to recreate human relapse studies and to distinguish frequency of persistence- and threshold-driven relapse in the human context. We show that cure-screened relapse is likely persistence-driven.

#### 2.3.1 - HostSim captures qualitative trends of human post-treatment relapse

To ensure that *HostSim* can bridge both animal and human relapse, we recreate several human relapse studies. For each, we allow the full virtual cohort of *n* = 500 virtual hosts to develop a mature Mtb infection, separately administer multiple indicated antibiotic regimens, then simulate hosts post-treatment for the indicated follow-up time before testing to assess cure or relapse. To capture the effect of study inclusion criteria, we examine relapse in virtual sub-cohorts based on human-like active-TB classification for fair comparison. Virtual hosts are screened for cure at treatment completion (except where noted) and assessed for relapse at follow-up time using virtual diagnostic tests analogous to the *in vivo* study (Methods).

Treatment of *HostSim* captures trends of human relapse studies (Table 3). Simulations show a decrease in relapse rates when using more potent and longer regimens, though simulated relapse rates are slightly higher than in clinical datasets. When simulating clinical studies, we find that *HostSim* provides an upper bound for human relapse likelihood, capturing more aggressive Mtb infection than the human data. Both real and simulated relapse rates are far higher in studies where cure-screening is not performed. Together, the results from this and the previous section show that a single model can mechanistically recapitulate relapse from multiple species and study designs, and that those reported relapse rates likely depend on several aspects of study design, notably including follow-up time and cure screening.

**Table 3:** Comparison between human relapse studies and analogous simulated relapse rates. The first three columns describe the reference human relapse study, treatment regimen (**Methods** for duration and dosage), and follow-up assessment times used to determine relapse. In the fourth and fifth columns, we show the corresponding *in vivo* and simulated relapse rates, respectively. **A** – This did not measure “relapse,” but rather was “still culture positive during continuing treatment”. This was still discussed as potential for relapse in literature^12^. **B** – This is a simplified percentage of 12/99 hosts with recurrent disease post-treatment; actual relationship between PET/CT inflammation and Mtb sterilization is discussed in the original study ^42^. **C** – Actual regimens unknown; we assume that these followed the standard of care.

| Replicated *in vivo* diagnostic test used to study human relapse | Treatment regimen | Follow-up time post treatment-completion | *In vivo* relapse % | Virtual relapse % out of analogous sub-cohort | Notes |
| --- | --- | --- | --- | --- | --- |
| Sputum culture conversion (<sup>71–73</sup>) | 2HRZE + 4HR | 14 days<sup>A</sup> | 60% | 88% (55/63) | Analogous simulations do not screen for cure. |
|  |  | 28 days<sup>A</sup> | 40% | 67% (42/63) |  |
|  |  | 60 days<sup>A</sup> | 10% | 22% (14/63) |  |
| PET/CT inflammation recurrence (<sup>42</sup>) | 2HRZE + 4HR | 6 months | 12%<sup>C</sup> | 43% (22/51) | Granuloma-scale test resolution measure via PET/CT |
| Rapid test (<sup>84</sup>) | 2HRZE + 4HR<sup>D</sup> | 2 years | 2.2% | 16% (60/378) | Clinical study admitted patients with any positive culture. |

#### 2.3.2 - Screening for cure at treatment-end characterizes subsequent relapse in simulations

In a simulated relapse study, we show that cure-screened relapse is most likely persistence-driven, regardless of diagnostic test accuracy. Specifically, we examine simulated rates of relapse one year after administering a 2-month short-course treatment of HRZE to 500 virtual hosts. We test for Mtb both at time of treatment completion and one year later by using *Virtual Mtb plate* (Methods), varying the LOD of both initial and follow-up tests from 0 to 100 CFU. We also vary whether or not virtual diagnostics can detect non-replicating CFU. Extremely sensitive tests are unlikely to classify patients as cured, especially if they detect non-replicating bacteria (Figure 5 bottom row). As tests become less sensitive, more virtual hosts diagnose as cured with higher levels of MDxC. Figure 5 shows that in simulations, more than half of cure-screened relapses are persistence-driven, regardless of LOD or test ability to detect non-replicating Mtb. All relapses are more frequently threshold-driven (>50%) if the diagnostic tests do not detect non-replicating bacteria. We observe the same outcome when simulating RMZE (Appendix A6), indicating that this trend maintains for regimens including drugs with more in-caseum bactericidal activity than isoniazid (here, moxifloxacin^85^).

**Figure 5:**
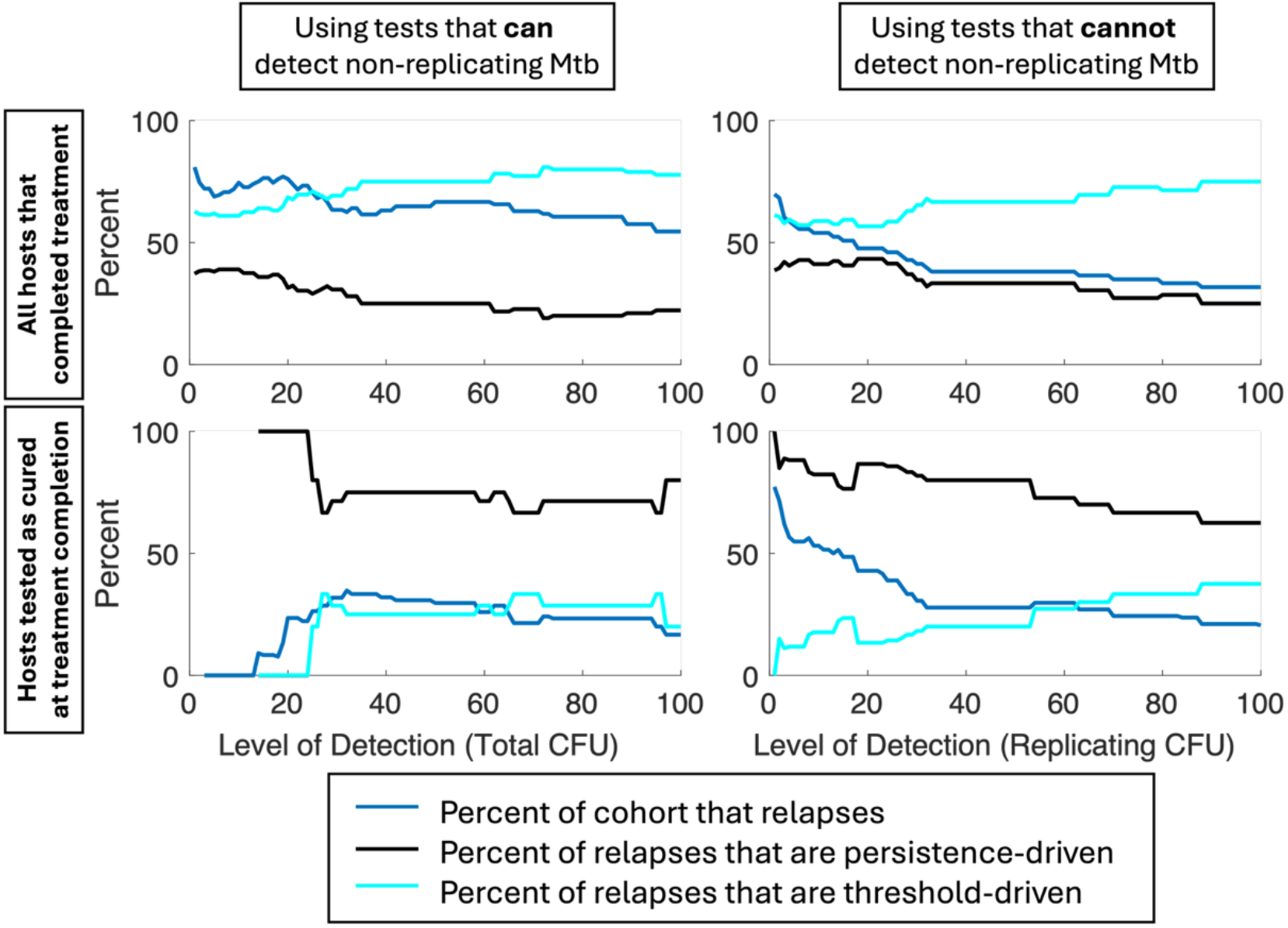
Persistence versus threshold relapse rates by test LOD. Each of our plots show how simulated relapse rates and the relative frequency of their driving mechanisms are impacted by LOD of the diagnostic test used to assess cure and/or relapse. These analyses used the *Virtual Mtb plate* test to mimic sputum culture conversion tests one year after active TB hosts complete 2 months of virtual HRZE treatment (Methods). We repeated each analysis assuming the *Virtual Mtb plate* either can (Left column) or cannot (Right column) detect non-replicating bacteria. We also repeated each analysis varying whether relapse is defined of all hosts that complete treatment (Top row) or only those that are misdiagnosed as “cured” upon treatment completion (Bottom row).

## 3. Discussion

Understanding how to safely shorten antibiotic treatment for TB is a critical step towards eradicating the world’s leading cause of death by single infectious agent. Aside from development of resistance, the main threat of shortening treatment regimen administration is relapse, where treated infections later recur. Experimental and clinical relapse cannot conclusively determine which of two hypothesized mechanisms of relapse drive what is observed in the clinic: persistence-driven relapse (caused by of non-replicating Mtb) and/or threshold-driven relapse (caused by incomplete sterilization of intracellular-Mtb)^12^. Moreover, it is not clear whether these mechanisms equally underpin clinical relapse or *in vivo* experimental models thereof. Understanding relapse is critical as current guidelines for re-treatment of drug-susceptible TB are thin and based only on observation^86^, leaving us little mechanistic understanding as we try to shorten treatment.

Simulations suggest that overall, relapses in patients screened for cure at treatment completion are more likely to be persistence-driven, and relapses in unscreened patients are likely threshold-driven (Figure 6). This finding was robust for all of the virtual diagnostic tests we examine, including varying whether the test could detect non-replicating CFU and a LOD varied between 1 and 100 CFU (Figure 5). Exceptions become more likely in patients that begin treatment with higher bacterial burdens; thus, future studies may find it fruitful to report relapse events stratified by pre-treatment CFU (e.g., high burden vs low burden of disease) if possible. In humans or animal models that form Mtb-controlling granulomas, follow-up studies may find that effective re-treatment involves using antibiotic regimens that effectively penetrate into caseum and kill Mtb trapped there in a non-replicating state—e.g., rifamycins such as rifapentine, rifabutin, rifalazil; and to a lesser extent moxifloxacin^85,87,88^. Unscreened patients may benefit more from second-line treatments like BPaLM (Figure 6). If our model assumption is indeed correct that persistence-driven relapse is characterized by a slow, continuous migration of Mtb from caseum, re-treatment regimens may be well-supplemented by low dose antibiotics that excel at killing non-caseum-bound bacteria, e.g., bedaquiline or moxifloxacin^89,90^. Finally, note that these results are not discussing the probability of relapse; indeed, predicting relapse from pre-treatment features is challenging^29^. Rather, our simulated results speak to the relative likelihood of mechanisms underpinning relapse when it occurs.

**Figure 6:**
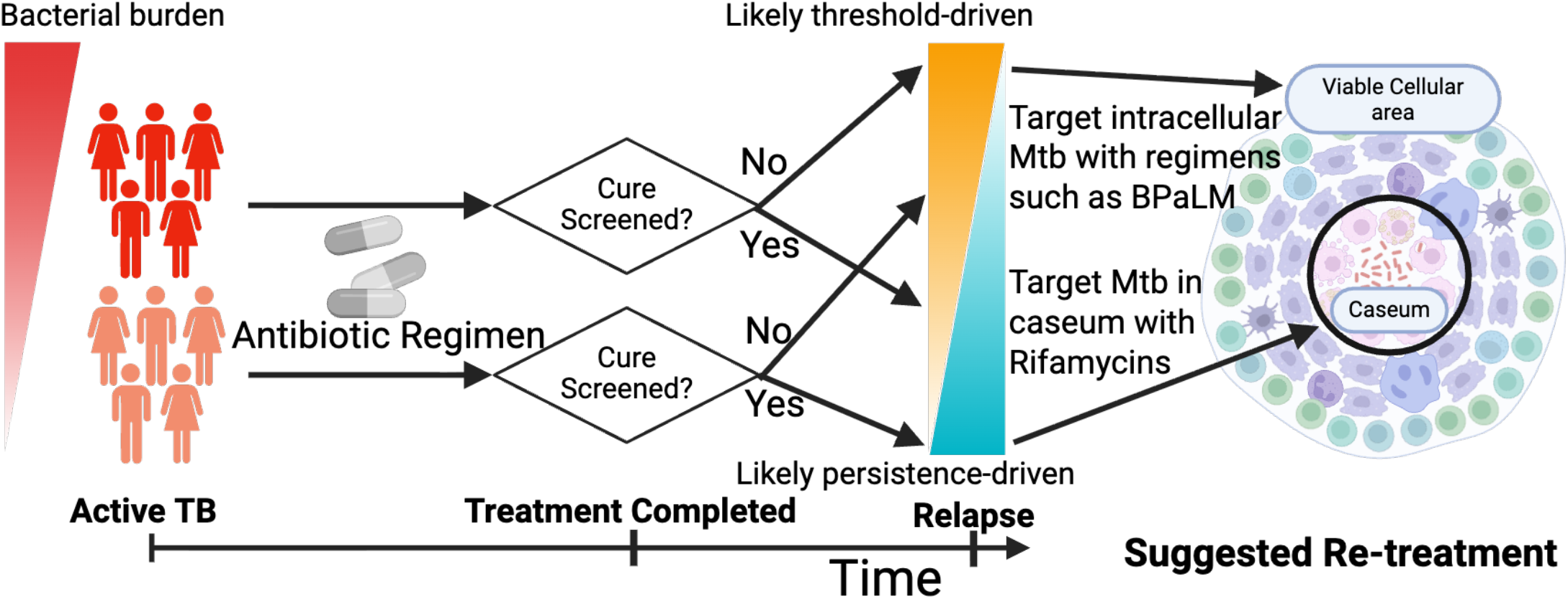
Predictors of relapse modality in simulations. For a variety of simulated relapse scenarios, we find that relapse after a host has microbiologically been screened for cure is more likely to be persistence-driven relapse (non-replicating Mtb trapped in caseum). The overall probability of threshold-driven relapse increases if a host has high CFU levels pre-treatment. Re-treatment strategies can be informed based on these considerations. (Created in BioRender. Michael, C. (2026) https://BioRender.com/u3x50cy.)

Our computational model bridges relapse studies in both human and animal contexts. We simulated reasonable rates of misdiagnosed cure after a variety of short-course antibiotic treatment regimens, focusing on antibiotics presently used in standard-of-care treatments such as HRZE^9,23^ (Methods). Simulations suggest that several regimens, including HRZE and most mono-treatments, are far more likely to result in MDxC, where non-sterile infection falls below a LOD at the time of treatment completion, but is not cleared (Figure 3). To capture mechanisms of how MDxC leads to relapse, we also incorporated a mechanism allowing non-replicating Mtb to be transported from caseum, allowing for new granuloma reseeding as we have done previously^77^. With it, we were able to simulate a recent short-course treatment relapse study in NHPs^29^.

Calibrating and validating a relapse model pose delicate problems, as calibration targets (such as reactivation rates) and validation targets (such as MDxC trends and NHP relapse rates) do not come from a single or full population of relapsing Mtb-infected individuals. Typically, to validate a computational model of, say, relapse, we would generate several virtual hosts that mechanistically recapitulate a relapse-like phenomenon. The multitude of measurements and proxies for relapse, gathered by study design, do not collectively paint a picture of a single “type of relapse” *a priori*. Indeed, our results tell us that a host’s species, state prior to treatment, whether the host was cure-screened, and the method of that screening all subtly inform what phenomena we operationally classify as relapse and consequently impact the mechanism of relapse.

Notably, we find that reported relapse rates are considerably impacted by study-specific relapse definition— specifically, whether it is based on (1) cure status upon treatment completion (not standardized between clinical and experimental studies) and (2) whether subclinical patients are admitted to relapse studies (Tables 1 and 3). This remains true when recreating several other relapse studies in humans, mice and NHPs. When analyzing datasets as closely as possible to their *in vivo* counterparts, we recapitulate qualitative trends of relapse (e.g., longer treatment duration leads to lower rates of relapse). However, we also find a paucity of calibration-relevant measurements in the literature, such as LOD of many diagnostic tests used. Our simulated relapse rates are higher than those reported in literature (Table 2). Higher relapse rates are reasonable since our virtual diagnostic tests are optimistically sensitive, whereas true sensitivity based on Mtb in sputum or BALF is likely lower; in absence of literature connecting granuloma and sputum or BALF CFU, we assume one replicating CFU in the lungs corresponds to one CFU in a sputum or BAL sample. For RMMs, our relapse rates are closer to those observed in caseum-forming C3HeB/FeJ mice than non-caseum-forming BALB/c mice. The low rates of relapse among non-C3HeB/FeJ mice reinforce that without caseum, RMMs exclusively capture threshold-driven relapse. Our simulations show that threshold-driven relapse is unlikely to occur for lower-burden hosts who are treated for longer periods of time, especially in populations screened for cure.

We noticed that some measures of relapse that employ extremely-sensitive tests may capture perseverance of subclinical disease with Mtb levels below LOD, rather than symptomatic clinical relapse. Proxies for relapse from animal models (e.g., Mtb-positivity of tissue homogenates) may implicitly classify as relapse even if only small amounts of Mtb are present, which may more closely resemble delayed, bacteriologically-detectable asymptomatic infection after a MDxC (see dark green line in Figure 4). Still, information is valuable as recurrent subclinical disease is plausibly a silent contributor toward drug resistance with regimen-specific risks. An absence of data surrounding how diagnostic test sensitivity relates to symptoms leaves little guidance for further simulations, as simulation results depend on assumed LODs.

We have made several important simplifications to reduce model complexity. We do not model drug resistance, though drug resistance is believed to correlate with relapse rates^16^. Lymph nodes are known to be important reservoirs of Mtb during TB disease and relapse, so we will combine our recent model of lymph node infection into *HostSim* in future work to see how this impacts our predictions^6,17,19,29^. Moreover, we calibrate our persistence-like mechanism using reactivation rates subsequent to TNF_α_ depletion (Methods), which may ignore subtleties of TB disease during immunosuppression, evident in how TNF_α_-induced reactivation of TB still often test skin-test negative, and may have other qualitatively distinct immune factors^12^. Our simulations do not capture lung cavitation, and the animal models we used for calibration did not exhibit significant cavitary disease. Lung cavitation is a known risk factor for relapse^91,92^ that bears distinctions from non-cavitating disease dynamics, including differences in oxygen concentration and recommended treatment, results in extremely high levels of CFU in lung tissue and sputum^91,93,94^, which we assume makes MDxC unlikely. However, if cavitating patients are similar to our virtual high-CFU hosts, then relapse is more likely threshold-driven. Finally, the potential for diagnostic tests to yield false positive results follows from test-specific causes that are beyond the scope of this work. For example, simulating false-positive results from immune assays likely requires detailed representations of markers used in that assay, whereas predicting false positive results from culture-based tests would depend on detailed representation of BALF sampling or laboratory contamination.

There are gaps in literature precluding any comprehensive model of relapse. Virtual diagnostics rely on a coarse-grain representations of patient states—e.g., predicting symptoms via the fold-increase of CFU over 200 days, or assuming direct proportionality between CFU in BALF, sputum and lung tissue. Such abstractions are necessary as biological studies relating host infection state appear noisy, and it is not mechanistically clear why some hosts with active TB are culture-negative while some subclinical hosts are culture-positive^4,95^. Our simulations cannot mechanistically distinguish between LTBI and subclinical disease as we lack the mechanistic detail connecting lung state to these clinical markers. However, *HostSim* (as many model frameworks) is modular, allowing us to refine components as datasets become available to form a more comprehensive symptom and diagnostic model of TB to predict relapse. One major benefit of being able to predict within-host Mtb and diagnostic-relevant inflammatory responses from patient-specific datasets is the potential to create a TB treatment *digital twin*, a personalized model that uses patient data to forecast specific patient outcomes during treatment. Digital twins have successfully been created for a variety of complex diseases, but require a method to use limited patient-specific datasets to inform model states^96–99^. Once it is possible to predict infection state from patient datasets, we would be able to create a TB relapse digital twin that could provide case-personalized decision support.

In conclusion, simulations based on a combination of human and NHP datasets can plausibly recreate observed relapse rates, but those relapse rates are a union of both persistence- and threshold-driven relapse. In regimens that effectively penetrate caseum, simulations well-match murine relapse rates, and do not otherwise (though are closer to caseum-forming C3HeB/FeJ mice in this case). On the other hand, it is challenging to link observed human relapse without leaning on Mtb burden as a proxy until we know more about how continuously Mtb sheds from caseum into BALF, or how this relates to clinical symptoms. It appears that, at least while assuming caseum-bound Mtb are not clinically detectable, screening for cure within individual hosts as part of relapse study design essentially removes from clinical studies the very threshold-driven relapse events best modeled in mice. While both types of relapse are relevant at bedside, in clinical trials, and in experiments; separating studies of relapse risk by-mechanism may shed light on overall treatment efficacy, particularly for regimens that are caseum-limited.

## 4. Methods

Recently, we developed a detailed and complex *in silico* whole-host model of pulmonary Mtb infection and treatment, *HostSim*^61,65,66,100^. Model details are in section 4.1 and on the online technical description of *HostSim* at http://malthus.micro.med.umich.edu/lab/supplements/Host-Sim-3/. *HostSim* contains a whole-host PK/PD model, which we used to perform virtual pre-clinical trials to characterize bactericidal activity of various antibiotic regimens^61^.

For all analyses, we generate and calibrated a set of *n* = 500 virtual hosts, each with 13 primary granulomas. Parameter values defining each host were sampled using the Laten hypercube sampling method from calibrated and validated parameter ranges as previously described^61^. Our untreated virtual cohort has 89.2% virtual hosts with LTBI, 0.8% sterilizing, and 10% virtual hosts with active TB disease at 300 days post-infection.

### 4.1 Overview of *HostSim*

Briefly, *HostSim* is a multi-scale hybrid computational model of pulmonary TB, including multiple lung granuloma agents within a single virtual host lung that is coordinated with both a blood compartment and an uninfected virtual lymph node system^61,65^. As pulmonary TB is typically contained to these 3 compartments, we refer to this as a “whole host” for TB. Each virtual granuloma is defined by a set of ordinary differential equations that describe interactions between host immune cells (CD4+ T cells, CD8+ T cells, and macrophages) communicating, polarizing, and differentiating in response to cytokine signals (IFN-γ, TNF_α_, IL-4, IL-10, and IL-12). These immune cells respond to three distinct Mtb subpopulations: intracellular (within macrophages), extracellular within the granuloma, and non-replicating. We model all non-replicating Mtb as being trapped within caseum. As Mtb are killed, antigen accumulates and traffics to the lung draining lymph node system, which is represented as another set of ODEs. The lymph node ODE tracks priming of Mtb-specific CD4+ and CD8+ immune cells which migrate to the host’s blood, which is the third and final set of ODEs in *HostSim*. CD4+ and CD8+ Mtb-specific and nonspecific T cells may then be recruited to the virtual lung granulomas based on the granuloma state. This model also has stochastic elements such as granuloma dissemination (i.e., seeding of new granulomas) and the persistence-driven mechanism of Mtb transport from caseum to macrophages.

Each ODE term and inter-physiological compartmental transition term describe key dynamics of pulmonary TB, including both untreated infection progression and the PK/PD governing treatment efficacy. The terms in *HostSim* include (i) metabolic differences between replicating Mtb (internalized by macrophages or not) and non-replicating Mtb; (ii) dynamic priming of T cells and the (de)activation of monocyte-derived macrophages; (iii) the dynamic accumulation of caseum based on macrophage necrosis; (iv) antibiotic penetration into caseum; and (v) synergistic/antagonistic drug-drug impacts on PD. Details of parameter ranges and ODEs are in Appendix B.

### 4.2 Classifying virtual host state

To systematically examine case-specific patient outcomes with or without treatment, we need to determine how those outcomes are measured.

#### 4.2.1 Classifying virtual host subclinical versus active TB by species

Estimating which virtual hosts suffer active disease is a nuanced challenge, as there is no currently known mechanism connecting CFU (lung or BALF) to symptoms. As an estimate, if CFU levels reach a species-specific threshold for fold-increase over when we would have expected hosts with LTBI to stabilize (∼200 days), then we label the hosts as having active TB. We also predict hosts with large-CFU levels (>10,000 for over 30 day timeframe) also experience active TB, as we have in previous work^61^. Hosts with CFU levels below a given LOD are listed as “sterilizing” (as we assume clearance cannot be distinguished from paucibacillary infection). All other virtual hosts are considered to be subclinical or LTBI. See Appendix A4 for more details.

#### 4.2.2 Virtual diagnostic tests

Clinically or experimentally, disease state is assessed using one or more diagnostics (Appendix Table A1). To assess virtual host outcomes, we mimic those detection methods *in silico* using virtual diagnostics (Table 4). These are the building-blocks of testing for more complex outcomes like relapse or reactivation. By tuning each of the virtual diagnostic tests in Table 4, we are able to mimic most of the virtual diagnostic tests used to assess host disease state *in vivo* (Appendix Table A1).

**Table 4:** Virtual diagnosis overview. Each virtual diagnostic test has free parameters adjustable to mimic one or more real diagnostics. See Methods for details.

|  |  |  |
| --- | --- | --- |
| **Diagnostic Test name** | **Brief description** | **Free parameters in their calculation** |
| *Virtual Mtb plate* | Evaluating for presence of Mtb derived from a patient sample. Patient sample can be derived from a lung homogenate culture, smear, etc. Descriptions of how we configure *Virtual Mtb plate* to mimic various tests are in Methods.<br>Binary - returns 1 (*not cured*) or 0 (*cured*). | - LOD<br>- Ratio of CFU in sample to CFU in lung (or granuloma)<br>- Whether non-replicating Mtb may be present in virtual sample<br>- Whether or not recently-dead Mtb may be counted towards CFU levels |
| *Virtual PET/CT* | Examining numbers of metabolically-active immune cells in granulomas allows for the prediction of 18F-fluorodeoxyglucose (FDG) avidity (output of PET/CT scans<sup>101</sup>). Sufficiently increasing FDG avidity indicates a positive diagnosis.<br>Binary - returns 1 (*Positive Increase*) or 0 (*No change*). | - LOD<br>- Fold-increase of FDG-avidity required for a positive diagnosis<br>- Predicted relative contribution of metabolically-active immune cells toward FDG avidity (Methods). |
| *Virtual clinic score* | Implemented as a combination of the above diagnostics. We assume that if a virtual host's CFU increase is uncontrolled or if a granuloma disseminates, they will have symptoms and receive a clinical score.<br>Binary - Returns 1 (Either or both of the above are both positive, likely symptomatic), or 0 (Neither of the above are positive, likely asymptomatic). | - All of the parameters of *Virtual Mtb Plate* and *Virtual PET/CT* |

A virtual diagnostic test takes a virtual host at a given time and assesses their infection state. All host-scale tests assume that pulmonary TB infection states can be precisely measured by looking at the lung state, but this may be expanded in future studies to include more detailed regarding the role of lymph node infection^19^.

##### Virtual Mtb plate

This virtual test recreates various culture, smear, or NAAT-based tests (Table 5) to assess a host as either *cured* or *not cured* at a given time. Virtual Mtb plates sum together CFU counts of relevant Mtb bacilli and compare them against a LOD. The test returns *not cured* if the sum of Mtb exceeds the LOD, and it returns *cured* otherwise. The exact definition for which Mtb are relevant is configured by checking a number of biological assumptions; in principle, we can calibrate these further once more biology is known.

**Table 5:**
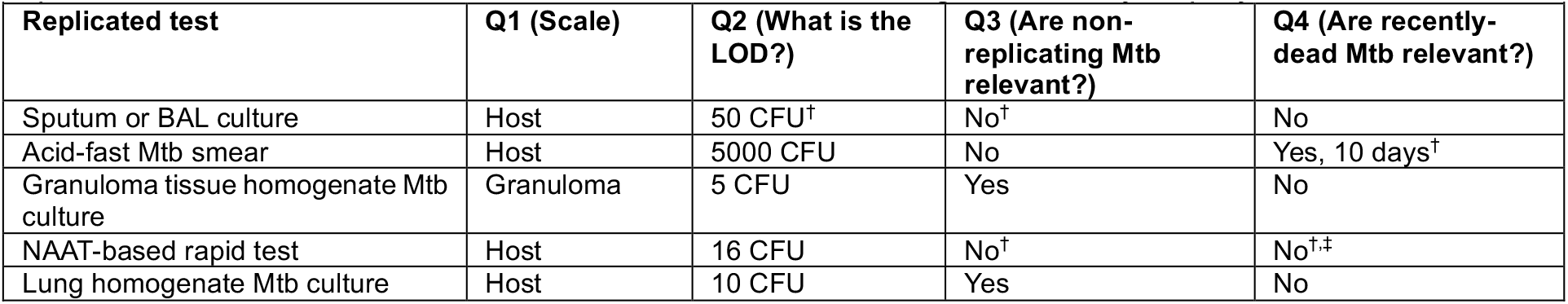
Choices and assumptions that we make to configure virtual Mtb plates into various clinical diagnostics *in silico*. The configuration-based questions are: (1) Does the diagnostic look at the entire host (H) or a specific granuloma tissue (G)? (2) What is the LOD of the test? (3) Are non-replicating or caseum-bound bacteria detectable with this test? (4) Are dead Mtb able to be detected in this test? If so, how long can bacilli be detected for after they have been killed? †Assume direct proportionality between lung-tissue CFU counts and sputum/BAL CFU counts. ^‡^This assumes that the nucleic acid being tested decays rapidly.

##### Virtual PET/CT

Combined positron emission tomography (PET) and computed tomography (CT) is a radiographic measure of inflammation (Appendix A1). As previously^100^, we measure virtual FDG avidity (measured by PET/CT) as a weighted sum of metabolically-active immune cells, including activated macrophages and T cells within each granuloma. In summary, FDG avidity is calculated as

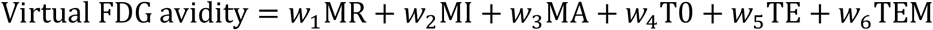

where ⟨_*wi*_⟩ = ⟨0,5,6,2,4,3⟩ are the relative weights of resting, infected, and activated macrophages; and primed, effector, and effector memory cell populations within a granuloma. Resting macrophages have a contribution of 0 as we assume they are at the same level of background activity as the uninvolved lung tissue.

The *Virtual PET/CT* test classifies hosts as relapsed if:

1. Any new granulomas disseminate (Appendix A5).
2. There is >20% increase of virtual FDG avidity between time of treatment completion and time of assessment for any granuloma within the host.

##### Virtual clinic score

Virtual clinic score is a combination of the above tests as a proxy for clinical evaluation. We assume that a host will present with symptoms if new granulomas continue to form, and that a clinician has access to a rapid test (e.g., Xpert^102^). A host is determined to be *cured* if none of the following return *not cured*:

1. A *Virtual Mtb plate* with LOD = 16 CFU (as described in Table 5), assuming that non-replicating bacteria cannot be detected via a rapid test such as Xpert^102^ (see Appendix Table A1).
2. If any new granulomas have formed within the host since the last virtual diagnostic, even below LOD— we assume that this is indicative of symptom recurrence as is assumed in ^29^.

### 4.3 Mechanism for persistence-driven relapse: stochastic transport of Mtb from caseum

To simulate both persistence- and threshold-driven relapse, we include persistence and threshold mechanisms in *HostSim*. Simulations allow an activated immune system (such as neutrophils) to transport Mtb from caseum, transitioning them to an intracellular state as suggested by our previous work^77^. We assume that the persistence mechanism is involved in both relapse and reactivation. Subclinical hosts may go years without reactivation— only 10% of subclinical infection hosts with no comorbidities reactivate at any point in their life^4,103^—so we calibrate our reactivation mechanism for studying relapse model in the context of reactivation induced by TNF_α_ depletion. Details are in Appendix A2.

### 4.4 Simulated antibiotic regimens

Relapse has been studied in the context of shortening standard antibiotic regimens. These typically include a combination of Isoniazid (INH; H), Rifampicin (RIF; R), Ethambutol (ETH; E), and Pyrazinamide (PZA; Z), and together are the historic standard treatment regimen HRZE^9^. Second-line regimens for RIF-resistant TB include a combination of Bedaquiline (BDQ; B), Pretomanid (PTM; Pa), Linezolid (LZD; L), and (sometimes) Moxifloxacin (MXF; M), called BPaL(M) ^7,37^.The standard treatment is 6-9 months of HRZE, or 2 months of HRZE and 4 months of HR^104^. Several studies have tested for sputum culture conversion after only 2 months of antibiotic treatment^38^.

We simulated two families of antibiotic regimens, summarized in Table 6. To determine variability in the level of non-detectable bacteria across multiple regimens, we reproduce a family of regimens including H, R, Z, E, B, Pa, L, and M tested in our previous work^61^ (group i). We also simulate scenarios wherein virtual hosts are treated with multi-phase regimens (group ii), such as the WHO standard of care (Regimen 2HRZE4HR) that discontinues use of PZA and EMB after two months. After virtual treatments end, we continue simulations for without treatment to examine relapse events, though regimens may be assessed at more than one follow-up time (Table 3).

**Table 6:** Regimens from previous studies recreated using *HostSim*. This table is partly based on Table 5 from our previous work^61^. Columns labeled by antibiotic name (e.g., INH) indicate the human-equivalent dosage of that antibiotic in human-equivalent mg/kg, administered daily. Each regimen in set (i) lasts 6 months (180 days). Regimens in set (ii) are split into phases, with each phase duration and the drugs administered during that phase indicated by regimen name—e.g., 2HR1R indicates 2 months of INH+RIF followed by 1 month of RIF monotreatment. *Antibiotic halted after initial phase. ^†^Culture positivity was reported during treatment, so we predicted relapse at days-post-treatment-start for these specific regimens and times. ^‡^Specific timing of virtual HRZE and virtual SIV are given in the Results text, and are adapted from an NHP model^29. §^Dose adjusted to the human equivalent standard dose.

| Regimen Name | INH | RIF | PZA | EMB | BDQ | PTM | LZD | MXF |
| --- | --- | --- | --- | --- | --- | --- | --- | --- |
| **Regimen set (i)** - Marmoset studies from a previous *in silico* / NHP study<sup>105</sup> (set reproduced from our previous *HostSim* study<sup>61</sup>). |  |  |  |  |  |  |  |  |
| RMZE | - | 10 | 25 | 20 | - | - | - | 7 |
| BPa | - | - | - | - | 20 | 20 | - | - |
| BPaL | - | - | - | - | 20 | 20 | 90 | - |
| BL | - | - | - | - | 20 | - | 90 | - |
| RM | - | 10 | - | - | - | - | - | 7 |
| HRZE | 5 | 10 | 25 | 20 | - | - | - | - |
| PaL | - | - | - | - | - | 20 | 90 | - |
| Bedaquiline | - | - | - | - | 20 | - | - | - |
| Pretomanid | - | - | - | - | - | 20 | - | - |
| RZ | - | 10 | 25 | - | - | - | - | - |
| HZ | 5 | - | 25 | - | - | - | - | - |
| Moxifloxacin | - | - | - | - | - | - | - | 7 |
| Pyrazinamide | - | - | 25 | - | - | - | - | - |
| Rifampicin | - | 10 | - | - | - | - | - | - |
| Isoniazid | 5 | - | - | - | - | - | - | - |
| **Regimen set (ii)** - Multi-phase regimens used in standard of care and/or experimental relapse studies. |  |  |  |  |  |  |  |  |
| 2HR + SIV<sup>‡</sup> | 6 | 10 | - | - | - | - | - | - |
| 2HRZE4HR | 6 | 10 | 25* | 20* | - | - | - | - |
| 2RMZ2RM<sup>§ (41)</sup> | - | 10 | 25* | - | - | - | - | 7 |
| 2HRZE2HR<sup>§ (34)</sup> | 5 | 10 | 25* | 20* | - | - | - | - |
| 2RMZE1RM<sup>§ (34)</sup> | - | 10 | 25* | 20* | - | - | - | 7 |
| 2HRZ1HR<sup>§ (33)</sup> | 6 | 10 | 25* | - | - | - | - | - |
| 1BPamZ1BPam<sup>§ (32)</sup> | - | - | 25* | - | 20 | 20 | - | 7 |

## Acknowledgements

C.T.M. was supported by the Molecular Mechanisms in Microbial Pathogenesis Training Program (T32 AI007528). This work was by National Institutes of Health Grant R01 AI50684 (D.K.) and the Center for Data-Driven Drug Development and Treatment Assessment (DATA; D.K. and M.B.). We thank Paul Wolberg for computational assistance and support.

## Footnotes

a We refer to lung granulomas unless otherwise specified, though granulomas commonly form in thoracic lymph nodes^17^, likely impacting adaptive immune priming^18,19^; granulomas may also form during extrapulmonary TB—e.g., liver^20^.

b “Persister Mtb” are not synonymous with “persistence-driven relapse”; describing a Mtb bacillus as a “persister” typically means it survives antibiotic treatment^26^ and could be used to describe Mtb in either persistence- or threshold-driven relapse.

c We use the phrasing of MDxC rather than “tested as false-negative” to avoid confusion between “Mtb-negative” versus “cure-negative”.

d Inflammatory markers are often used to diagnose TB, such as QuantiFERON-TB Gold In-Tube, which measures interferon gamma release. More diagnostic methods are shown in Appendix A.

