## Appendix A for "Cure-screening predicts mechanism of post-antibiotic relapse in Tuberculosis"

### A1 - Extended Tuberculosis background

#### *A1.1 - Granulomas*

To understand observed variability in responses to TB treatment, it is necessary to consider the immunology and heterogeneity of *lung granulomas* that arise during Mtb infection. Granulomas are spatially-complex structures that both physically and immunologically quarantine Mtb[1]. Most often, granulomas, feature an outer region containing multiple innate and adaptive immune cells and bacteria and  a necrotic, hypoxic region filled with *caseum* - an aggregate of dead cells, cellular debris, and non-replicating Mtb[1]. Both granulomas and caseum within them are a primary pharmacokinetic and pharmacodynamic barrier to TB treatment[2], as antibiotics do not uniformly penetrate granulomas or into caseum and are not uniformly effective at killing Mtb across metabolic states. Finally, wide variations of individual granuloma outcomes are observed within individual Mtb-infected hosts[3].

#### *A1.2 - Combined positron emission tomography (PET) and computed tomography (CT)*

PET/CT measures the metabolism of the radiolabeled 18F-fluorodeoxyglucose (FDG) as a measure of inflammation[4,5]. FDG avidity, measured from PET/CT, is known to correlate with CFU levels in NHPs[6]. It was found that many clinically-cured patients have ongoing inflammation and detectable Mtb mRNA post-treatment[7], yet inflammation reduction has been noted during antibiotic treatment in mice[8,9] and NHPs[10].

#### *A1.3 - Diagnostic tests for clinical TB*

Multiple diagnostic tests have been used to diagnose TB, assess for LTBI, and experimentally assess animals or *ex vivo* tissues for Mtb. In Table S1 we provide an overview of many common methods, based in part on a previous review[9].

**Table A1**: Common methods for assessing *M. tuberculosis* infection and/or TB disease *in vivo*, based in part on [9].

| Diagnostic | Advantages | Drawbacks | CFU Level of Detection (LOD) | References |
|---|---|---|---|---|
| Solid or liquid culture of BAL | Gold standard for positive diagnosis when repeated | Can take weeks to months, requires 2+ tests. Poor predictor of relapse during early infection | 1 CFU within BALF or sputum (ideally) | 9,11,12 |
| WHO-approved rapid diagnostics (e.g., Xpert ultra) | Fast, accurate | Expensive, can bear false positive after previous infection | 16 CFU/mL (BALF or sputum) | 13,14 |
| Acid-fast sputum or BALF smear | Fast, inexpensive | Yields false positives after previous infection, false-positive rate may depend on treatment regimen | ~5,000 CFU/mL (BALF or sputum) | 9,15,16 |
| Whole-lung or tissue homogenate | Highly accurate, detects infections during active TB or subclinical disease | Animal model only (Performed post-necropsy) | ~5CFU/sample | 17(NHP); 18,19(Mouse) |
| Clinical radiographic tests (Chest X rays / Chest CT) | Non-invasive, fast, widely available, used to detect evidence of progressive TB | Does not quantify or assess inflammation levels; unknown precise Mtb LOD; poor specificity; reader inconsistency. | Varies | 20,21 |
| Other radiographic tests (e.g., checking for dissemination using a 18F- | Non-invasive, spatially resolves individual granuloma disseminations, | Measures gross fibrosis, cavitation, or inflammation; typically associated with disease severity or | - Not a direct measure of CFU, but FDG avidity from PET/CT is proportional to CFU[6]. | 6,17 (NHP); 4,20(Human) |

| | | | | |
|---|---|---|---|---|
| fluorodeoxyglucose radio-tagged PET/CT probe) | correlates with Mtb burden, longitudinal data available | bacterial burden but can be host response, which may persist post-cure; unknown precise Mtb LOD | (Returns a measure of inflammation in host) | |
| Immune assays | Some, like tuberculin skin tests (TSTs) or interferon gamma release assays (IGRAs), only measure Mtb infection. Standard diagnostic tests include TSTs and QuantiFERON-TB Gold In-Tube (an IGRA). | Unclear as to how long inflammation lasts after Mtb are cleared, not always able to distinguish active TB from subclinial disease<br>Low quality evidence for the ability to detect LTBI | - Not based on CFU<br><br>(Returns a positive/negative based on sufficient presence/absence of TB-associated immune markers) | [22–25] |
| Recording clinical TB indicators: New prolonged cough (≥14 days), weight loss, etc. | Noninvasive. Motivates the patient / care team to test for TB that may be otherwise missed | Insufficient to diagnose TB by themselves | Unclear connection to CFU<br><br>(Gives clinical picture not linked to infection measures) | [26] |

### A2 - Detailed Persistence Mechanism of Relapse

#### *A2.1 - Persistence Model of relapse description*

The ODE components of our hybrid ODE/agent-based model yield continuous trajectories of cell populations, including both caseum-bound bacteria $B_N$ and intracellular bacteria $B_I$. We capture persistence by using a stochastic impulse into our system, called *release events*. When a release event occurs, we move one unit from non-replicating bacteria trapped within caseum, $B_N$ and add one unit to the intracellular bacteria population outside of caseum, $B_I$. We remove one unit from resting macrophages and add one to the number of infected macrophages, representing macrophages internalizing that single, trafficked bacterium. We allow for a release event to occur at most once-per-day occurring within each granuloma using the following probability:

$$Prob(\text{Release Event}) = \frac{{B_N}^2}{{B_N}^2 + {C_{N,\text{release}}}^2} \cdot \left(1 - \frac{[\sum \text{T}]^2}{[\sum \text{T}]^2 + {C_{T,\text{release}}}^2}\right)$$

where $B_N$ is the population size of non-replicating bacteria trapped in caseum, $\sum T$ is the total number of Interleukin-10 producing T-cell species in our simulation (as reductions of such T cells are associated with higher rates of reactivation[27] and other presentations of Mtb dissemination[28]). The parameters $C_{N,\text{release}}$ and $C_{T,\text{release}}$ are phenomenological parameters to calibrate.

#### *A2.2 - Calibration of persistence mechanism using anti-TNFα immunosuppression*

Reactivation occurs both in humans and NHPs and is inducible by TNFα-neutralizing and anti-TNFα therapy[29–36]. Anti-TNF medications such as infliximab, etanercept, and adalimumab either bind TNF or inhibit its production. Not all mechanisms of TNF depletion have the same impact on TB outcomes *in vivo*; reactivation of LTBI is roughly five times more likely when taking infliximab (a monoclonal TNF antibody) versus etanercept (a fusion protein of TNF)[36–39]. In *HostSim*, we coarse-grain these mechanisms together, and achieve the drug effects by zeroing-out all production terms to TNF—most similar to the action of monoclonal antibodies such as infliximab.

Previous studies have observed 70-80% reactivation of LTBI in NHPs when TNFα is neutralized by either Adalimumab (8 weeks) or p55-TNFRI (19 weeks)[31,32], though this reactivation can be slowed by a combination treatment of Isoniazid and Rifampicin or with Metronidazole[31]. Approximately half of a cohort of NHPs with depleted TNF showed new granulomas within 8 weeks[30]. In humans, the precise likelihood of reactivation with TNF suppression is difficult to estimate because the size of the non-reactivating population is unknown. However, a hidden Markov model in 2004 estimated that approximately 20% of humans with LTBI would reactivate each month[33]. We also know that out of the patients who experience relapse consequent to treatment with Infliximab, around 44% relapse within 90 days of receiving treatment[40], although other anti-TNF drugs such as Etanercept and Adalimumab report lower levels of reactivation[36,38–40].

Beyond CFU, dissemination has also been noted in TNF-driven reactivation. In NHP experiments, reactive disease has been measured by the presence of new disseminations[30] and, in that study, 50% of NHPs with subclinical TB and had anti-TNF treatment, experienced reactive disease. Elsewhere, it was reported more than half of NHPs treated with TNF-neutralizing drugs exhibited signs of reactivation[32], and another study showed up to 70% reactivation[31,32]. We calibrate our model to have more than half of TB-controlling hosts result in relapse upon depletion of TNF for eight weeks.

We calibrate our virtual persistence mechanism (i.e., $C_{N,\text{release}}$ and $C_{T,\text{release}}$) by recreating these expected trends in TNF-depletion studies, notably to have reactivation rates between 70% to 80%. As a coarse-grained model for predicting symptomatic TB, we assume that fold-increase of CFU between 2x and 10x in lung tissue is indicative of symptoms. We simulate Mtb infection progression in a cohort of 500 virtual hosts for 365 days. At day 365, we virtually suppress TNF in all hosts by setting TNF production terms to 0 for a period of 8 weeks. We then measured the portion of previously-LTBI hosts that became clinically-infected. We manually calibrated our parameters to $C_{N,\text{release}} = 10$, $C_{T,\text{release}} = 300$, which results in 74.4% (88.8%) of LTBI hosts experiencing at least a 10-fold (2-fold) increase of CFU or new disseminations.

*A2.2 - Further validation with virtual IFNγ depletion*

Another immune cytokine that serves as a validation target is IFNγ, a pro-inflammatory cytokine that is produced by T cells and is a key player of one pathway by which the immune system regulates PD-1 / PD-L1[41]. The authors found few studies of IFNγ depletion experiments during Mtb treatment, an experiment in the 90s showed that IFNγ knockout mice suffer rapid and fatal Mtb infections[42], although more specialized mouse models may further verify these results (e.g., the C3H/HeJ and C3HeB/FeJ "Kramnik" mice that form necrotic lesions[18]). There have been case studies of humans being treated for cancer with PD-1 inhibitors that subsequently developed active tuberculosis[43]. The IFNγ blocker Emapalumab (AKA Gamifant), used to treat hemophagocytic lymphohistiocytosis, requires screening for latent TB infection and prophylactic treatment for subclinical TB[44].

For further validation, we performed a virtual IFNγ depletion experiment similar to our TNFα depletion experiment above, assuming that all new IFN$\gamma$ production is halted. After eight weeks of IFN$\gamma$ depletion, we find that 50% of virtual hosts experience at least a 2-fold increase of CFU or new disseminations, and 0.5% of latent hosts experience at least a 10-fold increase of CFU or new disseminations.

**A3 - Virtual HIV-1/SIV infection**

Coinfection with both HIV and TB is one of the most well-known and dangerous disease outcomes[22,45]. Infection with HIV causes, among other effects, a near-complete depletion of CD4+ effector T cells[46]. HIV infects and kills different immune subpopulations preferentially - notably the CCR5+ memory cell population[46]. We assume HIV-1 and SIV virtual T-cell depletion behaves similarly[47–50]. Note that (1) HIV-1 and SIV are not identical - e.g., SIV rarely causing AIDS-like symptoms in natural hosts[47]. and (2) there is evidence that SIV-induced relapse is not identical to CD4+ T-cell depletion-induced relapse[48].

In *HostSim,* we capture co-infection with HIV via a killing term to all memory and effector cell populations. Specifically, we represent virtual HIV infection at time $t$ as an additional death term that is 0 prior to $t$ and at the time of infection, and then linearly increases to a value of $k_{HIV}$ at time $t + t_{onset}$. That is, for all CD4+ effector and memory T-cell populations, we impose

$$\frac{d}{dt}T = \cdots - \underbrace{(\Psi \cdot k_{HIV} T)}_{\text{HIV loss}}$$

where $\Psi$ is a continuous and piecewise linear scaling term that is 0.25 prior to $t$, 1 after $t + t_{onset}$, and linearly connected between those two times. We virtually infect our hosts with HIV twelve months after the end of TB treatment and assume an onset time of $t_{onset} = 2$ years. The HIV loss term introduces a new steady-state that the system reaches on a time-scale faster than the linear decrease itself, meaning that by slowly adjusting $\Psi$, we also slowly adjust the T-cell concentration as in **Figure S1.** A roughly linear drop of between 20-50% over two years was reported in the 1990s[51], and we achieve this after adapting published parameters of an older model[52] $k_{HIV} = 0.0082$.

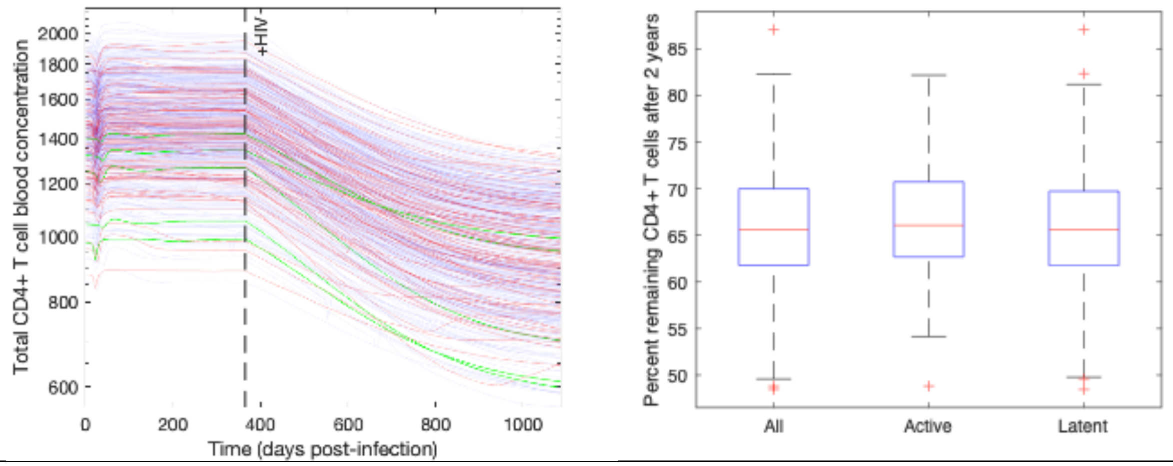


**Figure A1: Simulated CD4+ T-cell depletion during virtual HIV/SIV infection.** (Left) Blood concentration of CD4+ T cells during the first 2 years of virtual HIV infection. (Right) Percent remaining CD4+ T cells in the virtual blood compartment after 2 years of simulated HIV infection, showing a trajectory similar to that in literature[51].

### A4 - Bacterial loads versus disease state prediction method

As described in the main text, we coarse-grain our prediction of virtual host disease state by the fold-increase of CFU in that virtual host. Each virtual host has a predicted CFU trajectory based on model parameters. Parameters are in **Appendix B**, and discussed in detail in our previous work[53] as well as our website at http://malthus.micro.med.umich.edu/lab/supplements/Host-Sim-3/.

Without an accurate symptom-prediction model, we use estimate a virtual host's disease state based purely on lung bacterial load. If CFU levels increase by $R$-fold levels between days 100 (post-transient-peak) and 300 (mature infection), then we define uncontrolled infection as symptomatic disease. In *HostSim*, we use $R = 10$ to capture the 90% LTBI / 10% active TB distributions observed in human studies for decades[23,30,54,55]. However, lowering $R = 1.25$ redefines the classification, bringing the percentage active TB to 25%, closer to the 1:1 observed in cynomolgus macaques[56]. Figure S2 shows CFU counts for a single set of 500 virtual hosts, illustrating which trajectories are classified as active with NHP-like or human-like classification of active TB disease. We assume $R = 1.01$ for mice, as we assume that without confining Mtb to granulomas, any increase of bacterial burden over time is indicative of poor control.

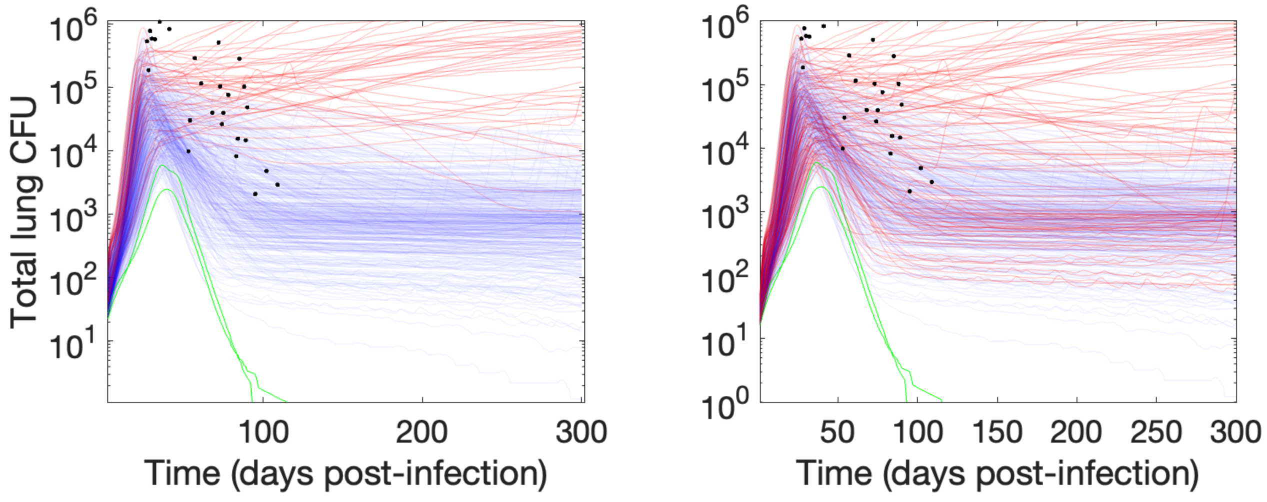


**Figure A2: Simulated CFU levels colored by predicted virtual host state.** Both panels show the same CFU trajectories of the same 500 virtual hosts based on human-like active-TB classification ($R = 10$, left) or NHP-like active-TB classification ($R = 1.25$, right). When hosts are classified at 300 days post-infection, we predict either 10% active hosts (human-like) or 25% active hosts (NHP-like).

### A5 - Dissemination

If Mtb disseminate, they can found new granulomas elsewhere in the lung[1]. This has been used as an indicator of progressive infection, and we assume that within-lung dissemination occurs in *HostSim*. We represent these as discrete events wherein a new granuloma is created within a virtual host, as in previous work with *HostSim*[53,57,58]. Disseminations in *HostSim* can be local (spatially proximal to the original infection site) or non-local (distal, possibly in other lung lobes). There are two mechanisms that contribute to a per-day probability of granuloma dissemination in *HostSim*.

*A4.1 - CFU-threshold based dissemination*

Granulomas that maintaining very high CFU levels are likely to disseminate. We assume that granulomas with CFU > $\approx 10^6$ are not well controlled and likely to progress in infection. This is measured as a per-day probability of a granuloma disseminating as

$$P(dissemination) = \lambda \cdot \frac{\text{CFU}}{\text{CFU} + 10^5},$$

where $\lambda \in [0.02, 0.04]$ for local dissemination and $\lambda \in [0.002, 0.004]$ for non-local dissemination, as calibrated previously[53].

*A4.2 - Uncontrolled growth of Mtb*

We assume that continued increase of CFU over time after the granuloma is matured (e.g., a constant increase from 100CFU to 500CFU in a years-old granuloma) is a marker for risk of dissemination. Our base estimates root from the following observations.

First is that hosts with LTBI do not disseminate frequently. As part of the dissemination of LTBI, we assume that we should expect perhaps 1 dissemination over many years. Secondly, we estimate that if CFU in a granuloma increases by >5% CFU per day, approximately 1 dissemination occurs in a month. Third, we assume that immunosuppression induces more dissemination.

We compute the probability of dissemination from this mechanism from a Weibull distribution, where

$$P(dissemination) = 1 - \exp\left(-\left(\Delta t/(L)\right)^{K(q)}\right)$$

where

$$q = \frac{(\text{previous CFU})}{(\text{current CFU})}$$

and

$$K(q) = K_{Stable} + (K_{Unstable} - K_{Stable}) \cdot \frac{(q' - 1)^2}{(q' - 1)^2 + 0.02^2};\ q' = \max(1, q),$$

and the parameters are $K_{stable} = 4.38$ and $K_{unstable} = 0.75$ for local dissemination, and $K_{stable} = 4.74$ and $K_{unstable} = 1.11$ for non-local dissemination. We chose the Weibull scale parameter $L$, to a characteristic time-scale for dissemination. We estimated $L = 90$ days as (i) dissemination events are a marker for reactivation[30], and (ii) between 50% and 75% of infliximab-induced reactivation occurs within 90 days, observed in both humans[33] and simulations[40]. For local dissemination, we compute these parameters directly from the system of equations emergent from our assumptions: $E\big(P(Dissemination\ in\ 1\ year\ with\ q = 1)\big) \approx 10^{-6}$ and $E\big(P(Dissemination\ in\ 30\ days\ with\ q = 1/1.05)\big) \approx 1$. Values for non-local dissemination are calculated with one-fifth the expected probability.

### A6 - Repeated persistence-threshold analysis using RMZE

We examine simulated rates of relapse 1 year after administering a 2-month short-course treatment of RMZE to 500 virtual hosts. We test for Mtb both at time of treatment completion and one year later by using *Virtual Mtb plate* (Methods), varying the LOD of both initial and follow-up tests from 0 to 100 CFU. We also vary whether or not these diagnostics can detect non-replicating CFU. Qualitative results are identical to those shown in HRZE (Figure A3), with the exception of a single instance of threshold-driven relapse occurring at lower LOD in virtual tests that cannot detect non-replicating bacteria.

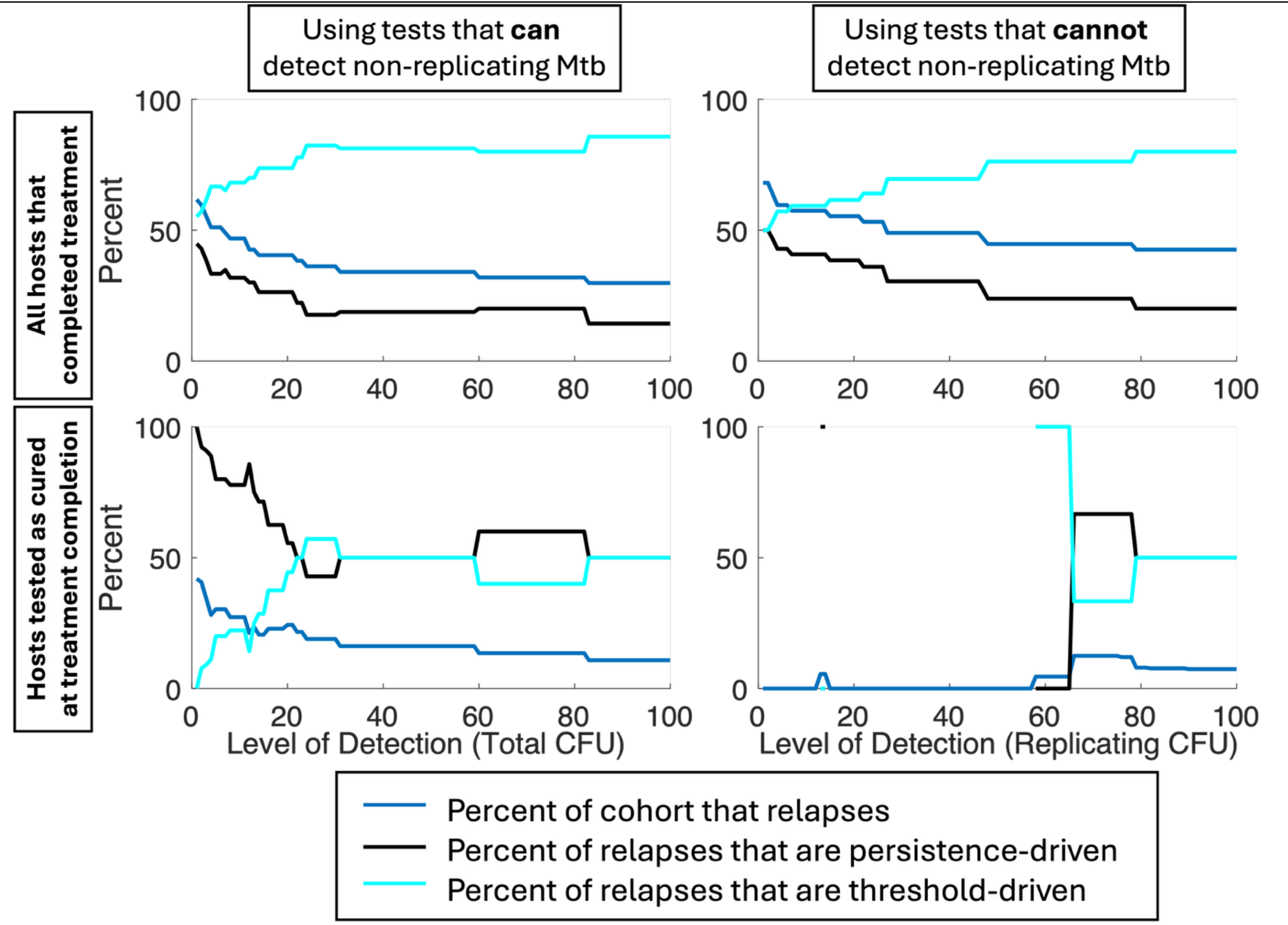


**Figure A3: Persistence versus threshold relapse rates by test LOD.** Each of our plots show how simulated relapse rates and the relative frequency of their driving mechanisms are impacted by LOD of the diagnostic test used to assess cure and/or relapse. These analyses used the *Virtual Mtb plate* test to mimic sputum culture conversion tests one year after active TB hosts complete 2 months of virtual RMZE treatment (Methods). We repeated each analysis assuming the *Virtual Mtb plate* either can (Left column) or cannot (Right column) detect non-replicating bacteria. We also repeated each analysis varying whether relapse is defined of all hosts that complete treatment (Top row) or only those that are misdiagnosed as "cured" upon treatment completion (Bottom row).
