## Appendix B for "Cure-screening predicts mechanism of post-antibiotic relapse in Tuberculosis"

### APPENDIX B - *HOSTSIM* EQUATIONS AND PARAMETERS

This document contains the equations for each compartment in *HostSim*, how they are coupled, parameter ranges, and initial conditions. *HostSim* was originally published by Louis Joslyn, Jennifer Linderman, and Denise Kirschner in the article "A virtual host model of Mycobacterium tuberculosis infection identifies early immune events as predictive of infection outcomes" in 2022 in the *Journal of Theoretical Biology.*

This technical description contains reproductions of the algorithms, equations, and supplemental materials presented in previous publications, including:

- L. Joslyn et al. "A virtual host model of Mycobacterium tuberculosis infection identifies early immune events as predictive of infection outcomes". *The Journal of Theoretical Biology.* 2022.
- C.T. Michael et al. "A framework for multi-scale intervention modeling: virtual cohorts, virtual clinical trials, and model-to-model comparisons" in *Frontiers in Systems Biology.* 2024.
- C.T. Michael et al. "Rankings of tuberculosis antibiotic treatment regimens are sensitive to spatial scale, detection limit, and initial host bacterial burden". *The Journal of Theoretical Biology.* 2025.

We have separated this document into sections for readability. In Section 1, we detail all of the equations in each granuloma compartment. In Section 2, we detail the equations in the lymph node and blood compartments. In Section 3, we present the method by which all of our compartments are coupled. In Section 4, we describe how we model dissemination - that is, how we represent one granuloma causing another to form. In Section 5, we include the entirety of our drug computation process. In Section 6, we present tables for the parameter ranges and initial condition values for our granuloma (Section 6.1), lymph node and blood (Section 6.2), and pharmacokinetic (Section 6.3) model components. In each of these sections, we included both a symbolic variable for each variable (e.g., $M_R$ or $\xi_2$) and a corresponding non-symbolic variable (e.g., MR or xi2) for convenience of navigating this document via text search.

#### 1. Granuloma equations

Each granuloma has its own distinctly-parameterized instance of the equations presented in the subsections below. Each host has a "host-scale granuloma parameter base" selected by the LHS sampling scheme using the ranges presented in Section 5.1. To capture intra-host variability, individual granulomas within a host are given parameter values normally distributed around this base host value. Terms indicated by $Pulled\,[\cdot]_g$ involve quantities pulled from the lymph node and blood equations, and is described in Section 3. Below is an index of all state variables in granulomas.

| State Variable | Non-Symbolic | Units | Description |
|---|---|---|---|
| $M_R$ | MR | count | Resting macrophages |
| $M_I$ | MI | count | Infected macrophages |
| $M_A$ | MA | count | Activated macrophages |
| $B_I$ | BI | count | Intracellular Mtb |
| $B_E$ | BE | count | Extracellular Mtb |
| $B_N$ | BN | count | Non-replicating Mtb |
| $Ca$ | CA | mass of $M\phi$ | Necrotic tissue mass |
| $Ag$ | AG | mass Ag/Mtb | Antigen mass within granulomas |
| $T_0^4$ | T40 | count | Mtb-specific primed CD4$^+$ T-cells |
| $T_1^4$ | Th1 | count | Mtb-specific Th1 CD4$^+$ T-cells |
| $T_2^4$ | Th2 | count | Mtb-specific Th2 CD4$^+$ T-cells |
| $T_{Non}^4$ | CD4Non | count | Nonspecific CD4$^+$ T-cells |
| $T_{EM}^4$ | EMCD4 | count | Mtb-specific CD4$^+$ effector memory T-cells |
| $T_0^8$ | T80 | count | Mtb-specific primed CD8$^+$ T-cells |
| $T_C$ | TC | count | Mtb-specific cytotoxic CD8$^+$ T-cells |
| $T^8$ | T8 | count | Mtb-specific effector CD8$^+$ T-cells |
| $T_{EM}^8$ | EMCD8 | count | Mtb-specific CD8$^+$ effector memory T-cells |
| $T_{Non}^8$ | CD8Non | count | Nonspecific CD8$^+$ T-cells |
| $F_\alpha$ | TNF | pg/mL | TNF-$\alpha$ concentration |
| $I_\gamma$ | IG | pg/mL | IFN-$\gamma$ concentration |
| $I_{12}$ | I12 | pg/mL | Interleukin 12 concentration |
| $I_{10}$ | I10 | pg/mL | Interleukin 10 concentration |
| $I_4$ | I4 | pg/mL | Interleukin 4 concentration |

##### 1.1. **Macrophage equations.**

$M_R$ - MR - Resting macrophage count

$$\frac{d}{dt}(M_R) = \underbrace{\alpha_{4a}\left(M_A + w_2 M_I\right)}_{\text{Macrophage-driven recruitment}} + \underbrace{Sr_{4b}\left(\frac{F_\alpha}{F_\alpha + f_8 I_{10} + s_{4b}}\right)}_{\text{TNF driven recruitment}} + \underbrace{\psi_C \psi_S \left(\sum_{j=1}^{n} R_{IC/EC \text{ for drug j}}^{FIC}\right)^{1/FIC} k_I(C_{eff}) M_I}_{\text{Drug-induced reversion from } M_I}$$
$$- \underbrace{k_2 M_R \left(\frac{B_E}{B_E + c_9}\right)}_{\text{Infection of } M_R} - \underbrace{k_3 M_R \left(\frac{B_E + wB_I + \beta F_\alpha}{B_E + wB_I + \beta F_\alpha + c_8}\right)\left(\frac{I_\gamma}{I_\gamma + f_1 I_4 + f_7 I_{10} + s_1}\right)}_{\text{Activation of macrophages}} - \underbrace{\mu_{M_R} M_R}_{\text{Natural death}}$$

$M_I$ - MI - Infected macrophages

$$\frac{d}{dt}(M_I) = \underbrace{k_2 M_R \left(\frac{B_E}{B_E + c_9}\right)}_{\text{Infection of } M_R} - \underbrace{k_{17} M_I \left(\frac{B_I^2}{B_I^2 + (NM_I)^2}\right)}_{\text{Bursting of } M_I} - \underbrace{k_{14a} M_I \left(\frac{\left(\frac{T_C + w_3 T_1^4}{M_I}\right)}{\left(\frac{T_C + w_3 T_1^4}{M_I}\right) + c_4}\right)}_{\text{T-cell driven apoptosis of } M_I}$$
$$- \underbrace{k_{14b} M_I \left(\frac{F_\alpha}{F_\alpha + f_9 I_{10} + s_{4b}}\right)}_{\text{TNF driven apoptosis of } M_I} - \underbrace{k_{52} M_I \left(\frac{\left(\frac{T_C\left(\frac{T_1^4}{T_1^4 + C_{T_1^4}}\right) + w_1 T_1^4}{M_I}\right)}{\left(\frac{T_C\left(\frac{T_1^4}{T_1^4 + C_{T_1^4}}\right) + w_1 T_1^4}{M_I}\right) + c_{52}}\right)}_{\text{Cytotoxic T-cell driven } M_I \text{ apoptosis}}$$
$$- \underbrace{\psi_C \left(\sum_{j=1}^{n} R_{IC/EC \text{ for drug j}}^{FIC}\right)^{1/FIC} k_I(C_{eff}) M_I}_{\text{Drug-induced clearance}} - \underbrace{\mu_{M_I} M_I}_{\text{Natural death}}$$

$M_A$ - MA - Activated macrophages

$$\frac{d}{dt}(M_A) = \underbrace{k_3 M_R \left(\frac{B_E + wB_I + \beta F_\alpha}{B_E + wB_I + \beta F_\alpha + c_8}\right)\left(\frac{I_\gamma}{I_\gamma + f_1 I_4 + f_7 I_{10} + s_1}\right)}_{\text{Activation of resting macrophages}} - \underbrace{k_4 M_A \left(\frac{I_{10}}{I_{10} + s_8}\right)}_{\text{Deactivation by } I_{10}} - \underbrace{\mu_{M_A} M_A}_{\text{Natural death}}$$

1.2. **Mtb and caseum equations.**

$B_I$ - BI - Intracellular bacterium count

$$\frac{d}{dt}\left(B_I\right) = \underbrace{\alpha_{19}\frac{B_I}{M_I}M_I\left(1-\frac{(B_I/M_I)}{N}\right)}_{\text{Replication}} + \underbrace{k_2\frac{N}{2}M_R\left(\frac{B_E}{B_E+c_9}\right)}_{\text{Internalization of } B_E} - \underbrace{k_{17}NM_I\left(\frac{B_I^2}{B_I^2-\left(NM_I\right)^2}\right)}_{\text{Bursting of } M_I}$$

$$- \underbrace{k_{14a}\frac{B_I}{M_I}M_I\left(\frac{\left(\frac{T_C+w_3T_1^4}{M_I}\right)}{\left(\frac{T_C+w_3T_1^4}{M_I}\right)+c_4}\right)}_{\text{T-cell driven } M_I \text{ apoptosis}} - \underbrace{k_{14b}\frac{B_I}{M_I}M_I\left(\frac{F_\alpha}{F_\alpha+f_9I_{10}+s_{4b}}\right)}_{\text{TNF driven apoptosis}}$$

$$- \underbrace{k_{52}\frac{B_I}{M_I}M_I\left(\frac{\left(\frac{T_C\left(\frac{T_1^4}{T_1^4+C_{T_1^4}}\right)+w_1T_1^4}{M_I}\right)}{\left(\frac{T_C\left(\frac{T_1^4}{T_1^4+C_{T_1^4}}\right)+w_1T_1^4}{M_I}\right)+c_{52}}\right)}_{\text{Cytotoxic T-cell driven} M_I \text{ apoptosis}} - \underbrace{\mu_{B_I}B_I}_{\text{Natural death}} - \underbrace{\mu_{M_I}\frac{B_I}{M_I}M_I}_{\text{Natural death of } M_I} - \underbrace{k_I(C_{eff})B_I}_{\substack{\text{Drug killing of } B_I \text{ based}\\ \text{on effective killing rate}}}$$

$B_E$ - BE - Extacellular bacterium count

$$\frac{d}{dt}(B_E) = \underbrace{\alpha_{20}B_E\left(1-\frac{B_E}{10^6}\right)}_{\text{Replication}} + \underbrace{(1-C_N)}_{\substack{\text{Fraction outside}\\ \text{of caseum}}}\left[\underbrace{k_{17}NM_I\left(\frac{B_I^2}{B_I^2+\left(NM_I\right)^2}\right)}_{\text{Macrophage bursting}} + \underbrace{k_{14a}N_{fracc}\frac{B_I}{M_I}M_I\left(\frac{\left(\frac{T_C+w_3T_1^4}{M_I}\right)}{\left(\frac{T_C+w_3T_1^4}{M_I}\right)+c_4}\right)}_{\text{T-cell driven apoptosis}}\right.$$

$$\left. + \underbrace{k_{14b}N_{fraca}\frac{B_I}{M_I}M_I\left(\frac{F_\alpha}{F_\alpha+f_9I_{10}+s_{4b}}\right)}_{\text{TNF driven apoptosis}} + \underbrace{\mu_{M_I}N_{fracd}\frac{B_I}{M_I}M_I}_{\text{Natural death of } M_I}\right] + \underbrace{k_{Rev}N_{Ca}\left(M_A+M_I\right)Ca\cdot B_N}_{\text{Revealing of } B_N}$$

$$- \underbrace{k_2\frac{N}{2}M_R\left(\frac{B_E}{B_E+c_9}\right)}_{\text{Internalization of } B_E} - \underbrace{k_{15}M_AB_E}_{M_A \text{ killing of } B_E} - \underbrace{k_{18}M_RB_E}_{M_R \text{ killing of } B_E} - \underbrace{\mu_{B_E}B_E}_{\text{Natural death}} - \underbrace{k_E(C_{eff})B_E}_{\substack{\text{Drug killing of } B_E \text{ based}\\ \text{on effective killing rate}}}$$

$B_N$ - BN - Nonreplicating bacterium count

$$\frac{d}{dt}\left(B_N\right) = \underbrace{C_N}_{\substack{\text{Fraction inside}\\ \text{of caseum}}}\left\{\underbrace{k_{17}M_I\left(\frac{B_I^2}{B_I^2+\left(NM_I\right)^2}\right)}_{\text{Macrophage bursting}} + \underbrace{k_{14a}N_{fracc}\frac{B_I}{M_I}M_I\left(\frac{\left(\frac{T_C+w_3T_1^4}{M_I}\right)}{\left(\frac{T_C+w_3T_1^4}{M_I}\right)+c_4}\right)}_{\text{T-cell driven apoptosis}}\right.$$

$$\left. + \underbrace{k_{14b}N_{fraca}\frac{B_I}{M_I}M_I\left(\frac{F_\alpha}{F_\alpha+f_9I_{10}+s_{4b}}\right)}_{\text{TNF driven apoptosis}} + \underbrace{\mu_{M_I}N_{fracd}\frac{B_I}{M_I}M_I}_{\text{Natural death of } M_I}\right\}$$

$$- \underbrace{\mu_{B_N}B_N}_{\text{Natural death}} - \underbrace{B_Nk_{Rev}N_{Ca}Ca\cdot M_A}_{\text{Revealing of } B_N} - \underbrace{k_N(C_{eff})B_N}_{\substack{\text{Drug killing of } B_N \text{ based}\\ \text{on effective killing rate}}}$$

$Ca$ - CA - Caseum

$$N_f \frac{d}{dt}(Ca) = \underbrace{k_{17} M_I \left( \frac{B_I^2}{B_I^2 + (N M_I)^2} \right)}_{\text{Bursting of } M_I} + \underbrace{k_{14a} M_I \left( \frac{\left(\frac{T_C + w_3 T_1}{M_I}\right)}{\left(\frac{T_C + w_3 T_1}{M_I}\right) + c_4} \right)}_{\text{T-cell mediated apoptosis of } M_I} + \underbrace{k_{14b} M_I \left( \frac{F_\alpha}{F_\alpha + f_9 I_{10} + s_{4b}} \right)}_{\text{TNF-mediated apoptosis of } M_I}$$

$$+ \underbrace{k_{52} \frac{B_I}{M_I} M_I \left( \frac{\left( \frac{T_C \left( \frac{T_1^4}{T_1^4 + C_{T_1^4}} \right) + w_1 T_1^4}{M_I} \right)}{\left( \frac{T_C \left( \frac{T_1^4}{T_1^4 + C_{T_1^4}} \right) + w_1 T_1^4}{M_I} \right) + c_{52}} \right)}_{\text{Cytotoxic T-cell mediated apoptosis of } M_I} + \underbrace{\psi_C (1 - \psi_S) \left( \sum_{j=1}^{n} R_{IC/EC \text{ for drug j}}^{FIC} \right)^{1/FIC} k_I(C_{eff}) M_I}_{M_I \text{ killed during drug clearance}}$$

$$+ \underbrace{\mu_{M_I} M_I + \mu_{M_A} M_A + \mu_{M_R} M_R \left( \frac{F_\alpha}{F_\alpha + f_9 I_{10} + s_{4b}} \right)}_{\text{Natural death of all macrophage populations}} - \underbrace{N_{Ca} Ca M_A}_{\text{Neutrophil-driven caseum clearance}}$$

$Ag$ - AG - Mtb antigen

$$\frac{d}{dt}(Ag) = \underbrace{k_{15} M_A B_E}_{M_A \text{ killing of } B_E} + \underbrace{k_{18} M_R B_E}_{M_R \text{ killing of } B_E} + \underbrace{\mu_{B_E} B_E}_{\text{Natural death of } B_E}$$

$$+ \underbrace{k_I(C_{eff}) B_I}_{\substack{\text{Drug killing of } B_I \text{ based} \\ \text{on effective killing rate}}} + \underbrace{k_E(C_{eff}) B_E}_{\substack{\text{Drug killing of } B_E \text{ based} \\ \text{on effective killing rate}}} + \underbrace{k_N(C_{eff}) B_N}_{\substack{\text{Drug killing of } B_N \text{ based} \\ \text{on effective killing rate}}}$$

$$+ \underbrace{k_{14a} \frac{B_I}{M_I} M_I \left( \frac{\left(\frac{T_C + w_3 T_1^4}{M_I}\right)}{\left(\frac{T_C + w_3 T_1^4}{M_I}\right) + c_4} \right)}_{\text{T-cell driven } M_I \text{ apoptosis}} + \underbrace{k_{14b} \frac{B_I}{M_I} M_I \left( \frac{F_\alpha}{F_\alpha + f_9 I_{10} + s_{4b}} \right)}_{\text{TNF driven aptosis}}$$

$$+ \underbrace{k_{52} \frac{B_I}{M_I} M_I \left( \frac{\left( \frac{T_C \left( \frac{T_1^4}{T_1^4 + C_{T_1^4}} \right) + w_1 T_1^4}{M_I} \right)}{\left( \frac{T_C \left( \frac{T_1^4}{T_1^4 + C_{T_1^4}} \right) + w_1 T_1^4}{M_I} \right) + c_{52}} \right)}_{B_I \text{ death by cytotoxic T-cell driven } M_I \text{ apoptosis}} + \underbrace{\mu_{B_I} B_I}_{\text{Natural Death of } B_I}$$

$$+ \underbrace{\mu_{M_I} \frac{B_I}{M_I} M_I}_{B_I \text{ death by natural death of } M_I} - \underbrace{\mu_{Ag} Ag}_{\text{Antigen clearance}}$$

1.3. **CD4$^+$ T-cells.**

$T_0^4$ - T40 - Primed CD4$^+$ T-cell count

$$\frac{d}{dt}\left(T_0^4\right) = Pulled\left[T_0^4\right]_g + \underbrace{\alpha_2 T_0^4\left(\frac{M_A}{M_A+c_{15}}\right)}_{\text{Proliferation}} - \underbrace{k_6 I_{12} T_0^4\left(\frac{I_\gamma}{I_\gamma+f_1 I_4+f_7 I_{10}+s_1}\right)}_{\text{Differentiation to } T_1^4}$$
$$-\underbrace{k_7 T_0^4\left(\frac{I_4}{I_4+f_2 I_\gamma+s_2}\right)}_{\text{Differentiation to } T_2^4} - \underbrace{\mu_{T_0} T_0}_{\text{Natural death}}$$

$T_1^4$ - Th1 - Effector CD4$^+$ T-cell count

$$\frac{d}{dt}\left(T_1^4\right) = \underbrace{k_6 I_{12} T_0^4\left(\frac{I_\gamma}{I_\gamma+f_1 I_4+f_7 I_{10}+s_1}\right)}_{\text{Differentiation from } T_0^4} + \underbrace{k_{31} T_{EM}^4 M_I\left(\frac{M_I}{M_I+3}\right)}_{\text{Differentiation from } T_{EM}^4}$$
$$-\underbrace{\mu_{T_\gamma}\left(\frac{I_\gamma}{I_\gamma+c}\right) T_1^4 M_A}_{\text{IFN}\gamma\text{ apoptosis of } T_0^1} - \underbrace{\mu_{T_1^4} T_1^4}_{\text{Natural death}}$$

$T_2^4$ - Th2 - Effector Th2 CD4$^+$ T-cell count

$$\frac{d}{dt}\left(T_2^4\right) = \underbrace{k_7 T_0^4\left(\frac{I_4}{I_4+f_2 I_\gamma+s_2}\right)}_{\text{Differentiation from } T_0^4} + \underbrace{k_{32} T_{EM}^4 M_A}_{\text{Differentiation from } T_{EM}^4} - \underbrace{\mu_{T_2^4} T_2^4}_{\text{Natural death}}$$

$T_{EM}^4$ - EMCD4 - Effector memory CD4$^+$ T-cell count

$$\frac{d}{dt}\left(T_{EM}^4\right) = Pulled\left[T_{EM}^4\right]_g - \underbrace{k_{31} T_{EM}^4 M_I\left(\frac{M_I}{M_I+3}\right)}_{\text{Differentiation to } T_1^4} - \underbrace{k_{32} T_{EM}^4 M_A}_{\text{Differentiation to } T_2^4} - \underbrace{\mu_{T_{EM}^4} T_{EM}^4}_{\text{Natural death}}$$

$T_{Non}^4$ - CD4Non - Non-cognate CD4$^+$ T-cell count

$$\frac{d}{dt}\left(T_{Non}^4\right) = Pulled\left[T_{Non}^4\right]_g - \underbrace{\mu_{T_{Non}^4} T_{Non}^4}_{\text{Natural death}}$$

#### 1.4. **CD8$^+$ T-cells.**

$T_0^8$ - T80 - Primed CD8$^+$ T-cell count:

$$\frac{d}{dt}\left(T_0^8\right) = Pulled\left[T_0^8\right]_g + \underbrace{\alpha_2 T_0^8\left(\frac{M_A}{M_A+c_{15}}\right)}_{\text{Proliforation}}$$
$$-\underbrace{k_6 I_{12} T_0^8\left(\frac{I_\gamma}{I_\gamma+f_1 I_4+f_7 I_{10}+s_1}\right)}_{\text{Differentiation to } T^8 \text{ and } T_C} - \underbrace{\mu_{T_0} T_0^8}_{\text{Natural death}}$$

$T^8$ - T8 - Effector CD8$^+$ T-cell count

$$\frac{d}{dt}\left(T^8\right) = \underbrace{(m) k_6 I_{12} T_0^8\left(\frac{I_\gamma}{I_\gamma+f_1 I_4+f_7 I_{10}+s_1}\right)}_{\text{Differentiation from } T_0^8} + \underbrace{k_{34} T_{EM}^8 M_I\left(\frac{M_I}{M_I+3}\right)}_{\text{Differentiation from } T_{EM}^8} - \underbrace{\mu_{T_{C\gamma}}\left(\frac{I_\gamma}{I_\gamma+c_c}\right) T^8 M_A}_{\text{IFN}\gamma\text{ apoptosis}} - \underbrace{\mu_{T_C} T_8}_{\text{Natural death}}$$

$T_C$ - TC - Cytotoxic CD8$^+$ T-cell count

$$\frac{d}{dt}(T_C) = \underbrace{(1-m)k_6 I_{12} T_0^8 \left(\frac{I_\gamma}{I_\gamma + f_1 I_4 + f_7 I_{10} + s_1}\right)}_{\text{Differentiation from } T_0^8} + \underbrace{k_{33} T_{EM}^8 M_I \left(\frac{M_I}{M_I + 3}\right)}_{\text{Differentiation from } T_{EM}^8} - \underbrace{\mu_{T_{C\gamma}} \left(\frac{I_\gamma}{I_\gamma + c_c}\right) T_C M_A}_{\text{IFN}\gamma \text{ apoptosis}} + \underbrace{\mu_{T_C} T_C}_{\text{Natural death}}$$

$T_{EM}^8$- EMCD8 - Effector Memory CD8$^+$ T-cell count

$$\frac{d}{dt}\left(T_{EM}^8\right) = Pulled\left[T_{EM}^8\right]_g - \underbrace{k_{33} T_{EM}^8 M_I \left(\frac{M_I}{M_I+3}\right)}_{\text{Differentiation to } T_C} - \underbrace{k_{34} T_{EM}^8 M_I \left(\frac{M_I}{M_I+3}\right)}_{\text{Differentiation to } T^8} - \underbrace{\mu_{T_{EM}^8} T_{EM}^8}_{\text{Natural death}}$$

$T_{Non}^8$ -CD8Non - Non-cognate CD8$^+$ T-cell count

$$\frac{d}{dt}\left(T_{Non}^8\right) = Pulled\left[T_{Non}^8\right]_g - \underbrace{\mu_{T_{Non}^8} T_{Non}^8}_{\text{Natural death}}$$

1.5. **Cytokines.**

$F_\alpha$ - TNF - concentration of Tumor Necrosis Factor $\alpha$

$$\frac{d}{dt}(F_\alpha) = \underbrace{\alpha_{30} M_I}_{\text{Production by } M_I} + \underbrace{\alpha_{31} M_A \left(\frac{I_\gamma + \beta_2 (B_E + w B_I)}{I_\gamma + \beta_2 (B_E + w B_I) + f_1 I_4 + f_7 I_{10} + s_{10}}\right)}_{\text{IFN-IL mediated production of TNF}\alpha \text{ by } M_A}$$
$$+ \underbrace{\alpha_{32} T_1^4}_{\text{Production by } T_1^4} + \underbrace{\alpha_{33}\left(\frac{T_C + T^8}{2m}\right)}_{\text{Production by CD8}^+ \text{ T-cells}} - \underbrace{\mu_{F_\alpha} F_\alpha}_{\text{Clearance}}$$

$I_\gamma$ - IG - concentration of Interferon-$\gamma$

$$\frac{d}{dt}(I_\gamma) = \underbrace{s_g \left(\frac{B_E + w B_I}{B_E + w B_I + c_{10}}\right)\left(\frac{I_{12}}{I_{12} + s_7}\right)}_{\text{Other}} + \underbrace{\alpha_{5a} T_1^4 \left(\frac{M_A}{M_A + c_{5a}}\right)}_{\text{Production by } T_1^4} + \underbrace{\alpha_{5b} T^8 \left(\frac{M_A}{M_A + c_{5b}}\right)}_{\text{Production by } T^8} + \underbrace{\alpha_{5c} M_I}_{\text{Production by } M_I}$$
$$+ \underbrace{\alpha_7 T_0^4 \left(\frac{I_{12}}{I_{12} + f_4 I_{10} + s_4}\right) + \alpha_7 T_0^8 \left(\frac{I_{12}}{I_{12} + f_4 I_{10} + s_4}\right)}_{\text{Interleukin-mediated production by primed T-cells}} - \underbrace{\mu_{I_\gamma} I_\gamma}_{\text{Clearance}}$$

$I_{12}$ - I12 - concentration of Interleukin-12

$$\frac{d}{dt}(I_{12}) = \underbrace{s_{12}\left(\frac{B_E + w B_I}{B_E + w B_I + c_{230}}\right)}_{\text{Other}} + \underbrace{\alpha_{23} M_R \left(\frac{B_E + w B_I}{B_E + w B_I + c_{23}}\right)}_{\text{Production by } M_R} + \underbrace{\alpha_8 M_A \left(\frac{s}{s + I_{10}}\right)}_{\text{Production by } M_A} - \underbrace{\mu_{I_{12}} I_{12}}_{\text{Clearance}}$$

$I_{10}$ - I10 - concentration of Interleukin-10

$$\frac{d}{dt}(I_{10}) = \underbrace{\delta_7 (M_I + M_A)\left(\frac{s_6}{I_{10} + f_6 I_\gamma + s_6}\right)}_{\text{IFN-IL mediated production by macrophages}} + \underbrace{\alpha_{16} T_1^4 + \alpha_{17} T_2^4}_{\text{Production by CD4}^+ \text{ T-cells}} + \underbrace{\alpha_{18}\left(\frac{T_C + T^8}{2m}\right)}_{\text{Production by CD8}^+ \text{ T-cells}} - \underbrace{\mu_{I_{10}} I_{10}}_{\text{Clearance}}$$

$I_4$ - I4 - concentration of Interleukin-4

$$\frac{d}{dt}(I_4) = \underbrace{\alpha_{11} T_0^4}_{\text{Production by } T_0^4} + \underbrace{\alpha_{12} T_2^4}_{\text{Production by } T_2^4} - \underbrace{\mu_{I_4} I_4}_{\text{Clearance}}$$

#### 2. Lymph node and blood equations

The lymph node compartment will clonally expand and differentiate effector T cells to be sent to blood. Cells in blood can be recruited to granulomas based on their cytokine levels, which is represented by terms indicated by $Pulled\,[\cdot]_g$ (Section 3). We select each parameter in the following equations using the LHS sampling scheme using the ranges presented in Section 5.2. Below is an index of all state variables in the compartment representing lymph nodes and blood.

| Variable | Non-symbolic | Unit | Variable description |
|---|---|---|---|
| | **Lymph Node** | | |
| $APC$ | APC | count | Antigen-presenting cells in LN |
| $N_4^{LN}$ | LnN4 | count | Mtb-specific naive CD4$^+$ T-cell count in LN |
| $P_4^{LN}$ | LnP4 | count | Mtb-specific precursor CD4$^+$ T-cell count in LN |
| $E_4^{LN}$ | LnE4 | count | Mtb-specific effector CD4$^+$ T-cell count in LN |
| $CM_4^{LN}$ | LnCM4 | count | Mtb-specific central memory CD4$^+$ T-cell count in LN |
| $EM_4^{LN}$ | LnEM4 | count | Mtb-specific effector memory CD4$^+$ T-cell count in LN |
| $N_{Non,4}^{LN}$ | LnN4Non | count | Nonspecific naive CD4$^+$ T-cell count in LN |
| $CM_{Non,4}^{LN}$ | LnCM4Non | count | Nonspecific central memory CD4$^+$ T-cell count in LN |
| $N_8^{LN}$ | LnN8 | count | Mtb-specific naive CD8$^+$ T-cell count in LN |
| $P_8^{LN}$ | LnP8 | count | Mtb-specific precursor CD8$^+$ T-cell count in LN |
| $E_8^{LN}$ | LnE8 | count | Mtb-specific effector CD8$^+$ T-cell count in LN |
| $CM_8^{LN}$ | LnCM8 | count | Mtb-specific central memory CD8$^+$ T-cell count in LN |
| $EM_8^{LN}$ | LnEM8 | count | Mtb-specific effector memory CD8$^+$ T-cell count in LN |
| $N_{Non,8}^{LN}$ | LnN8Non | count | Nonspecific naive CD8$^+$ T-cell count in LN |
| $CM_{Non,8}^{LN}$ | LnCM8Non | count | Nonspecific central memory CD8$^+$ T-cell count in LN |
| | **Blood** | | |
| $N_4^B$ | BlN4 | pg/mL | Mtb-specific naive CD4$^+$ T-cell count in blood |
| $E_4^B$ | BlE4 | pg/mL | Mtb-specific effector CD4$^+$ T-cell count in blood |
| $CM_4^B$ | BlCM4 | pg/mL | Mtb-specific central memory CD4$^+$ T-cell count in blood |
| $EM_4^B$ | BlEM4 | pg/mL | Mtb-specific effector memory CD4$^+$ T-cell count in blood |
| $N_{Non,4}^B$ | BlN4Non | pg/mL | Nonspecific naive CD4$^+$ T-cell count in blood |
| $E_{Non,4}^B$ | BlE4Non | pg/mL | Nonspecific effector CD4$^+$ T-cell count in blood |
| $CM_{Non,4}^B$ | BlCM4Non | pg/mL | Nonspecific central memory CD4$^+$ T-cell count in blood |
| $EM_{Non,4}^B$ | BlEM4Non | pg/mL | Nonspecific effector memory CD4$^+$ T-cell count in blood |
| $N_8^B$ | BlN8 | pg/mL | Mtb-specific naive CD8$^+$ T-cell count in blood |
| $E_8^B$ | BlE8 | pg/mL | Mtb-specific effector CD8$^+$ T-cell count in blood |
| $CM_8^B$ | BlCM8 | pg/mL | Mtb-specific central memory CD8$^+$ T-cell count in blood |
| $EM_8^B$ | BlEM8 | pg/mL | Mtb-specific effector memory CD8$^+$ T-cell count in blood |
| $N_{Non,8}^B$ | BlN8Non | pg/mL | Nonspecific naive CD8$^+$ T-cell count in blood |
| $E_{Non,8}^B$ | BlE8Non | pg/mL | Nonspecific effector CD8$^+$ T-cell count in blood |
| $CM_{Non,8}^B$ | BlCM8Non | pg/mL | Nonspecific central memory CD8$^+$ T-cell count in blood |
| $EM_{Non,8}^B$ | BlEM8Non | pg/mL | Nonspecific effector memory CD8$^+$ T-cell count in blood |

2.1. **Antigen Presenting Cells.**

$APC$ - APC - Antigen presenting cells. Received APCs are sent from granulomas, which is detailed in Section 3.

$$\frac{d}{dt}(APC) = -\mu_5 APC + \sum_{i\in\{\text{Granulomas}\}} \text{Received from Granuloma } i$$

2.2. **CD4$^+$ T-cells in lymph nodes.**

$N_4^{LN}$ - LnN4 - CD4$^+$ Mtb-specific naive T-cell count

$$\frac{d}{dt}(N_4^{LN}) = \alpha\left[\underbrace{k_1 N_4^B\left(\frac{APC}{APC+hs_1}\right)}_{\text{Cytokine-driveen recruitment}} + \underbrace{\xi_1 N_4^B}_{\text{LN Influx}}\right] - \underbrace{\xi_2 N_4^{LN}}_{\text{LN Efflux}} - \underbrace{k_2 N_4^{LN} APC}_{\text{Differentiation into } P_4^{LN}}$$

$P_4^{LN}$ - LnP4 - CD4$^+$ Mtb-specific precursor T-cell count

$$\frac{d}{dt}(P_4^{LN}) = \underbrace{k_2 N_4^{LN} APC}_{\text{Differentiation from } N_4^{LN}} + \underbrace{k_3 CM_4^{LN} APC}_{\text{Differentiation from } CM_4^{LN}} + \underbrace{k_4 P_4^{LN}\left(1-\frac{P_4^{LN}}{\rho_1}\right)\left(\frac{APC}{APC+hs_4}\right)}_{\text{Proliferation}}$$
$$- \underbrace{k_5 P_4^{LN}\left(\frac{APC}{APC+hs_5}\right)}_{\text{Differention into } E_4^{LN}} - \underbrace{k_6 P_4^{LN}\left(1-\frac{APC}{APC+hs_5}\right)}_{\text{Differentiation into } CM_4^{LN}} - \underbrace{\mu_6 P_4^{LN}}_{\text{Natural death}}$$

$E_4^{LN}$- LnE4 - CD4$^+$ Mtb-specific effector T-cell count

$$\frac{d}{dt}(E_4^{LN}) = \underbrace{k_5 P_4^{LN}\left(\frac{APC}{APC+hs_5}\right)}_{\text{Differention from } P_4^{LN}} - \underbrace{\xi_3 E_4^{LN}}_{\text{LN Efflux}} - \underbrace{k_7 E_4^{LN}}_{\text{Differentiation into } EM_4^{LN}}$$

$CM_4^{LN}$- LnCM4 - CD4$^+$ Mtb-specific central memory T-cell count

$$\frac{d}{dt}(CM_4^{LN}) = \alpha\left[\underbrace{k_8 CM_4^B\left(\frac{APC}{APC+hs_8}\right)}_{\text{Cytokine-mediated recruitment}} + \underbrace{\xi_4 CM_4^B}_{\text{LN Influx}}\right] + \underbrace{k_6 P_4^{LN}\left(1-\frac{APC}{APC+hs_5}\right)}_{\text{Differentiation from } P_4^{LN}}$$
$$- \underbrace{k_3 CM_4^{LN} APC}_{\text{Differentiation to } P_4^{LN}} - \underbrace{\xi_5 CM_4^{LN}}_{\text{LN Efflux}}$$

$EM_4^{LN}$ - LnEM4 - CD4$^+$ Mtb-specific effector memory T-cell count

$$\frac{d}{dt}(EM_4^{LN}) = \underbrace{k_7 E_4^{LN}}_{\text{Differentiation from } E_4^{LN}} - \underbrace{\xi_6 EM_4^{LN}}_{\text{LN Efflux}}$$

$N_{Non,4}^{LN}$ - LnN4Non - Non-cognate naive CD4$^+$ T-cell count

$$\frac{d}{dt}(N_{Non,4}^{LN}) = \alpha\left(\underbrace{k_1 N_{Non,4}^B\left(\frac{APC}{APC+hs_1}\right)}_{\text{Cytokine-mediated recruitment}} + \underbrace{\xi_1 N_{Non,4}^B}_{\text{LN Influx}}\right) - \underbrace{\xi_2 N_{Non,4}^{LN}}_{\text{LN Efflux}}$$

$CM_{Non,4}^{LN}$ - LnCM4Non - Non-cognate central memory CD4$^+$ T-cell count

$$\frac{d}{dt}\left(CM_{Non,4}^{LN}\right) = \alpha\left(\underbrace{k_8 CM_{Non,4}^{B}\left(\frac{APC}{APC+hs_8}\right)}_{\text{Cytokine-mediated recruitment}} + \underbrace{\xi_4 CM_{Non,4}^{B}}_{\text{LN Influx}}\right) - \underbrace{\xi_5 CM_{Non,4}^{LN}}_{\text{LN Efflux}}$$

##### 2.3. CD8$^+$ T-cells in lymph nodes.

$N_8^{LN}$ - LnN8 - Mtb-specific naive CD8$^+$ T-cell count

$$\frac{d}{dt}\left(N_8^{LN}\right) = \alpha\left(\underbrace{k_{10} N_8^{B}\left(\frac{APC}{APC+hs_{10}}\right)}_{\text{Cytokine-mediated recruitment}} + \underbrace{\xi_7 N_8^{B}}_{\text{LN Influx}}\right) - \underbrace{\xi_8 N_8^{LN}}_{\text{LN Efflux}} - \underbrace{k_{11} N_8^{LN} APC\left(\frac{E_4^{LN}+w_{P_4}P_4^{LN}}{E_4^{LN}+w_{P_4}P_4^{LN}+hs_{11}}\right)}_{\text{Priming mediated by CD4-related cytokines}}$$

$P_8^{LN}$ - LnP8 - Mtb-specific precursor CD8$^+$ T-cell count

$$\frac{d}{dt}\left(P_8^{LN}\right) = \underbrace{k_{11} N_8^{LN} APC\left(\frac{E_4^{LN}+w_{P_4}P_4^{LN}}{E_4^{LN}+w_{P_4}P_4^{LN}+hs_{11}}\right)}_{\text{Priming mediated by CD4-related cytokines}} + \underbrace{k_{12} CM_8^{LN} APC}_{\text{Differentiation from } CM_8^{LN}} + \underbrace{k_{13} P_8^{LN}\left(1-\frac{P_8^{LN}}{\rho_1}\right)\left(\frac{APC}{APC+hs_{13}}\right)}_{\text{Cytokine-mediated proliferation}} - \underbrace{k_{15} P_8^{LN}\left(1-\left\{\frac{APC}{APC+hs_{14}}\right\}\right)}_{\text{Differentiation to } CM_8^{LN}} - \underbrace{k_{14} P_8^{LN}\left\{\frac{APC}{APC+hs_{14}}\right\}}_{\text{Differentiation to } E_8^{LN}} - \underbrace{\mu_7 P_8^{LN}}_{\text{Natural death}}$$

$E_8^{LN}$ - LnE8 - Mtb-specific effector CD8$^+$ T-cell count

$$\frac{d}{dt}\left(E_8^{LN}\right) = \underbrace{k_{14} P_8^{LN}\left\{\frac{APC}{APC+hs_{14}}\right\}}_{\text{Differentiation from } P_8^{LN}} - \underbrace{\xi_9 E_8^{LN}}_{\text{LN Efflux}} - \underbrace{k_{16} E_8^{LN}}_{\text{Differentiation to } EM_8^{LN}}$$

$CM_8^{LN}$ - LnCM8 - Mtb-specific central memory CD8$^+$ T-cell count

$$\frac{d}{dt}\left(CM_8^{LN}\right) = \alpha\left[\underbrace{k_{17} CM_8^{B}\left(\frac{APC}{APC+hs_{17}}\right)}_{\text{Cytokine-mediated recruitment}} + \underbrace{\xi_{10} CM_8^{B}}_{\text{LN Influx}}\right] + \underbrace{k_{15} P_8^{LN}\left(1-\left\{\frac{APC}{APC+hs_{14}}\right\}\right)}_{\text{Differentiation from } P_8^{LN}} - \underbrace{k_{12} CM_8^{LN} APC}_{\text{Cytokine-mediated differentiation to } P_8^{LN}} - \underbrace{\xi_{11} CM_8^{LN}}_{\text{LN Efflux}}$$

$EM_8^{LN}$ - LnEM8 - Mtb-specific effector memory CD8$^+$ T-cell count

$$\frac{d}{dt}\left(EM_8^{LN}\right) = \underbrace{k_{16} E_8^{LN}}_{\text{Differentiation from } E_8^{LN}} - \underbrace{\xi_{12} EM_8^{LN}}_{\text{LN Efflux}}$$

$N_{Non,8}^{LN}$ - LnN8Non - Non-cognate Naive CD8$^+$ T-cell count

$$\frac{d}{dt}\left(N_{Non,8}^{LN}\right) = \alpha\left[\underbrace{k_{10}N_{Non,8}^{B}\left(\frac{APC}{APC+hs_{10}}\right)}_{\text{Cytokine-mediated recruitment}} + \underbrace{\xi_7 N_{Non,8}^{B}}_{\text{LN Influx}}\right] - \underbrace{\xi_8 N_{Non,8}^{LN}}_{\text{LN Efflux}}$$

$CM_{Non,8}^{LN}$ - LnCM8Non - Non-cognate central memory CD8$^+$ T-cell count

$$\frac{d}{dt}\left(C_{Non,8}^{LN}\right) = \alpha\left[\underbrace{k_{17}CM_{Non,8}^{B}\left(\frac{APC}{APC+hs_{17}}\right)}_{\text{Cytokine-mediated recruitment}} + \underbrace{\xi_{10} CM_{Non,8}^{B}}_{\text{LN Influx}}\right] - \underbrace{\xi_{11} CM_{Non,8}^{LN}}_{\text{LN Efflux}}$$

2.4. **CD4$^+$ T-cells in blood.**

$N_4^B$ - BlN4 - Mtb-specific naive CD4$^+$ T-cell concentration

$$\frac{d}{dt}\left(N_4^B\right) = \underbrace{\lambda S_{N_4}}_{\text{Production in thymus}} + \underbrace{\alpha^{-1}\xi_2 N_4^{LN}}_{\text{LN Efflux}} - \underbrace{k_1 N_4^B\left(\frac{APC}{APC+hs_1}\right)}_{\text{Cytokine-mediated recruitment}} - \underbrace{\xi_1 N_4^B}_{\text{LN Influx}} - \underbrace{\mu_8 N_4^B}_{\text{Natural dath}}$$

$E_4^B$ - BlE4 - Mtb-specific effector CD4$^+$ T-cell concentration

$$\frac{d}{dt}\left(E_4^B\right) = \underbrace{\alpha^{-1}\xi_3 E_4^{LN}}_{\text{LN Efflux}} - \underbrace{\mu_1 E_4^B}_{\text{Natural death}} - \sum_{g\in Granulomas} Pulled\left[E_4^B\right]_g$$

$CM_4^B$ - BlCM4 - Mtb-specific central memory CD4$^+$ T-cell concentration

$$\frac{d}{dt}\left(CM_4^B\right) = \underbrace{\alpha^{-1}\xi_5 CM_4^{LN}}_{\text{LN Efflux}} - \underbrace{\xi_4 CM_4^B}_{\text{LN Influx}} - \underbrace{k_8 CM_4^B\left(\frac{APC}{APC+hs_8}\right)}_{\text{Cytokine-mediated recruitment}}$$

$EM_4^B$ - BlEM4 - Mtb-specific effector memory CD4$^+$ T-cell concentration

$$\frac{d}{dt}\left(EM_4^B\right) = \underbrace{\alpha^{-1}\xi_6 EM_4^{LN}}_{\text{LN Efflux}} - \underbrace{\mu_2 EM_4^B}_{\text{Natural death}} - \sum_{g\in Granulomas} Pulled\left[EM_4^B\right]_g$$

$N_{Non,4}^B$ - BlN4Non - Non-cognate naive CD4$^+$ T-cell concentration

$$\frac{d}{dt}\left(N_{Non,4}^B\right) = \underbrace{(1-\lambda)\, s_{N_4}}_{\text{Production in thymus}} + \underbrace{\alpha^{-1}\xi_2 N_{Non,4}^{LN}}_{\text{LN Efflux}} - \underbrace{k_1 N_{Non,4}^B\left(\frac{APC}{APC+hs_1}\right)}_{\text{Cytokine-mediated recruitment}} - \underbrace{\xi_1 N_{Non,4}^B}_{\text{LN Influx}} - \underbrace{\mu_8 N_{Non,4}^B}_{\text{Natural death}}$$

$E_{Non,4}^B$ - BlE4Non - Non-cognate effector CD4$^+$ T-cell concentration

$$\frac{d}{dt}\left(E_{Non,4}^B\right) = \underbrace{s_{E_{Non,4}}}_{\text{Production in thymus}} - \underbrace{\mu_4 E_{Non,4}^B}_{\text{Natural death}} - \sum_{g\in Granulomas} Pulled\left[E_{Non,4}^B\right]_g$$

$CM_{Non,4}^B$ - BlCM4Non - Non-cognate central memory CD4$^+$ T-cell concentration

$$\frac{d}{dt}\left(CM_{Non,4}^B\right) = \underbrace{\alpha^{-1}\xi_5 CM_{Non,4}^{LN}}_{\text{LN Efflux}} - \underbrace{\xi_4 CM_{Non,4}^B}_{\text{LN Influx}} - \underbrace{k_8 CM_{Non,4}^B\left(\frac{APC}{APC+hs_8}\right)}_{\text{Cytokine-mediated recruitment}}$$

$EM_{Non,4}^B$ - BlEM4Non - Non-cognate effector memory CD4$^+$ T-cell concentration

$$\frac{d}{dt}\left(EM^B_{Non,4}\right) = \underbrace{s_{EM_{Non,4}}}_{\text{Source from thymus}} - \underbrace{\mu_2 EM^B_{Non,4}}_{\text{Natural death}} - \sum_{g \in Granulomas} Pulled\left[EM^B_{Non,4}\right]_g$$

2.5. **CD8$^+$ T-cells in blood.**

$N^B_8$ - BlN8 Mtb-specific naive CD8$^+$ T-cell concentration

$$\frac{d}{dt}\left(N^B_8\right) = \underbrace{\lambda s_{N_8}}_{\text{Source from thymus}} + \underbrace{\alpha^{-1}\xi_8 N^{LN}_8}_{\text{LN efflux}} - \underbrace{k_{10} N^B_8 \left(\frac{APC}{APC + hs_{10}}\right)}_{\text{Cytokine-mediated recruitment}} - \underbrace{\xi_7 N^B_8}_{\text{LN Influx}} - \underbrace{\mu_9 N^B_8}_{\text{Natural death}}$$

$E^B_8$ - BlE8 - Mtb-specific effector CD8$^+$ T-cell concentration

$$\frac{d}{dt}\left(E^B_8\right) = \underbrace{\alpha^{-1}\xi_9 E^{LN}_8}_{\text{LN Efflux}} - \underbrace{\mu_3 E^B_8}_{\text{Natural death}} - \sum_{g \in Granulomas} Pulled\left[E^B_8\right]_g$$

$CM^B_8$ - BlCM8 - Mtb-specific central memory CD8$^+$ T-cell concentration

$$\frac{d}{dt}\left(CM^B_8\right) = \underbrace{\alpha^{-1}\xi_{11} CM^{LN}_8}_{\text{LN Efflux}} - \underbrace{\xi_{10} CM^B_8}_{\text{LN Influx}} - \underbrace{k_{17} CM^B_8 \left(\frac{APC}{APC + hs_{17}}\right)}_{\text{Cytokine-mediated recruitment}}$$

$EM^B_8$ - BlEM8 - Mtb-specific Effector memory CD8$^+$ T-cell concentration

$$\frac{d}{dt}\left(EM^B_8\right) = \underbrace{\alpha^{-1}\xi_{12} EM^{LN}_8}_{\text{LN Efflux}} - \underbrace{\mu_4 EM^B_8}_{\text{Natural death}} - \sum_{g \in Granulomas} Pulled\left[EM^B_8\right]_g$$

$N^B_{Non,8}$ - BlN8Non - Non-cognate naive CD8$^+$ T-cell concentration

$$\frac{d}{dt}\left(N^B_{Non,8}\right) = \underbrace{(1-\lambda)\, s_{N_8}}_{\text{Source from thymus}} + \underbrace{\alpha^{-1}\xi_8 N^{LN}_{Non,8}}_{\text{LN Efflux}}$$
$$- \underbrace{k_{10} N^B_{Non,8} \left(\frac{APC}{APC + hs_{10}}\right)}_{\text{Cytokine-mediated recruitment}} - \underbrace{\xi_7 N^B_{Non,8}}_{\text{LN Influx}} - \underbrace{\mu_9 N^B_{Non,8}}_{\text{Natural death}}$$

$E^B_{Non,8}$ - BlE8Non - Non-cognate effector CD8$^+$ T-cell concentration

$$\frac{d}{dt}\left(E^B_{Non,8}\right) = \underbrace{s_{E_{Non,8}}}_{\text{Source from thymus}} - \underbrace{\mu_3 E^B_{Non,8}}_{\text{Natural death}} - \sum_{g \in Granulomas} Pulled\left[E^B_{Non,8}\right]_g$$

$CM^B_{Non,8}$ - BlCM8Non - Non-cognate central memory CD8$^+$ T-cell concentration

$$\frac{d}{dt}\left(CM^B_{Non,8}\right) = \underbrace{\alpha^{-1}\xi_{11} CM^{LN}_{Non,8}}_{\text{LN Efflux}} - \underbrace{\xi_{10} CM^B_{Non,8}}_{\text{LN Influx}} - \underbrace{k_{17} CM^B_{Non,8} \left(\frac{APC}{APC + hs_{17}}\right)}_{\text{Cytokine-mediated recruitment}}$$

$EM^B_{Non,8}$ - BlEM8Non - Non-cognate effector memory CD8$^+$ T-cell concentration

$$\frac{d}{dt}\left(EM^B_{Non,8}\right) = \underbrace{s_{EM_{Non,8}}}_{\text{Source from thymus}} - \underbrace{\mu_4 EM^B_{Non,8}}_{\text{Natural death}} - \sum_{g \in Granulomas} Pulled\left[EM^B_{Non,8}\right]_g$$

#### 3. Coupling granuloma equations to lymph node and blood equations

**3.1. Migration of APCs from granulomas to lymph nodes.** Each granuloma begins to send a portion of its infected macrophages to the lymph node compartment. We use the sum of infected macrophages $M_I$ and antigen $Ag$ as a proxy for the number of dendritic cells that take Mtb to lung draining lymph nodes. Each granuloma sends APCs to the lymph nodes after time $\tau_s$ under the assumption that granulomas do not immediately begin to send dendritic cells to the lymphatic system. We compute APCs sent by granuloma $g$ to the lymph node as:

$$APC \text{ Sent by Granuloma } g \text{ at time } t \text{ post-founding} = \begin{cases} 0 & \text{if } t < \tau_s \\ A_{frac} \cdot (Ag + M_I) & \text{otherwise.} \end{cases}$$

**3.2. Granulomas request T-cells, pulling them from the blood compartment.** In *HostSim*, a given host begins with 13 granulomas. As infection progresses, each granuloma will begin to attempt to recruit six types of T-cells from the blood compartment: $T_0^4, T_0^8, T_{EM}^4, T_{EM}^8, T_{Non}^4$, and $T_{Non}^8$. At each time step, each granuloma will independently request a number of T-cells given below:

- Requests for $T_0^4$ :

$$Requested\left[T_0^4\right] = \underbrace{\alpha_{1a}\left(w_2 M_I + M_A\right)}_{\text{Macrophage-mediated request}} + \underbrace{Sr_{1b}\left(\frac{F_\alpha}{F_\alpha + f_8 I_{10} + s_{4b2}}\right)}_{\text{TNF mediated recruitment}}$$

- Requests for $T_0^8$ :

$$Requested\left[T_0^8\right] = \underbrace{\alpha_{1a}\left(M_A + w_2 M_I\right)}_{\text{Macrophage-mediated recruitment}} + \underbrace{Sr_{1b}\left(\frac{F_\alpha}{F_\alpha + f_8 I_{10} + s_{4b2}}\right)}_{\text{TNF mediated recruitment}}$$

- Requests for $T_{EM}^4$:

$$Requested\left[T_{EM}^4\right] = \underbrace{Sr_{4EM}\left(\frac{F_\alpha}{F_\alpha + hs_{4EM}}\right)}_{\text{TNF driven recruitment}}$$

- Requests for $T_{EM}^8$:

$$Requested\left[T_{EM}^8\right] = \underbrace{Sr_{8EM}\left(\frac{F_\alpha}{F_\alpha + hs_{8EM}}\right)}_{\text{TNF-mediated recruitment}}$$

- Requests for $T_{Non}^4$:

$$Requested\left[T_{Non}^4\right] = \underbrace{Sr_{4Non}\left(\frac{F_\alpha}{F_\alpha + hs_{4Non}}\right)}_{\text{TNF driven recruitment}}$$

- Requests for $T_{Non}^8$:

$$Requested\left[T_{Non}^8\right] = \underbrace{Sr_{8Non}\left(\frac{F_\alpha}{F_\alpha + hs_{8Non}}\right)}_{\text{TNF driven recruitment}}$$

The blood's response to these requests is to provide lung each granuloma with T-cells. Since we model the T-cell subpopulations in more granularity in the lymph node and blood compartments, we associate the following populations between the terminology of the lymph node and blood equations; and the granuloma equations:

(1) Granuloma requests for $T_0^4$ pull from Lymph/blood concentration $E_4^B$.
(2) Granuloma requests for $T_0^8$ pull from Lymph/blood concentration $E_8^B$.
(3) Granuloma requests for $T_{EM}^4$ pull from Lymph/blood concentration $EM_4^B$.
(4) Granuloma requests for $T_{EM}^8$ pull from Lymph/blood concentration $EM_8^B$.
(5) Granuloma requests for $T_{Non}^4$ pull from both Lymph/blood concentrations $E_{Non,4}^B$ and $EM_{Non,4}^B$.
(6) Granuloma requests for $T_{Non}^8$ pull from both Lymph/blood concentrations $E_{Non,8}^B$ and $EM_{Non,8}^B$.

For $T_0^4, T_0^8, T_{EM}^4$, and $T_{EM}^8$, the amount that is pulled from the blood to provide to granuloma $g$ is given by:

$$\underbrace{Pulled\,[T^*]_g}_{T\text{ - Cell count}} = \alpha \cdot \underbrace{Pulled\,[E^*]_g}_{E\text{ - Concentration}}$$
$$= \begin{cases} Requested\,[T^*]_g & \text{if } \sum_{\text{Granulomas } j} Requested\,[T^*]_j < \alpha \cdot E^* \\ Relative\ Request\,[T^*]_g & \text{otherwise.} \end{cases}$$

Here, if there are not enough T-cells in the blood to meet the amount requested by the lung granulomas, available T-cells will be divided among the requesting granulomas by using the following relative request. For granuloma $g$, its relative request is

$$Relative\ Request\,[T^*]_g = \frac{\alpha \cdot E^*}{\sum_{\text{All granulomas}j} Request\,[T^*]_j}.$$

For $T_{Non}^4$ and $T_{Non}^8$, we combine their two associated blood concentrations. For example,

$$Pulled\left[T_{Non}^4\right]_g = \alpha\left(Pulled\left[E_{Non,4}^B\right]_g + Pulled\left[EM_{Non,4}^B\right]_g\right)$$
$$= \begin{cases} Requested\left[T_{Non}^4\right]_g & \text{if } \sum_{\text{Granulomas } j} Requested\left[T_{Non}^4\right]_j < \alpha\left(E_{Non,4}^B + EM_{Non,4}^B\right) \\ Relative\ Request\left[T_{Non}^4\right]_g & \text{otherwise} \end{cases}.$$

If the blood has excess T-cells ($E_{Non,4}^B$ and $EM_{Non,4}^B$ in this case), the weight of their contribution to the granulomas is based on their current relative population:

$$\alpha \cdot Pulled\left[E_{Non,4}^B\right]_g = Requested\left[T_{Non}^4\right]_g \frac{E_{Non,4}^B}{\left(E_{Non,4}^B + EM_{Non,4}^B\right)};$$
$$\alpha \cdot Pulled\left[EM_{Non,4}^B\right]_g = Requested\left[T_{Non}^4\right]_g \frac{EM_{Non,4}^B}{\left(E_{Non,4}^B + EM_{Non,4}^B\right)}.$$

If there are not enough T-cells for the request, the relative requests are calculated similarly and the blood concentration is depleted:

$$\alpha \cdot Pulled\left[E_{Non,4}^B\right]_g = \frac{Requested\left[T_{Non}^4\right]_g}{\sum_{\text{Granulomas } j} Requested\,[T_{Non}^4]_j} \frac{E_{Non,4}^B}{\left(E_{Non,4}^B + EM_{Non,4}^B\right)};$$
$$\alpha \cdot Pulled\left[EM_{Non,4}^B\right]_g = \frac{Requested\left[T_{Non}^4\right]_g}{\sum_{\text{Granulomas } j} Requested\,[T_{Non}^4]_j} \frac{EM_{Non,4}^B}{\left(E_{Non,4}^B + EM_{Non,4}^B\right)}.$$

Pulled concentrations for $T_{Non}^8$ are calculated in the same way.

#### 4. Dissemination

In *HostSim*, nonsterile granulomas have a chance to disseminate. Each granuloma begins with a local dissemination probability adjustment $L_t$ and nonlocal dissemination probability adjustment $N_t$ when the granuloma is created (either primary at $t = 0$, or at some later time). At each timestep, a dissemination event is determined randomly determining:

$$\{\text{Local dissemination occurs at time } t\} = (L_t + U[0,1]) > 1$$

and

$$\{\text{Nonlocal dissemination occurs at time } t\} = (N_t + U[0,1]) > 1$$

At each time step, the probability adjustment of local dissemination increases:

$$L_t = L_{t-1} + \lambda_{local}\left(\frac{[B_I + B_E]\,(t)}{[B_I + B_E]\,(t) + k_{local}}\right).$$

Similarly, nonlocal dissemination probability adjustment builds as

$$N_t = N_{t-1} + \lambda_{nonlocal} \left( \frac{[B_I + B_E](t)}{[B_I + B_E](t) + k_{nonlocal}} \right).$$

A new locally-disseminated granuloma is created with its parameters selected to be normally distributed around the parameter values of the primary granuloma that it disseminated from with $\sigma = 10\%$ of the width of the parameter's range (log-transformed, if the parameter is log-uniformly sampled). This is because of the assumption that nearby lung tissue conditions give rise to similar dynamics. Nonlocally disseminated granlomas are given parameters uniformly sampled out of the ranges presented in Section 5.

Disseminated granulomas begin with one infected macrophage and an average number of intracellular bacteria from its parent ($B_I/M_I$ from the primary granuloma that seeded it).

#### 5. Antibiotic treatment model

Our antibiotic treatment model assumes that antibiotic treatment begins on a pre-specified day $t_i > 0$ where 0 is the day of pulmonary infection. For a virtual host undergoing virtual treatment, there is a pre-specified regimen of 1 or more antibiotics. This model is separated in to two major components: pharmacokinetics and pharmacodynamics. Broadly, we assume that there are no drug-drug interactions that affect pharmacokinetics, and the processes detailed for computing drug concentrations are repeated for each drug independently.

5.1. **Pharmacokinetics model.** All concentrations are assumed to be uniformly 0 for $t < t_i$. Concentrations and drug masses are tracked independently for each drug in the regimen. Parameters governing each drug's concentration in time are given in Section 6.

5.1.1. *Plasma and lung pharmacokinetics.* "*Dose*" may be considered to be a $\delta$-function impuse of drug once per given dosing interval. The magnitude of the impulse "i.e. dose amount" is a parameter measured in mg/kg. Note that a primary output of this set of equations is the concentration of drug in healthy lung tissue $C_{lung} = D_L/V_L$, which is updated once every hour of simulated time. Also, the variable $\delta_{iT}$ is set to either 1 or 0 based on however many transition compartments the specific drug is using.

$D_{T_1}$ - DT1 - Dose concentration compartment 1:

$$\frac{d(D_{T_1})}{dt} = Dose - \underbrace{\delta_{2T} k_d D_{T_1}}_{\text{To } T_2} - \underbrace{\delta_{1T} k_d D_{T_1}}_{\text{Direct to } P}$$

$D_{T_2}$ - DT2 - Dose concentration 2:

$$\frac{d(D_{T_2})}{dt} = \underbrace{\delta_{2T} k_d D_{T_1}}_{\text{From } T_1} - \underbrace{\delta_{2T} k_d D_{T_2}}_{\text{To } P}$$

$D_{PT}$ - DPT - Drug concentration in peripheral tissue:

$$\frac{d(D_{PT})}{dt} = \underbrace{Q\left(\frac{D_P}{V_P} - \frac{D_{PT}}{V_{PT}}\right)}_{\text{Drug uptake into peripheral tissue}}$$

$D_P$ - DP - Drug concentration in plasma:

$$\frac{d(D_P)}{dt} = \underbrace{\delta_{1T} k_d D_{T_1}}_{\text{Direct from } T_1} + \underbrace{\delta_{2T} k_d D_{T_2}}_{\text{From } T_2} - \underbrace{\mu_{D_P} \cdot \frac{D_P}{V_P}}_{\text{Clearance in plasma}} - \underbrace{Q\left(\frac{D_P}{V_P} - \frac{D_{PT}}{V_{PT}}\right)}_{\text{Drug uptake into peripheral tissue}}$$

$D_L$ - DL - Drug concentration in the lung:

$$\frac{d}{dt}(D_L) = \underbrace{Q\left(P_L \frac{D_P}{V_P} - \frac{D_L}{V_L}\right)}_{\text{From plasma}} - \underbrace{\mu_D D_L}_{\text{Lung clearance}}$$

5.1.2. *Granuloma pharmacokinetics.* Note that the quantity $C_{lung}$ is considered as a constant in these equations, and is updated hourly from the whole-lung pharmacokinetics. Also note that the parameter $\mu_D$ is inherited from the virtual host's whole-lung parameter $\mu_D$, as we assume that the pharmacokinetics of the viable cellular area are the same as the lung tissue that the granuloma is found in.

$D_a$ - DA - Drug in cellular area, measured in mg

$$\frac{d}{dt}(D_a) = \underbrace{Q_a SA_g \left(P_a C_{lung} - \frac{D_a}{V_a}\right)}_{\text{Lung to Cellular}} - \underbrace{Q_c SA_c \left(P_c \frac{D_a}{V_a} - \frac{D_c}{V_c}\right)}_{\text{Cellular to caseum}} - \underbrace{\mu_D D_a}_{\text{Clearance}}$$

$D_C$ - DC - Drug in caseum, measured in mg

$$\frac{d}{dt}(D_c) = \underbrace{Q_c SA_c \left(P_c \frac{D_a}{V_a} - \frac{D_c}{V_c}\right)}_{\text{Cellular to caseum}} - \underbrace{\mu_D D_c}_{\text{Clearance}}$$

5.1.3. *Effective caseum concentration.* We assume that antibiotics in caseum are effective proportional to their concentration in the caseum. However, Instead of using $C_c = D_c/V_c$ directly, we consider a modified value

$$C_c^* = C_c \cdot \beta_{drug}^{r_c \cdot \tau / r_0}$$

where $r_c$ is the radius of caseum and $\tau$ is a fitting parameter. This means that if $r_c \ll 1$, then $C_c^* \approx C_c$ (suggesting it's easier to treat smaller granulomas)

We choose the base $\beta$ - the fold-reduction of effective concentration of the drug buried in $r_0$ =0.4mm of caseum. This gives us the **distribution-like** function $\beta_{Drug}^{\tau}$. The lab of Veronique Dartois

https://www.ncbi.nlm.nih.gov/pmc/articles/PMC5607931/ listed fraction unbound (fu%), where 100% means the drug flows freely into caseum and 0% means the drug is entirely bound by caseum macromolecules. For each drug, we choose the range of $\beta$ to be the confidence interval of the range of fu% in that work.

#### 5.2. **Pharmacodynamics model.**

5.2.1. *Minimum inhibitory concentration and active drugs.* We assume that drugs will not affect the system unless they are present in sufficient concentration. Drugs that are present in sufficient concentration, *active drugs*, are used in all calculations in the following subsections. Minimum inhibitory concentrations for each drug are given in the following table.

| Antibiotic | Minimum inhibitory concentration (mg/L) |
|---|---|
| Isoniazid | 0.004 |
| Rifampacin | 0.002 |
| Pyrazinamide | 3 |
| Ethambutol | 0.06 |
| Bedaquiline | 0.006 |
| Pretomanid | 0.008 |
| Linezolid | 0.05 |
| Moxifloxacin | 1 |

5.2.2. *Pharmacodynamics modeling.* We find KillBI/BE/BN $\delta_I, \delta_E, \delta_N$ based on the concentrations of active drugs from the PK model. Note if $V_a, V_c = 0$, the Hill fraction is considered to be 1). Units of $D_x$ are in mg, and $V_a$ are calculated in $L$. The equations to determine kill-rates in the absence of drug-drug interaction are

$$k_I = E_{B_I,max}\left(\frac{\left(\frac{D_a}{V_a}\right)^{h_{B_I}}}{\left(\frac{D_a}{V_a}\right)^{h_{B_I}} + c_{50,B_I}^{h_{B_I}}}\right);$$

$$k_E = E_{B_E,max}\left(\frac{\left(\frac{D_a}{V_a}\right)^{h_{B_E}}}{\left(\frac{D_a}{V_a}\right)^{h_{B_E}} + c_{50,B_E}^{h_{B_E}}}\right);$$

$$\text{and } \delta_N = E_{B_N,max}\left(\frac{\left(\frac{D_c}{V_c}\right)^{h_{B_N}}}{\left(\frac{D_c}{V_c}\right)^{h_{B_N}} + c_{50,B_N}^{h_{B_N}}}\right).$$

5.2.3. *Modeling drug-drug interactions.* We incorporate drug-drug interactions into our model regimens when multiple drugs are present within a physiological compartment (e.g., cellular area or caseum). Briefly, we adjust effective concentrations of active drugs using fractional inhibitory concentrations (FICs) of drug combinations predicted by an *in silico* tool, INDIGO-MTB (inferring drug interactions using chemogenomics and orthology optimized for Mtb), as we have done previously. INDIGO-MTB is a machine learning based tool that uses known drug interactions and drug transcriptomics data to predict unknown drug interactions in the form of FICs. FIC values lower or higher than 1 means the drugs are synergistic or antagonistic, respectively, whereas an FIC value of 1 means the drugs do not interact, i.e., they are additive.

We model drug interaction by converting concentrations of each active drug $i$ ($C_i$) to equipotent concentrations of drug $i_{max}$, the drug with highest maximal killing rate (i.e., highest $E_{max}$). To do that, we calculate the adjusted concentration of $i$ ($C_{i,adj}$), which is the concentration of $i_{max}$ that would kill Mtb with the same rate as drug $i$ with concentration $C_i$:

$$C_{i,adj} = \left(\frac{C_{i_{max},50}^{h_{i_{max}}} C_i^{h_i}}{\frac{E_{max,i_{max}}}{E_{max,i}}\left(C_i^{h_i} + C_{i,50}^{h_i}\right) + C_i^{h_i}}\right)^{1/h_{i_{max}}}$$

where $C_{i_{max},50}$ and $C_{i,50}$ are the concentration of $i_{max}$ and $i$ at which half maximal killing is achieved, respectively, $E_{max,i_{max}}$ and $E_{max,i}$ are the maximal killing rate constants of drug $i_{max}$ and drug $i$, respectively, and $h_{i_{max}}$ and $h_i$ are the Hill coefficients of drug $i_{max}$ and drug $i$, respectively. Once we calculate the adjusted concentrations of all drugs in a compartment, we determine the effective concentration ($C_{eff}$) with the following equation:

$$C_{eff} = \left(\sum_{i=1}^{n} C_{i,adj}^{FIC}\right)^{1/FIC}$$

where $C_{i,adj}$ is the adjusted concentration of drug i, n is the number of drugs in a compartment and FIC is the FIC value predicted for these n drugs by INDIGO-MTB. Then, we calculate the effective killing rate $k$ by using $C_{eff}$ and Hill parameters of $i_{max}$ ($E_{max,i_{max}}$, $C_{50,i_{max}}$, $h_{i_{max}}$):

$$k\left(C_{eff}\right) = E_{max,i_{max}} \frac{C_{eff}^{h_{i_{max}}}}{C_{eff}^{h_{i_{max}}} + C_{i_{max},50}^{h_m}}.$$

5.2.4. *Clearance of infected macrophages by drugs.* We add the following terms to macrophage and caseum dynamics. Parameters $\psi_i$ represent granuloma-driven factors contributing to heterogeneity macrophage clearance. We assume that the heterogeneity of macrophage clearance driven by drug PK/PD variability is accounted for by other factors in the drug-induced macrophage clearance terms in Section 1. Notably, those factors depend upon $R_{IC/EC}$, the ratio of intracellular to extracellular drugs. The values we use are in the table below.

| Antibiotic | $R_{IC/EC}$ |
|---|---|
| Isoniazid | 1.3 |
| Rifampacin | 6.5 |
| Pyrazinamide | 0.6 |
| Ethambutol | 10.3 |
| Bedaquiline | 482 |
| Pretomanid | 6.6 |
| Linezolid | 3.6 |
| Moxifloxacin | 17.5 |

#### 6. Parameter Ranges

In all parameter tables, distributions listed as 'u' are uniformly sampled within the given range, and 'l' are log-uniformly sampled from within the given range. Note that the rate constants $k_i$ in the granuloma compartment and LN/blood compartment are distinct. Certain parameters indicated with * have values computed dynamically by other model components. Also note that for space, we refer to "rates" as shorthand for "rate constants" in all following tables.

##### 6.1. **Granuloma parameter ranges.**

| **Parameter** | **Non-symbolic** | **Units** | **Granuloma parameter description** | **Minimum Value** | **Maximum Value** | **Distribution** |
|---|---|---|---|---|---|---|
| $\psi_C$ | psiC | - | Antibiotic clearance rate of $M_I$ | 0.0001 | 0.001 | u |
| $\psi_S$ | psiS | - | Survival rate of cleared $M_I$ | 0 | 1 | u |
| $C_{eff}$ | Ceff | - | Effective concentration of antibiotic | * | * | - |
| $\alpha_{4a}$ | alpha4a | 1/day | Macrophage-driven $M_R$ recruitment rate | 0.7246 | 0.9097 | u |
| $\beta$ | Beta | 1/pg | Scaling factor for $F_\alpha$ for $M_R$ activation | 8652476.9683 | 11390938.3080 | u |
| $w$ | w | - | Contribution of $B_I$ to $M_R$ recruitment | 0.2601 | 0.3632 | u |
| $w_2$ | w2 | - | Contribution of $M_I$ to $M_R$ recruitment | 0.1 | 0.5 | u |
| $w_3$ | w3 | - | Contribution of $T_1^4$ to $M_I$ apoptosis | 0.9572 | 1.1725 | u |
| $Sr_{4b}$ | Sr4b | 1/day | $F_\alpha$ recruitment rate of $M_R$ | 400.3165 | 961.7117 | u |
| $f_8$ | f8 | - | Ratio adjustment of $I_{10}/F_\alpha$ on $M_R$ recruitment | 0.02568 | 0.9748 | u |
| $f_9$ | f9 | - | Ratio adjustment of $I_{10}/F_\alpha$ for apoptosis rates | 0.1 | 1.0 | u |
| $s_{4b}$ | s4b | pg/mL | Half saturation of $F_\alpha$ on $M_R$ recruitment | 43.2587 | 432.1618 | l |
| $k_4$ | k4 | 1/day | $M_A$ decativation by $I_{10}$ | 0.06138 | 0.1816 | u |
| $s_8$ | s8 | pg/mL | Half saturation of $I_{10}$ on $M_A$ deactivation | 18.2688 | 156.2867 | l |
| $k_2$ | k2 | 1/day | $M_R$ infection rate | 0.1126 | 0.4900 | u |
| $c_9$ | c9 | count | Half saturation of $B_E$ on $M_R$ infection | 1305.8725 | 8590.4539 | u |
| $k_3$ | k3 | 1/day | $M_R$ activation rate | 0.1080 | 0.5012 | u |
| $f_1$ | f1 | - | Ratio adjustment of $I_4/I_\gamma$ on $M_R$ activation | 132.009929 | 477.7016 | u |
| $s_1$ | s1 | pg/mL | Half saturation of $I_\gamma$ dependent $M_R$ activation | 2.6104 | 120.4874 | l |
| $c_8$ | c8 | count | Half saturation of $B_E$ and $B_I$ on $M_R$ activation | 345.3760 | 681.6706 | u |
| $\mu_{M_R}$ | muMR | 1/day | $M_R$ death rate | 0.004432 | 0.005695 | u |
| $k_{17}$ | k17 | 1/day | Maximum rate of $M_I$ bursting | 0.03598 | 0.3017 | u |
| $N$ | N | count | Carrying capacity of Mtb in one $M_I$ | 5 | 40 | u |

| Parameter | Non-symbolic | Units | Granuloma parameter description | Minimum Value | Maximum Value | Distribution |
|---|---|---|---|---|---|---|
| $k_{14a}$ | k14a | 1/day | T-cell induced apoptosis rate of $M_I$ | 0.2726 | 0.45 | u |
| $c_4$ | c4 | count | Half saturation of $T_1^4/M_I$ on T-cell apoptosis | 905.8568 | 9420.9084 | l |
| $k_{14b}$ | k14b | 1/day | $F_\alpha$-induced apoptosis rate | 0.2230 | 0.4355 | u |
| $k_{52}$ | k52 | 1/day | Cytotoxic killing rate of $M_I$ | 0.08592 | 0.3519 | u |
| $w_1$ | w1 | - | Contribution of $T_1^4$ to cytotoxic killing | 0.2192 | 0.7413 | u |
| $c_{52}$ | c52 | count | Half-saturation of $T_C$ on cytotoxic $M_I$ killing | 2588.9284 | 8088.5625 | u |
| $C_{T_1^4}$ | cT1 | count | Half saturation of $T_1^4$ on cytotoxic killing | 80.6446 | 300.4901 | u |
| $\mu_{M_I}$ | muMI | 1/day | $M_I$ death rate | 0.002965 | 0.003799 | u |
| $\mu_{M_A}$ | muMA | 1/day | $M_A$ death rate | 0.05 | 0.1 | u |
| $\alpha_{1a}$ | alpha1a | 1/day | Macrophage request rate of $T_0^4$ | 0.1769 | 0.4336 | u |
| $Sr_{1b}$ | Sr1b | 1/day | $F_\alpha$ dependent $T_0^4$ recruitment | 23767.7947 | 55329.7803 | u |
| $s_{4b2}$ | s4b2 | pg/mL | Half saturation of $F_\alpha$ dependent $T_0^4$ recruitment | 462.3252 | 1057.4677 | l |
| $\alpha_2$ | alpha2 | 1/day | Maximum growth rate of $T_0^4$ | 0.2444 | 0.9666 | u |
| $c_{15}$ | c15 | count | Half saturation of $M_A$ proliferation of $T_0^4$ | 531.8574 | 9574.4150 | u |
| $k_6$ | k6 | 1/day | Maximum $T_0^4$ to $T_1^4$ differentiation rate | 0.01372 | 0.09727 | u |
| $f_7$ | f7 | - | Effect of $I_{10}$ on $I_\gamma$ induced $T_0^4 \to T_1^4$ differentiation | 5.7185 | 45.2677 | u |
| $k_7$ | k7 | 1/day | Maximum rate of $T_0^4 \to T_2^4$ differentiation | 0.2437 | 0.6525 | u |
| $f_2$ | f2 | - | Ratio adjustment of $I_\gamma/I_4$ on $T_0^4 \to T_2^4$ differentiation | 0.1938 | 0.4207 | u |
| $s_2$ | s2 | pg/mL | Half saturation of $I_4$ for $T_0^4 \to T_2^4$ differentiation | 100.3818 | 964.1090 | l |
| $\mu_{T_0}$ | muT0 | 1/day | $CD3^+$ primed T-cell death rate | 0.1957 | 0.2474 | u |
| $m$ | m | - | Fraction of differentiating $T_0^8$ that become $T^8$ (not $T_C$) | 0.1064 | 0.9043 | u |
| $\mu_{T_g}$ | muTg | 1/day | $T_C$ death rate | 0.005308 | 0.01990 | l |
| $c$ | c | pg/mL | Half saturation of $I_\gamma$ on $T_1^4$ apoptosis | 462.9043 | 3161.3955 | u |
| $\mu_{T_1^4}$ | muT1 | 1/day | $T_1^4$ death rate | 0.2848 | 0.3698 | u |
| $\mu_{T_2^4}$ | muT2 | 1/day | $T_2^4$ death rate | 0.2913 | 0.3678 | u |
| $\mu_{T_{C\gamma}}$ | muTCg | 1/day | $I_\gamma$ driven apoptosis rate of $T_C$ and $T^8$ | 0.006021 | 0.09506 | l |
| $c_c$ | cc | pg/mL | Half saturation of $I_\gamma$ on $T_C$ and $T^8$ apoptosis | 368.2888 | 9634.1767 | u |
| $\mu_{T_C}$ | muTC | 1/day | $T_C$ death rate | 0.2570 | 0.3325 | u |
| $\alpha_{30}$ | alpha30 | 1/day | Rate of $F_\alpha$ production by $M_I$ | 0.04521 | 0.09694 | u |

| Parameter | Non-symbolic | Units | Granuloma parameter description | Minimum Value | Maximum Value | Distribution |
|---|---|---|---|---|---|---|
| $\alpha_{31}$ | alpha31 | 1/day | Rate of $F_\alpha$ production by $M_A$ | 0.03113 | 0.09610 | u |
| $\beta_2$ | beta2 | 1/pg | Scaling factor of Mtb on $F_\alpha$ production by $M_A$ | 10944.8638 | 13022.3032 | u |
| $s_{10}$ | s10 | pg/mL | Half saturation of $I_\gamma$ on $F_\alpha$ production by $M_A$ | 102.8084 | 1500.4010 | u |
| $\alpha_{32}$ | alpha32 | pg/(mL*day) | $F_\alpha$ production rate by $T_1^4$ | 0.1843 | 0.3102 | u |
| $\alpha_{33}$ | alpha33 | pg/(mL*day) | $F_\alpha$ production rate by $T^8$ and $T_C$ | 0.1633 | 0.2997 | u |
| $\mu_{TNF}$ | muTNF | 1/day | $F_\alpha$ clearance rate | 0.9334 | 1.5101 | u |
| $s_g$ | sg | pg/(mL*day) | $I_\gamma$ production from other sources (e.g. dendritic cells) | 846.3332 | 9484.1532 | l |
| $c_{10}$ | c10 | count | Half saturation of $I_\gamma$ production from other sources | 315332.8289 | 536623.8464 | u |
| $s_7$ | s7 | pg/mL | Half saturation of $I_{12}$ on $I_\gamma$ production from other sources | 891.2493 | 1151.7672 | u |
| $\alpha_{5a}$ | alpha5a | pg/day | Rate of $I_\gamma$ production by $T_1^4$ | 0.5443 | 0.8698 | u |
| $c_{5a}$ | c5a | count | Half saturation of $M_A$ on $I_\gamma$ production by $T_1^4$ | 303.7365 | 687.08808 | u |
| $\alpha_{5b}$ | alpha5b | pg/day | $I_\gamma$ production by $T^8$ | 1.1445 | 15.1796 | u |
| $\alpha_{5c}$ | alpha5c | pg/day | $I_\gamma$ production by $M_I$ | 0.5592 | 0.9184 | u |
| $c_{5b}$ | c5b | count | Half saturation of $M_A$ on $I_\gamma$ production by $T^8$ | 235.8238 | 846.5511 | u |
| $\alpha_7$ | alpha7 | pg/day | $I_\gamma$ production by $T_0^4$ | 0.08541 | 0.3098 | u |
| $f_4$ | f4 | - | Adjustment of $I_{10}/I_{12}$ on $I_\gamma$ production | 1.2966 | 1.6715 | u |
| $s_4$ | s4 | pg/mL | Half saturation of I12 on $I_\gamma$ | 321.7448 | 865.2110 | u |
| $\mu_{I_\gamma}$ | muIG | 1/day | $I_\gamma$ clearance rate | 6.4475 | 12.6934 | u |
| $\alpha_{23}$ | alpha23 | pg/day | Rate of $I_{12}$ production by $M_R$ | 0.003485 | 0.004652 | u |
| $c_{23}$ | c23 | count | Half saturation of Mtb on $I_{12}$ production by $M_R$ | 157.08445 | 525.4198 | u |
| $\alpha_8$ | alpha8 | pg/day | Rate of production of $I_{12}$ by $M_A$ | 0.3764 | 0.8612 | u |
| $s_{12}$ | s12 | pg/day | Rate of production of $I_{12}$ by dendritic cells | 2361.1941 | 4060.7654 | u |
| $c_{230}$ | c230 | count | Half-saturation of Mtb in $I_{12}$ production by DCs | 365.6847 | 761.7284 | u |
| $\mu_{I_{12}}$ | muI12 | 1/day | $I_{12}$ clearance rate | 0.9319 | 1.2420 | u |
| $s$ | s | pg/mL | $I_{10}$ effect on $I_{12}$ production by $M_A$ | 191.7181 | 694.3849 | u |
| $\delta_7$ | delta7 | pg/day | $I_{10}$ production by $M_A$ | 0.1106 | 0.6169 | u |
| $s_6$ | s6 | pg/mL | Half saturation of $I_{10}$ on self | 587.3834 | 858.9373 | u |
| $f_6$ | f6 | - | Ratio adjustment of $I_\gamma$ on $I_{10}$ | 0.3011 | 0.3889 | u |
| $\alpha_{16}$ | alpha16 | pg/day | $I_{10}$ production by $T_1^4$ | 0.4280 | 0.6693 | u |

| Parameter | Non-symbolic | Units | Granuloma parameter description | Minimum Value | Maximum Value | Distribution |
|---|---|---|---|---|---|---|
| $\alpha_{17}$ | alpha17 | pg/day | $I_{10}$ production by $T_2^4$ | 0.4149 | 0.4786 | u |
| $\alpha_{18}$ | alpha18 | pg/day | $I_{10}$ production by $T_C$ and $T^8$ | 0.5184 | 0.6648 | u |
| $\mu_{I_{10}}$ | muI10 | 1/day | $I_{10}$ clearance rate | 0.5904 | 4.007931 | u |
| $\alpha_{11}$ | alpha11 | pg/day | $I_4$ production by $T_0^4$ | 0.02944 | 0.06404 | u |
| $\alpha_{12}$ | alpha12 | pg/day | $I_4$ production by $T_2^4$ | 0.02110 | 0.06407 | u |
| $\mu_{I_4}$ | muI4 | 1/day | $I_4$ clearance rate | 2.3700 | 3.08806 | u |
| $\alpha_{19}$ | alpha19 | 1/day | $B_I$ growth rate | 0.1081 | 1.6183 | l |
| $\alpha_{20}$ | alpha20 | 1/day | $B_E$ growth rate | 0.1927 | 0.6209 | u |
| $N_{fracc}$ | Nfracc | - | Fraction of surviving $B_I$ released by T-cell $M_I$ apop. | 0.3074 | 0.7893 | u |
| $N_{fraca}$ | Nfraca | - | Fraction of surviving $B_I$ released by TNF $M_I$ apop. | 0.2680 | 0.5658 | u |
| $k_{15}$ | k15 | 1/day | $B_E$ killing rate by $M_A$ | 0.04058 | 0.1173 | u |
| $k_{18}$ | k18 | 1/day | $B_E$ killing rate by $M_R$ | 0.0003264 | 0.0005505 | u |
| $N_{fracd}$ | Nfracd | - | Fraction of surviving $B_I$ released by $M_I$ natural death. | 0.0008721 | 0.001098 | u |
| $\mu_{B_I}$ | muBI | 1/day | $B_I$ death rate | 0.00002973 | 0.00004521 | u |
| $\mu_{B_E}$ | muBE | 1/day | $B_E$ death rate | 0.000000002053 | 0.000000003702 | u |
| $Sr_{4Non}$ | Sr4Non | 1/day | Rate of TNF-driven $T_{Non}^4$ recruitment | 158.6388 | 450.4727 | u |
| $hs_{4Non}$ | hs4Non | pg/mL | Half saturation of TNF in $T_{Non}^4$ recruitment | 6.1706 | 49.6606 | u |
| $\mu_{4Non}$ | mui4Non | 1/day | Death rate of $T_{Non}^4$ | 0.2631 | 0.3579 | u |
| $Sr_{8Non}$ | Sr8Non | 1/day | Rate of TNF-driven $T_{Non}^8$ recruitment | 156.08771 | 450.8067 | u |
| $hs_{8Non}$ | hs8Non | pg/day | Half saturation of TNF in $T_{Non}^8$ recruitment | 4.3815 | 40.4821 | u |
| $\mu_{T_{Non}^8}$ | mui8Non | 1/day | Death rate of $T_{Non}^8$ | 0.2644 | 0.3543 | u |
| $Sr_{4EM}$ | Sr4EM | 1/day | Rate of TNF-driven $T_{EM}^4$ recruitment | 5145.1195 | 10500.04020 | u |
| $hs_{4EM}$ | hs4EM | pg/mL | Half saturation of TNF in $T_{EM}^4$ recruitment | 100.0 | 200.0 | u |
| $\mu_{T_{EM}^4}$ | mui4EM | 1/day | Death rate of $T_{EM}^4$ | 0.001368 | 0.002737 | u |
| $k_{31}$ | k31 | 1/day | Differentiation rate of $T_{EM}^4 \to T_1^4$ | 0.05592 | 0.1 | l |
| $k_{32}$ | k32 | 1/day | Differentiation rate of $T_{EM}^4 \to T_2^4$ | 0.0006898 | 0.001086 | u |
| $Sr_{EM^8}$ | Sr8EM | 1/day | Rate of TNF-driven $T_{EM}^8$ recruitment | 60.2088 | 115.9625 | u |
| $hs_{EM^8}$ | hs8EM | pg/mL | Half saturation of TNF in $T_{EM}^8$ recruitment | 100 | 300 | u |
| $\mu_{T_{EM}^8}$ | mui8EM | 1/day | Death rate of $T_{EM}^8$ | 0.001368 | 0.002737 | u |

| Parameter | Non-symbolic | Units | Granuloma parameter description | Minimum Value | Maximum Value | Distribution |
|---|---|---|---|---|---|---|
| $k_{33}$ | k33 | 1/day | Differentiation rate of $T^8_{EM} \rightarrow T_C$ | 0.06339 | 0.1 | l |
| $k34$ | k34 | 1/day | Differentiation rate of $T^8_{EM} \rightarrow T^8$ | 0.05329 | 0.1 | l |
| $\tau_s$ | taus | days | Time that granuloma begins sending APCs to LN | 4 | 17 | u |
| $A_{frac}$ | APCpercent | - | Fraction of $M_I$ and $Ag$ that gets sent as APCs | 0.1723 | 0.2893 | u |
| $k_{local}$ | localDissemCFUHalf | count | Half-saturation of Mtb for local dissemination | 10000 | 50000 | u |
| $\lambda_{local}$ | localDissemLambda | 1/day | Maximum rate of local dissemination | 0.0005 | 0.009 | l |
| $k_{nonlocal}$ | nonLocalDissemCFUHalf | count | Half-saturation of Mtb for nonlocal dissemination | 5223.04303 | 10608.01033 | u |
| $\lambda_{nonlocal}$ | nonLocalDissemLambda | 1/day | Maximum rate of nonlocal dissemination | 0.0001 | 0.002 | u |
| $\mu_{B_N}$ | muBN | 1/day | Death rate of $B_N$ | 0 | 0 | u |
| $C_N$ | CN | - | Fraction of released $B_I$ that become $B_N$ | 0.0003125 | 0.08172 | l |
| $\mu_{Ag}$ | muAg | 1/day | Decay rate of Mtb antigen | 0.003985 | 0.02686 | u |
| $N_f$ | Nf | M$\phi$/caseum | Macrophage biomass that becomes caseum | 13 | 18 | u |
| $N_{Ca}$ | Nca | 1/(day*caseum) | Clearance rate of caseum by neutrophils | 0.000001173 | 0.000008621 | u |

| **Initial Condition** | **Non-Symbolic** | **Units** | **Granuloma initial condition description** | **Min initial value** | **Max initial value** | **Distribution** |
|---|---|---|---|---|---|---|
| $M_R(0)$ | MR | count | Resting macrophages | 1.0 | 5.0 | u |
| $M_I(0)$ | MI | count | Infected macrophages | 1.0 | 1.0 | u |
| $M_A(0)$ | MA | count | Activated macrophages | 1.0 | 1.0 | u |
| $B_I(0)$ | BI | count | Intracellular Mtb | 1.0 | 1.0 | u |
| $B_E(0)$ | BE | count | Extracellular Mtb | 0 | 0 | u |
| $B_N(0)$ | BN | count | Non-replicating Mtb | 0 | 0 | u |
| $T_0^4(0)$ | T40 | count | Mtb-specific primed $CD4^+$ T-cells | 0 | 0 | u |
| $T_1^4(0)$ | Th1 | count | Mtb-specific Th1 $CD4^+$ T-cells | 0 | 0 | u |
| $T_{Non}^4(0)$ | CD4Non | count | Nonspecific $CD4^+$ T-cells | 0 | 0 | u |
| $T_2^4(0)$ | Th2 | count | Mtb-specific Th2 $CD4^+$ T-cells | 0 | 0 | u |
| $T_{EM}^8(0)$ | EMCD4 | count | Mtb-specific $CD4^+$ effector memory T-cells | 0 | 0 | u |
| $T_0^8(0)$ | T80 | count | Mtb-specific primed $CD8^+$ T-cells | 0 | 0 | u |
| $T_C(0)$ | TC | count | Mtb-specific cytotoxic $CD8^+$ T-cells | 0 | 0 | u |
| $T^8(0)$ | T8 | count | Mtb-specific effector $CD8^+$ T-cells | 0 | 0 | u |
| $T_{EM}^8(0)$ | EMCD8 | count | Mtb-specific $CD8^+$ effector memory T-cells | 0 | 0 | u |
| $T_{Non}^8(0)$ | CD8Non | count | Nonspecific $CD8^+$ T-cells | 0 | 0 | u |
| $F_\alpha(0)$ | TNF | pg/mL | TNF-$\alpha$ concentration | 0 | 0 | u |
| $I_\gamma(0)$ | IG | pg/mL | IFN-$\gamma$ concentration | 0 | 0 | u |
| $I_{12}(0)$ | I12 | pg/mL | Interleukin 12 concentration | 0 | 0 | u |
| $I_{10}(0)$ | I10 | pg/mL | Interleukin 10 concentration | 0 | 0 | u |
| $I_4(0)$ | I4 | pg/mL | Interleukin 4 concentration | 0 | 0 | u |
| $Ca(0)$ | CA | mass of $M\phi$ | Necrotic tissue mass | 0 | 0 | u |
| $Ag(0)$ | AG | mass Ag/Mtb | Antigen mass within granulomas | 0 | 0 | u |

6.2. **Lymph Node and Blood parameter ranges.** Note that the rate constants $k_i$ in the granuloma compartment and LN/blood compartment are different.

| **Parameter** | **non-symbolic** | **Units** | **LN/blood parameter description** | **Lower range** | **Upper range** | **Distribution** |
|---|---|---|---|---|---|---|
| $\alpha$ | alpha | $\mu$L | Conversion from Blood concentration to LN cell counts | 360000.0 | 360000.0 | u |
| Host LNs | hostLn | count | Number of involved LNs in host (Used for initial conditions) | 5.0 | 5.0 | - |
| $\lambda$ | lambda | - | Frequency of Mtb-specific naive cells in the host | 0.0001 | 0.0001 | u |
| $hs_1$ | hs1 | count | Half saturation of $APC$ in $N_4^B$ recruitment to LN | 1092.8546 | 8061.6792 | l |
| $hs_{10}$ | hs10 | count | Half saturation of $APC$ in $N_8^B$ recruitment to LN | 45.5413 | 87.7386 | u |
| $hs_{11}$ | hs11 | count | Half saturation of CD4$^+$ surrogates in $N_8^{LN}$ priming | 12.9592 | 47.9200 | u |
| $hs_{13}$ | hs13 | count | Half saturation of $APC$ in $P_8^{LN}$ proliferation | 2684.2696 | 4055.8241 | u |
| $hs_{14}$ | hs14 | count | HS of $APC$ in $P_8^{LN} \rightarrow E_8^{LN}$ and $P_8^{LN} \rightarrow CM_8^{LN}$differentiation | 1904.3871 | 4144.4838 | u |
| $hs_{17}$ | hs17 | count | Half saturation of $APC$ in $CM_8^B$ recruitment to LN | 66.1225 | 403.04107 | u |
| $hs_4$ | hs4 | count | Half saturation of $APC$ in $P_4^{LN}$ proliferation | 4057.5368 | 28401.2710 | u |
| $hs_5$ | hs5 | count | HS of $APC$ in $P_4^{LN} \rightarrow E_4^{LN}$ and $P_4^{LN} \rightarrow CM_4^{LN}$ differentiation | 4134.3783 | 8006.2583 | u |
| $hs_8$ | hs8 | count | Half saturation of $APC$ in $CM_{Non,4}^B$ recruitment to LN | 40.4710 | 57.04244 | u |
| $k_1$ | k1 | 1/day | Rate of $N_4^B$ recruitment to LN | 0.6880 | 0.8540 | u |
| $k_{10}$ | k10 | 1/day | Rate of $N_8^B$ recruitment to LN | 0.5380 | 0.6764 | u |
| $k_{11}$ | k11 | 1/day | Rate of $N_8^{LN}$ priming | 0.0001043 | 0.0002272 | u |
| $k_{12}$ | k12 | 1/day | $CM_8^{LN}$ reactivation rate | 0.0001206 | 0.0007482 | u |
| $k_{13}$ | k13 | 1/day | Rate of $P_8^{LN}$ proliferation | 0.1961 | 0.7954 | u |
| $k_{14}$ | k14 | 1/day | Rate of $P_8^{LN} \rightarrow E_8^{LN}$ differentiation | 0.2548 | 0.7351 | u |
| $k_{15}$ | k15 | 1/day | Rate of $P_8^{LN} \rightarrow CM_8^{LN}$ differentiation | 0.5286 | 0.8602 | u |
| $k_{16}$ | k16 | 1/day | Rate of $E_8^{LN} \rightarrow EM_8^{LN}$ differentiation | 0.1573 | 0.8217 | u |
| $k_{17}$ | k17 | 1/day | Rate of $CM_8^B$ recruitment to LN | 0.3483 | 0.9012 | u |
| $k_2$ | k2 | 1/day | Rate of $N_4^{LN} \rightarrow P_4^{LN}$ differentiation | 0.3279 | 0.8656 | u |
| $k_3$ | k3 | 1/day | Rate of $CM_4^{LN} \rightarrow P_4^{LN}$ differentiation | 0.02161 | 0.07979 | u |
| $k_4$ | k4 | 1/day | Rate of $P_4^{LN}$ proliferation | 2.08336 | 4.7319 | u |
| $k_5$ | k5 | 1/day | Rate of $P_4^{LN} \rightarrow E_4^{LN}$ differentiation | 0.2721 | 0.8993 | u |
| $k_6$ | k6 | 1/day | Rate of $P_4^{LN} \rightarrow CM_4^{LN}$ differentiation | 0.3110 | 0.8645 | u |
| $k_7$ | k7 | 1/day | Rate of $E_4^{LN} \rightarrow EM_4^{LN}$ differentiation | 0.6220 | 0.9249 | u |

| Parameter | non-symbolic | Units | LN/blood parameter description | Lower range | Upper range | Distribution |
|---|---|---|---|---|---|---|
| $k_8$ | k8 | 1/day | Rate of $CM_4^B$ recruitment to LN | 0.02620 | 0.06984 | u |
| $\mu_1$ | mu1 | 1/day | Death rate of $E_4^B$ | 0.2 | 0.2 | u |
| $\mu_2$ | mu2 | 1/day | Death rate of $EM_4^B$ | 0.001368 | 0.002737 | u |
| $\mu_3$ | mu3 | 1/day | Death rate of $E_8^B$ | 0.2 | 0.2 | u |
| $\mu_4$ | mu4 | 1/day | Death rate of $EM_8^B$ | 0.001368 | 0.002737 | u |
| $\mu_5$ | mu5 | 1/day | Death rate of $APC$ | 0.05 | 0.05 | u |
| $\mu_6$ | mu6 | 1/day | Death rate of $P_4^{LN}$ | 0.0005 | 0.0005 | u |
| $\mu_7$ | mu7 | 1/day | Death rate of $P_8^{LN}$ | 0.015 | 0.015 | u |
| $\mu_8$ | mu8 | 1/day | Death rate of $N_4^B$ | 0.3 | 0.3 | u |
| $\mu_9$ | mu9 | 1/day | Death rate of $\mu_9^{LN}$ | 0.05 | 0.05 | u |
| $\rho_1$ | rho1 | count | Precursor cell carrying capacity | 3467280.5534 | 14650451.8418 | u |
| $w_{P_4}$ | Wp4 | - | Contribution of $P_4^{LN}$ in $P_8^{LN}$ priming | 0.7355 | 0.7355 | u |
| $\xi_{11}$ | xi11 | 1/day | $CM_8^{LN} \to CM_8^B$ LN efflux rate | 0.2750 | 1.2233 | u |
| $\xi_{12}$ | xi12 | 1/day | $EM_8^{LN} \to EM_8^B$ LN efflux rate | 0.2190 | 1.5214 | u |
| $\xi_2$ | xi2 | 1/day | $N_4^{LN} \to N_4^B$ LN efflux rate | 1.9942 | 4.5066 | u |
| $\xi_3$ | xi3 | 1/day | $E_4^{LN} \to E_4^B$ LN efflux rate | 16.9937 | 23.6017 | u |
| $\xi_5$ | xi5 | 1/day | $CM_4^{LN} \to CM_4^B$ LN Efflux rate | 1.07179 | 3.9185 | u |
| $\xi_6$ | xi6 | 1/day | $EM_4^{LN} \to EM_4^B$ LN Efflux rate | 0.1567 | 29.2311 | u |
| $\xi_8$ | xi8 | 1/day | $N_8^{LN} \to N_8^B$ LN Efflux rate | 0.6344 | 2.2140 | u |
| $\xi_9$ | xi9 | 1/day | $E_8^{LN} \to E_8^B$ LN Efflux rate | 1.8734 | 4.4935 | u |
| $\xi_1$ | xi1 | 1/day | $N_4^B \to N_4^{LN}$ LN Influx rate | 0.3988 | 0.9013 | u |
| $\xi_4$ | xi4 | 1/day | $CM_4^B \to CM_4^{LN}$ LN Influx rate | 0.2143 | 0.7837 | u |
| $\xi_7$ | xi7 | 1/day | $N_8^B \to N_8^{LN}$ LN Influx rate | 0.1268 | 0.4428 | u |
| $\xi_{10}$ | xi10 | 1/day | $CM_8^B \to CM_8^{LN}$ LN Influx rate | 0.05500 | 0.2446 | u |
| $s_{N_4}$ | Sn4 | 1/day*$\mu L$ | Production rate of naive CD4$^+$ T-cells | 20.1207 | 47.9125 | u |
| $s_{N_8}$ | Sn8 | 1/day*$\mu L$ | Production rate of naive CD8$^+$ T-cells | 0.3830 | 0.4868 | u |
| $s_{E_{Non,4}}$ | Senc4 | 1/day*$\mu L$ | Production rate of effector non-specific CD4$^+$ T-cells | 15.3195 | 65.7082 | u |
| $s_{EM_{Non,4}}$ | Semnc4 | 1/day*$\mu L$ | Production rate of effector memory non-specific CD4$^+$ T-cells | 6.4041 | 17.04971 | u |
| $s_{E_{Non,8}}$ | Senc8 | 1/day*$\mu L$ | Production rate of effector non-specific CD8$^+$ T-cells | 10.05032 | 69.3024 | u |

| Parameter | non-symbolic | Units | LN/blood parameter description | Lower range | Upper range | Distribution |
|---|---|---|---|---|---|---|
| $s_{EM_{Non,8}}$ | Semnc8 | 1/day*$\mu L$ | Production rate of effector memory non-specific CD8$^{+}$ T-cells | 2.8856 | 5.5002 | u |

| Initial condition | Non-Symbolic | Unit | LN/Blood initial condition description | Lower range | Upper range | Distribution |
|---|---|---|---|---|---|---|
| $APC(0)$ | APC | count | Antigen-presenting cells in LN | 0 | 0 | u |
| $N_4^{LN}(0)$ | LnN4 | count | Mtb-specific naive CD4$^+$ T-cell count in LN | 2897.3857 | 6899.4095 | u |
| $P_4^{LN}(0)$ | LnP4 | count | Mtb-specific precursor CD4$^+$ T-cell count in LN | 0 | 0 | u |
| $E_4^{LN}(0)$ | LnE4 | count | Mtb-specific effector CD4$^+$ T-cell count in LN | 0 | 0 | u |
| $CM_4^{LN}(0)$ | LnCM4 | count | Mtb-specific central memory CD4$^+$ T-cell count in LN | 0 | 0 | u |
| $EM_4^{LN}(0)$ | LnEM4 | count | Mtb-specific effector memory CD4$^+$ T-cell count in LN | 0 | 0 | u |
| $N_4^{B}(0)$ | BlN4 | pg/mL | Mtb-specific naive CD4$^+$ T-cell concentration in blood | * | - | u |
| $E_4^{B}(0)$ | BlE4 | pg/mL | Mtb-specific effector CD4$^+$ T-cell concentration in blood | * | - | u |
| $CM_4^{B}(0)$ | BlCM4 | pg/mL | Mtb-specific central memory CD4$^+$ T-cell concentration in blood | * | - | u |
| $EM_4^{B}(0)$ | BlEM4 | pg/mL | Mtb-specific effector memory CD4$^+$ T-cell concentration in blood | * | - | u |
| $N_8^{LN}(0)$ | LnN8 | count | Mtb-specific naive CD8$^+$ T-cell count in LN | 5515.3085 | 7010.3557 | u |
| $P_8^{LN}(0)$ | LnP8 | count | Mtb-specific precursor CD8$^+$ T-cell count in LN | 0 | 0 | u |
| $E_8^{LN}(0)$ | LnE8 | count | Mtb-specific effector CD8$^+$ T-cell count in LN | 0 | 0 | u |
| $CM_8^{LN}(0)$ | LnCM8 | count | Mtb-specific central memory CD8$^+$ T-cell count in LN | 0 | 0 | u |
| $EM_8^{LN}(0)$ | LnEM8 | count | Mtb-specific effector memory CD8$^+$ T-cell count in LN | 0 | 0 | u |
| $N_8^{B}(0)$ | BlN8 | pg/mL | Mtb-specific naive CD8$^+$ T-cell concentration in blood | * | - | u |
| $E_8^{B}(0)$ | BlE8 | pg/mL | Mtb-specific effector CD8$^+$ T-cell concentration in blood | * | - | u |
| $CM_8^{B}(0)$ | BlCM8 | pg/mL | Mtb-specific central memory CD8$^+$ T-cell concentration in blood | * | - | u |
| $EM_8^{B}(0)$ | BlEM8 | pg/mL | Mtb-specific effector memory CD8$^+$ T-cell concentration in blood | * | - | u |
| $N_{Non,4}^{LN}(0)$ | LnN4Non | count | Nonspecific naive CD4$^+$ T-cell count in LN | 28970960.05181 | 68987195.6821 | u |
| $CM_{Non,4}^{LN}(0)$ | LnCM4Non | count | Nonspecific central memory CD4$^+$ T-cell count in LN | 13572957.9382 | 32571421.8047 | u |
| $N_{Non,4}^{B}(0)$ | BlN4Non | pg/mL | Nonspecific naive CD4$^+$ T-cell concentration in blood | 402.3744 | 958.1554 | u |
| $E_{Non,4}^{B}(0)$ | BlE4Non | pg/mL | Nonspecific effector CD4$^+$ T-cell concentration in blood | 76.5979 | 328.5413 | u |
| $CM_{Non,4}^{B}(0)$ | BlCM4Non | pg/mL | Nonspecific central memory CD4$^+$ T-cell concentration in blood | 188.5133 | 452.3808 | u |
| $EM_{Non,4}^{B}(0)$ | BlEM4Non | pg/mL | Nonspecific effector memory CD4$^+$ T-cell concentration in blood | 160.1049 | 426.2429 | u |
| $N_{Non,8}^{LN}(0)$ | LnN8Non | count | Nonspecific naive CD8$^+$ T-cell count in LN | 55147570.5476 | 70096547.5890 | u |
| $CM_{Non,8}^{LN}(0)$ | LnCM8Non | count | Nonspecific central memory CD8$^+$ T-cell count in LN | 8169880.9373 | 30928893.4342 | u |
| $N_{Non,8}^{B}(0)$ | BlN8Non | pg/mL | Nonspecific naive CD8$^+$ T-cell concentration in blood | 765.9384 | 973.5631 | u |

| **Initial condition** | **Non-Symbolic** | **Unit** | **LN/Blood initial condition description** | **Lower range** | **Upper range** | **Distribution** |
|---|---|---|---|---|---|---|
| $E^B_{Non,8}(0)$ | BlE8Non | pg/mL | Nonspecific effector CD8$^+$ T-cell concentration in blood | 50.2516 | 346.5122 | u |
| $CM^B_{Non,8}(0)$ | BlCM8Non | pg/mL | Nonspecific central memory CD8$^+$ T-cell concentration in blood | 113.4705 | 429.5679 | u |
| $EM^B_{Non,8}(0)$ | BlEM8Non | pg/mL | Nonspecific effector memory CD8$^+$ T-cell concentration in blood | 160.3165 | 305.5719 | u |

*We assume that at $t = 0$, blood and lymph node concentrations are at equilibrium, so blood concentrations are calculated via their lymph-node starting count via the conversion factor $\alpha$ and HostLNs.

6.3. **Pharmacokinetic parameter ranges.** Note that all initial conditions are set to 0, because no drug is present in the patient prior to dosing. We uniformly sampled all host-scale PK parameters and host-baseline values of granuloma-scale parameters. Finally, note that we allow for distinct values of $V_L$, $V_{PT}$, and $V_P$ within each host. This is because each drug's PK model is calibrated to a distinct datasets to replicate those experimental trajectories. This is also why we do not include PK parameters in the granuloma and host-scale sensitivity analyses, as this variation is best considered as a form of noise in the model.

| Parameter | Non-symbolic | Units | PK parameter description | Scale | Drug | Lower range | Upper range | Distribution |
|---|---|---|---|---|---|---|---|---|
| $Q_a$ | QA | $\frac{m}{h}$ | Permeability of drug from lung into viable cellular area | Granuloma | INH | 1.5 | 2 | u |
| | | | | | RIF | 1 | 1.2 | u |
| | | | | | PZA | 1 | 1.2 | u |
| | | | | | EMB | 0.05 | 0.09 | u |
| | | | | | BDQ | 0.5 | 5 | u |
| | | | | | PTM | 0.1 | 0.14 | u |
| | | | | | LZD | 0.5 | 0.9 | u |
| | | | | | MXF | 0.05 | 0.09 | u |
| $P_A$ | PA | - | Drug partition coefficient between lung and granuloma | Granuloma | INH | 0.95 | 1 | u |
| | | | | | RIF | 0.4 | 1.1 | u |
| | | | | | PZA | 0.4 | 1.1 | u |
| | | | | | EMB | 0.8 | 1.1 | u |
| | | | | | BDQ | 0.9 | 10 | u |
| | | | | | PTM | 2 | 3 | u |
| | | | | | LZD | 0.8 | 1.0 | u |
| | | | | | MXF | 0.8 | 1.1 | u |

| Parameter | Non-symbolic | Units | PK parameter description | Scale | Drug | Lower range | Upper range | Distribution |
|---|---|---|---|---|---|---|---|---|
| $Q_c$ | QC | $\frac{L}{h}$ | Permeability of drug from viable cellular area into caseum | Granuloma | INH | 0.08 | 0.1 | u |
| | | | | | RIF | 0.8 | 1 | u |
| | | | | | PZA | 0.8 | 1 | u |
| | | | | | EMB | 0.8 | 1 | u |
| | | | | | BDQ | 0.08 | 0.1 | u |
| | | | | | PTM | 0.8 | 1 | u |
| | | | | | LZD | 0.8 | 1 | u |
| | | | | | MXF | 0.08 | 0.1 | u |
| $P_C$ | PC | - | Drug partition coefficient between viable cellular area and caseum | Granuloma | INH | 0.8 | 0.8 | u |
| | | | | | RIF | 0.4 | 1 | u |
| | | | | | PZA | 0.5 | 0.8 | u |
| | | | | | EMB | 0.04 | 0.8 | u |
| | | | | | BDQ | 0.012 | 0.05 | u |
| | | | | | PTM | 1 | 1 | - |
| | | | | | LZD | 0.6 | 0.8 | u |
| | | | | | MXF | 0.2 | 0.55 | u |
| $\beta$ | beta | - | Drug penetration exponentiation base | Granuloma | INH | 0.999 | 1 | u |
| | | | | | RIF | 0.0493 | 0.0533 | u |
| | | | | | PZA | 0.999 | 1 | u |
| | | | | | EMB | 1 | 1 | - |
| | | | | | BDQ | .01 | .01 | - |
| | | | | | PTM | .0511 | .0951 | u |
| | | | | | LZD | 0.227 | 0.329 | u |
| | | | | | MXF | 0.098 | 0.172 | u |

| Parameter | Non-symbolic | Units | PK parameter description | Scale | Drug | Lower range | Upper range | Distribution |
|---|---|---|---|---|---|---|---|---|
| $\mu_C$ | muC | $\frac{1}{h \cdot kg}$ | Drug degradation and clearance or binding within caseum | Granuloma | INH | 0.8 | 25 | u |
| | | | | | RIF | 0.08 | 0.15 | u |
| | | | | | PZA | 0.08 | 0.15 | u |
| | | | | | EMB | 0.01 | 0.012 | u |
| | | | | | BDQ | 0.1 | 0.12 | u |
| | | | | | PTM | 0.32 | 0.8 | u |
| | | | | | LZD | 0.61 | 3 | u |
| | | | | | MXF | 0.01 | 0.05 | u |
| $k_d$ | kd | $\frac{1}{h}$ | Transport rate from drug transit compartments | Host | INH | 0.5 | 6 | l |
| | | | | | RIF | 0.2 | 0.4 | l |
| | | | | | PZA | 0.55 | 0.75 | l |
| | | | | | EMB | 1 | 2.75 | l |
| | | | | | BDQ | 0.03 | 0.1 | l |
| | | | | | PTM | 0.35 | 1.2 | l |
| | | | | | LZD | 0.48 | 1.3 | l |
| | | | | | MXF | 0.025 | 3.9 | l |
| - | Transit | - | Number of drug transit compartments | Host | INH | 2 | 2 | - |
| | | | | | RIF | 2 | 2 | - |
| | | | | | PZA | 1 | 1 | - |
| | | | | | EMB | 1 | 1 | - |
| | | | | | BDQ | 1 | 1 | - |
| | | | | | PTM | 2 | 2 | - |
| | | | | | LZD | 1 | 1 | - |
| | | | | | MXF | 1 | 1 | - |

| Parameter | Non-symbolic | Units | PK parameter description | Scale | Drug | Lower range | Upper range | Distribution |
|---|---|---|---|---|---|---|---|---|
| $Q$ | Q | $\frac{L}{h \cdot kg}$ | Permeability of drug into peripheral tissue | Host | INH | 0.2 | 7 | u |
| | | | | | RIF | 1.7 | 5 | u |
| | | | | | PZA | 0.1 | 0.7 | u |
| | | | | | EMB | 0.04 | 0.9 | u |
| | | | | | BDQ | 0.2 | 0.8 | u |
| | | | | | PTM | 3 | 4 | u |
| | | | | | LZD | 10.36 | 45 | u |
| | | | | | MXF | 0.05 | 40 | u |
| $\mu_D$ | muD | $\frac{L}{h \cdot kg}$ | Decay rate of drug within lung tissue | Host | INH | 0.08 | 2.5 | u |
| | | | | | RIF | 0.08 | 0.15 | u |
| | | | | | PZA | 0.08 | 0.15 | u |
| | | | | | EMB | 0.01 | 0.012 | u |
| | | | | | BDQ | 0.01 | 0.012 | u |
| | | | | | PTM | 0.32 | 0.8 | u |
| | | | | | LZD | 0.61 | 3 | u |
| | | | | | MXF | 0.01 | 0.05 | u |
| $\mu_{DP}$ | muDP | $\frac{L}{h \cdot kg}$ | Decay rate of drug within peripheral tissue | Host | INH | 0.08 | 2.5 | u |
| | | | | | RIF | 0.08 | 0.15 | u |
| | | | | | PZA | 0.01 | 0.05 | u |
| | | | | | EMB | 1 | 2.5 | u |
| | | | | | BDQ | 1.5 | 4 | u |
| | | | | | PTM | 0.2 | 0.8 | u |
| | | | | | LZD | 0.04 | 0.91 | u |
| | | | | | MXF | 0.05 | 0.6 | u |

| Parameter | Non-symbolic | Units | PK parameter description | Scale | Drug | Lower range | Upper range | Distribution |
|---|---|---|---|---|---|---|---|---|
| $V_P$ | VP | $\frac{L}{kg}$ | Volume of Plasma distribution | Host | INH | 0.5 | 3.0 | u |
| | | | | | RIF | 0.08 | 0.6 | u |
| | | | | | PZA | 0.25 | 0.75 | u |
| | | | | | EMB | 2.7 | 4.5 | u |
| | | | | | BDQ | 0.05 | 5 | u |
| | | | | | PTM | 1 | 4 | u |
| | | | | | LZD | 0.16 | 1.71 | u |
| | | | | | MXF | 0.65 | 2.5 | u |
| $P_L$ | PL | - | Drug partition coefficient between lung and plasma | Host | INH | 1 | 1 | - |
| | | | | | RIF | 0.07 | 1 | u |
| | | | | | PZA | 1 | 1 | - |
| | | | | | EMB | 12 | 16 | u |
| | | | | | BDQ | 10 | 500 | u |
| | | | | | PTM | 1.2 | 20 | u |
| | | | | | LZD | 1.1 | 2 | u |
| | | | | | MXF | 1 | 9 | u |
| $V_{PT}$ | VPT | $\frac{L}{kg}$ | Volume of peripheral tissue distribution | Host | INH | 25 | 40 | u |
| | | | | | RIF | 0.05 | 0.2 | u |
| | | | | | PZA | 0.01 | 0.05 | u |
| | | | | | EMB | 0.08 | 1 | u |
| | | | | | BDQ | 0.1 | 50 | u |
| | | | | | PTM | 3.5 | 6 | u |
| | | | | | LZD | 0.2 | 1.9 | u |
| | | | | | MXF | 0.03 | 0.4 | u |

| Parameter | Non-symbolic | Units | PK parameter description | Scale | Drug | Lower range | Upper range | Distribution |
|---|---|---|---|---|---|---|---|---|
| $V_L$ | VL | $\frac{L}{kg}$ | Volume of lung distribution | Host | INH | 0.08 | 0.08 | - |
| | | | | | RIF | 0.08 | 0.6 | u |
| | | | | | PZA | 0.08 | 0.6 | u |
| | | | | | EMB | 1 | 6 | u |
| | | | | | BDQ | 1 | 6 | u |
| | | | | | PTM | 1 | 6 | u |
| | | | | | LZD | 1 | 6 | u |
| | | | | | MXF | 0.05 | 0.6 | u |
